# RNA-mediated SOX2 temporal dynamics govern dual chromatin-engagement modes at *Nanog*

**DOI:** 10.1101/2024.09.10.612363

**Authors:** Bitong Li, Yew Yan Wong, Yoshiaki Kobayashi, Neftali Flores-Rodriguez, Tara-Lynne Davidson, Matthew S Graus, Valeriia Smialkovska, Shikha Pachauri, Susav Pradhan, Xiaohui Gao, Satoshi Uchino, Hiroaki Ohishi, Angelika Feldmann, Hiroshi Ochiai, Mathias Francois

## Abstract

Stem cell self-renewal relies on finely tuned transcriptional oscillations of pluripotency genes, yet the sequence by which transcription factors (TFs) govern “on” and “off” states remains unclear. Here, we integrate trimodal single-molecule imaging - SOX2 mobility, *Nanog* locus diffusion, and real-time *Nanog* mRNA synthesis using the STREAMING-tag reporter - to visualise endogenous regulatory dynamics in living cells. We identify two SOX2 binding modes linked to transcriptional priming and termination and show coordinated temporal activity of SOX2, OCT4, BRD4, and MED22 at the *Nanog* locus. Live-cell and fixed-cell measurements reveal that local RNA fluctuations modulate SOX2 visiting-frequency rates. During the active elongation, high nascent enhancer and mRNA levels transiently sequester SOX2 through non-specific interactions, reducing its effective repetitive rebinding and limiting interference with elongating polymerase. After transcript release, declining RNA density permits SOX2 re-engagement, with high-visiting frequency with longer residency and increased locus mobility. These dynamics reveal RNA-dependent feedback that alternately buffers and primes transcriptional re-initiation.

## Main text

The exquisite regulation of gene expression in space and time dictates cell fate decisions and lineage specification at the molecular level. Two components are essential to this process: 1) the communication between genome regulatory regions, including promoters and enhancers; 2) a molecular machinery able to coordinate the activity of the genome control panel. The latter include transcription factors (TFs) that regulate messenger RNA (mRNA) synthesis in *trans* via binding to DNA *cis*-regulatory elements.

Recent progress in imaging and single-cell analysis technologies has revealed that transcription, production of RNA molecules from a corresponding gene, occurs in a discontinuous and dynamic manner. Nascent mRNA synthesis oscillates between sustained production in an "on" state and minimal synthesis in an "off" state ^1^. The timescale of this on-and-off cycle varies between genes but typically ranges from several minutes to hours ^2^. While transcriptional activity dictates the levels of gene mRNA synthesis, the relationship between TFs dynamics and transcription burst kinetics is not yet fully understood.

TFs activity is highly dependent on chromatin region accessibility, DNA affinity and the recruitment of cell-type specific co-factors ^3–5^. In addition, multiple biophysical constraints shape TFs mobility and fine-tune their behaviour. TFs move through a complex chromatin landscape, which displays a sophisticated higher-order organisation. Importantly, TFs navigate a crowded environment while eliciting interactions with different classes of molecules, such as proteins, RNA, and DNA. Recent advances in high-resolution imaging, such as single-molecule tracking (SMT) ^6–9^, have revealed key biophysical features, such as a diffusion rate or confined mobility, that drive TF-specific activity. These quantifiable behaviours have determined TF search pattern mechanism, offering new insights into TF biology, on how a TF selects its target gene. For instance, a landmark study has demonstrated, that SOX2, which is essential for stem cell pluripotency, searches for a specific target through the combination of diffusion and sliding ^10^. Different diffusion rates of SOX2 were shown to modulate asymmetric gene expression programs during early embryo formation, limiting stem cell fates to either extra-embryonic annexes or the embryo proper ^11^. Despite a clear role for TF biophysical parameters in controlling cell fate decision, there is still a gap in our understanding of how these behavioural features directly contribute to the modulation of gene expression levels.

To unravel the mechanisms that underpin the dynamics of transcriptional activation, it is critical to be able to determine the TF/DNA binding kinetics requirement and establish a functional relationship of this characteristic with the nascent mRNA synthesis rate. Despite important findings brought with SMT-based methods, they have several limitations. Firstly, it is difficult to biologically interpret the activity kinetics of different TF molecule sub-populations (e.g. different types of confined mobility). Furthermore, the lack of sequence specificity of SMT-based techniques is not sufficient to determine the precise chromosomal coordinates of TF interaction.

To overcome these limitations, complementary methods that capture genome locations have been used to assess the activity of a protein of interest in a specific region. Some studies based on locus-centric approaches have taken advantage of anchor-based genetic reporter systems to label specific genome locations ^12^. While this approach enables the visualisation of specific DNA regions and the analysis of protein behaviour through SMT, the information about these regions, such as whether gene transcription is on or off, remains obscure. As a result, it is difficult to discriminate between how a specific molecular behaviour is functionally related to a specific step in the transcription cycle.

Investigating the relationship between TF binding kinetics and the rate of transcriptional activity is complicated by significant differences in the timescales of these two biological processes. We addressed this challenge by using a multimodal imaging approach that enables us to explore the coordinated activity of regulatory factors and nascent mRNA synthesis at a specific gene locus in live cells, as well as to measure these activities in fixed cells in a correlative manner.

In this study, we use the *Nanog* gene locus, well known to exhibit robust transcriptional bursts ^13^, as a proof-of-principle that it is possible to quantify its coordinated transcriptional activity along with its direct regulators, the SOX2 and OCT4 TFs, in real-time in pluripotent stem cells. Importantly, the SOX2/OCT4 pair has been shown to directly regulate *Nanog* transcription to maintain pluripotency with both their loss- and gain-of-function leading to impaired *Nanog* expression^14–18^.

We have achieved this by combining STREAMING-tag and SMT-based imaging techniques. While STREAMING-tag reporter system allows for profiling of locus mobility and transcription bursts at allelic resolution via labelling with different fluorescence markers ^19^, SMT provides a readout of TF behaviour at the locus of interest. This work provides a time-based picture of multifaceted SOX2 behaviours modulated by local RNA concentration and linked to distinct phases of the *Nanog* gene locus transcription cycle.

### Multi-modal microscopy to quantify SOX2 visiting frequency at the *Nanog* gene locus relative to its transcriptional on-state

To address the main limitation of converting biophysical observations into a relevant biological interpretation in the context of TF mobility and gene transcription, we opted for a paired measurement that combines two live imaging methods: SMT and the STREAMING-tag reporter system. We used a multimodal imaging platform to measure TF long dwell times on the chromatin (e.g. over 4 sec or more) also known as confined mobility in parallel with processive transcriptional stages. The assessment of STREAMING-tag reporter activity via Spinning Disk Confocal (SDC) module allows us to measure nascent mRNA synthesis and locus mobility, while the slow tracking SMT using the HILO module displays TF chromatin binding dynamics at a single locus resolution (see Methods). This readout gives access to TF behaviours at a gene-specific locus relative to mRNA nascent transcription and allows quantifying TF binding dynamics and dwell time relative to the “on” and “off” states. This approach was established in mouse embryonic stem cells (mESCs) by introduction of the STREAMING-tag reporter cassette (MS2 and TetO repeats) at the *Nanog* locus with expression of reporter proteins (MCP-RFP and mTetR-GFP), and the SNAP-tag knocked-in into either the *Sox2, Oct4, Med22* or *Brd4* locus ^19^ (Figure 1A).

**Figure 1.**
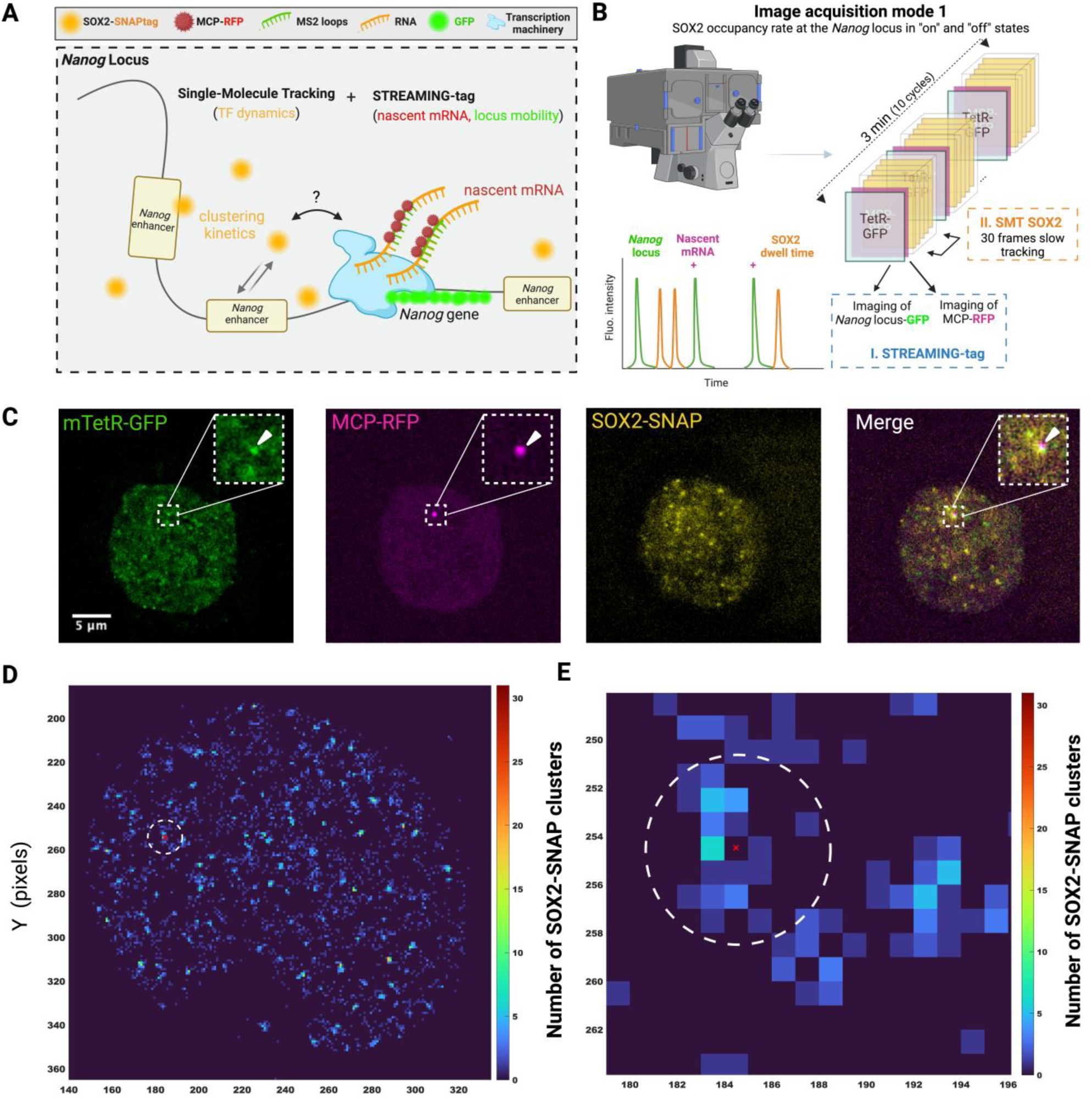
Multimodal imaging method to visualise transcription factor activity at a single locus resolution relative to its transcriptional state. (**A**) Schematic of the principle for dual imaging to capture SOX2 clustering dynamics at the *Nanog* locus relative to the event of nascent mRNA synthesis. The STREAMING-tag cassette at the *Nanog* locus enables the detection of the *Nanog* mRNA synthesis (MS2/MCP-RFP) along with the *Nanog* locus mobility (TetO/mTetR-GFP) while SNAP-tag knocked-in at the *Sox2* locus reveals chromatin binding dynamics rate of this TF via single molecule tracking (SMT). (**B**) Principle of the imaging acquisition mode 1 set up to capture the fluorescent signals from STREAMING-tag and SMT detection methods in a live cell. Multimodal image acquisition uses a combination of SDC (STREAMING-tag) and HILO (SMT) microscopy (**C**) Fluorescent signals from the *Nanog* locus (mTetR-GFP), *Nanog* mRNA (MCP-RFP) and SOX2-SNAPtag. (**D**) Representative heat map of most visited locations accumulated over a 3 min period for the detected SOX2 molecules across the cell nucleus. The heat map reveals an overlap between SOX2 long dwell time and the region of interest for the *Nanog* locus (red cross). (**E**) Higher magnification of the dashed circle shown in panel D demonstrates SOX2 clustering activity within a 4-pixel (440 nm) radius ROI (circle dashed line) centred on the *Nanog* locus when both GFP (red cross) and RFP signals are detected.

Previous studies on fixed cells using super resolution-based approaches have shown that around 10 to 20 molecules of regulatory factors cluster on enhancers scaffolds that come into proximity of their target gene locus ^20,21^. Although our imaging detects only the fluorescently labelled fraction, under our sparse-labelling conditions and using dark-state / stoichiometry calibrations similar to those described in refs ^20,21^, we estimate that each fluorescent spot typically corresponds to a small assembly of multiple SOX2 molecules rather than a true single molecule. We therefore use the term spatial clustering to describe these nanoscale protein assemblies. In addition, our live-cell approach reveals a second, conceptually distinct phenomenon, which we refer to as temporal clustering - bursts of high-frequency revisiting of the locus by these assemblies over time. Both spatial and temporal clustering are central features of SOX2 behaviour and are treated separately in our analyses.

To further validate the dynamics for this type of SOX2 behaviour in our reporter system, we performed a simultaneous two-colour SMT experiment with sparse labelling (JF549) in one channel and saturated labelling (JF646) in another channel in the same cells (Figures S1A-C). Fluctuations in fluorescent signal intensity under saturated labelling conditions were used to construct masks to define regions of different SOX2 concentration, with high as the top 8%, following 30% as medium and the remaining signal as low (see Methods). The sparse SOX2 labelling signal was overlaid onto the mask to assess the dark fraction contribution to individual SMT-detected molecules (Figures S1B and C). This approach further confirms work on fixed cells and suggests that on average over 50% of the SMT-detected SOX2 molecules may accumulate in domains of high or medium intensity. This indicates that most imaged single SOX2 molecules are involved with foci made of at least two or more SOX2 molecules (Figure S1C and Video S1).

Because TF chromatin binding dynamics occur on a shorter time scale (seconds) compared to transcription bursts (minutes), we chose a two-step measurement method that continuously monitors SMT interspersed with a regular assessment of transcriptional activity at the *Nanog* locus. To accomplish this, we utilised a dual image acquisition technique that discriminates between SOX2 chromatin binding dynamics and *Nanog* mRNA elongation kinetics (3 minutes) (Figure 1B, image acquisition mode 1). This imaging approach relies on continuously measuring by SMT for 30 frames (exposure time 500 ms, no interval time), alternating with 1 frame that records both GFP and RFP signals over a 3-minute period. Fluorescent beads were used to evaluate the changes in light path occurring between shifts from SDC to HILO modalities. Post-image acquisition, we applied a custom script to correct for image misalignment (see Methods) (Figures S2A-F). The STREAMING-tag system has been designed to denote transcription at a gene locus. This is done through tagging a specific gene locus with a GFP signal and actively transcribed mRNA with RFP. With this reporter system, the presence of a GFP-only signal denotes that the transcription of the *Nanog* locus is "off". Co-localisation of the GFP and RFP spots within a 440 nm region of interest (ROI, 4-pixel radius) determines that nascent *Nanog* transcription is occurring, defining an “on” state. Of importance, the STREAMING-tag cassette is knocked at around 300 bp downstream from the transcription start site (TSS), while transcriptional pausing has been shown to occur about 50-60 bp downstream of the TSS ^22–24^. Therefore, with the STREAMING-tag reporter system, only the productive elongation after pause-release is observed (Figure S2G). This means that the “off” state, as we define it in this study, also includes the early steps of transcriptional initiation up to the release stage. The “on” state enables the detection of nascent mRNA produced during transcriptional elongation. When the RFP signal is lost, this means that splicing has occurred, or termination has been completed.

To measure the SOX2 visiting frequency in a locus-specific manner while *Nanog* is in an “on” state (Figure 1C), we used an ROI of 4-pixel radius, (440 nm) that centres on the GFP spot when both GFP and RFP signals are detected (Figures 1D and E). Such a distance was defined based on the previous observation that elevated concentration of regulatory factors around the *Nanog* locus ranges between 200 and 400 nm ^19,21^. Unsurprisingly, we find multiple locations with SOX2 enrichments all throughout the nucleus (Figure 1D, heatmap). Interestingly, within 440 nm of the GFP detection, we systematically observe an enrichment of SOX2 events (Figure 1E, ROI dashed circle), suggesting that clustering activity is temporally correlated with the *Nanog* locus activity.

### Distinct types of SOX2 chromatin binding dynamics at the *Nanog* locus during on and off-state

Here, we measured the SOX2 visiting frequency at the *Nanog* locus to determine its temporal clustering profile while the transcription is “on”. It revealed distinct modes of visiting frequency that occur at different rates throughout transcription elongation (Figures 2A and B and Video S2). The first mode indicates no to low clustering (Figure 2A), while the second shows persistent locus visiting frequency of moderate intensity (Figure 2B).

**Figure 2.**
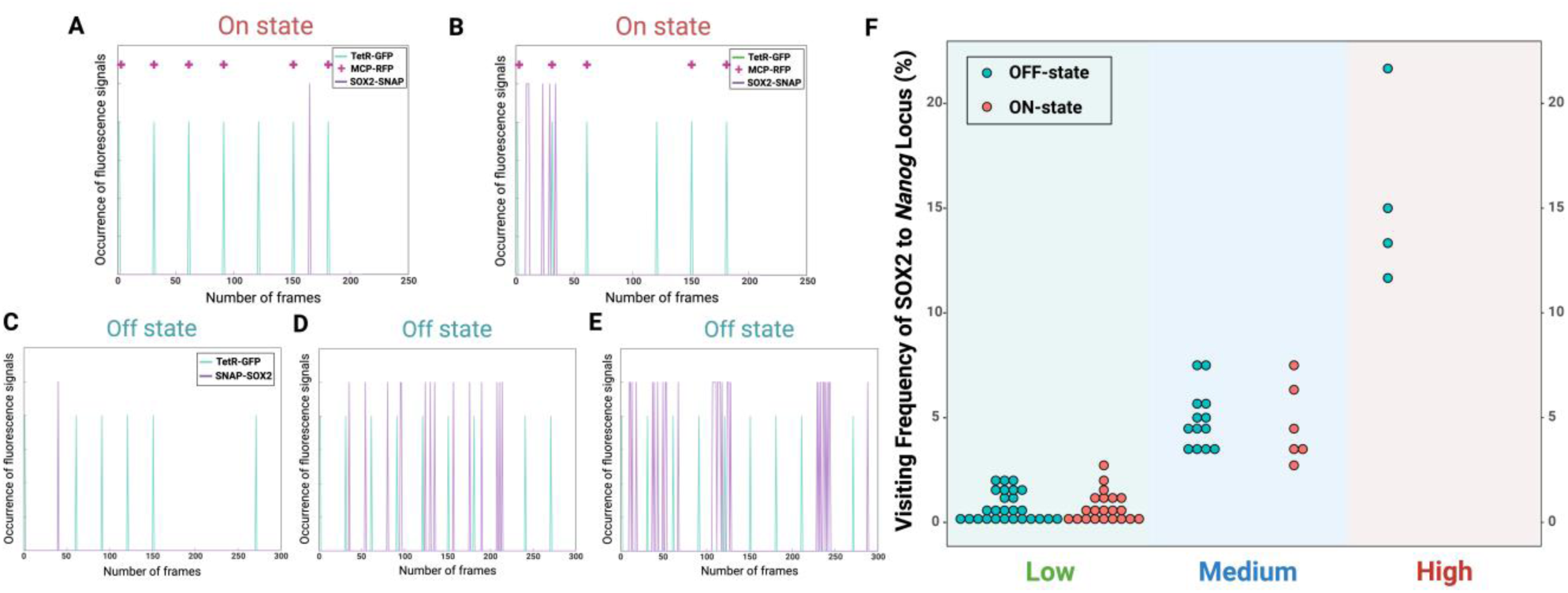
Multiple SOX2 clustering dynamics behaviours at the *Nanog* locus during “off” state. (**A-B**) On-state. Occurrence of fluorescence signals for GFP (green) for the *Nanog* locus, SOX2 SMT signal (purple) and RFP (magenta +) for *Nanog* transcriptional activity (on-state). **(C-E)** Off-state. Occurrence of fluorescence signals for GFP (green) for the *Nanog* locus, SOX2 SMT signal (purple) showing the visiting frequency of SOX2 when the locus is available, but production of nascent mRNA is off. **(F)** Quantitation of the different SOX2 behaviours during the on-state (n = 27) and off-state (n = 43) classified based on the visiting frequency (the total number of SOX2 molecules that have visited this GFP positive spot normalised to the total number of frames). The visiting frequency accounts for the number of SOX2 molecules that have visited the GFP spot.

It is known that the SOX2 protein binds to enhancers at the *Nanog* locus to induce transcriptional transactivation ^25,26^; however, the timing of this event in relation to transcription cycle remains elusive. Hence, to understand how SOX2 dynamics correlates with the timing of transcription burst, we next set out to evaluate the visiting frequency of this TF during the “off” state period.

The assessment of SOX2 visiting frequency while the locus is not actively transcribing RNA uncovers a third type of molecular behaviour characterised by a high rate of visiting events over a short number of frames (e.g SOX2 detected at least in 30 frames out 300, high rate ≥ 10% or more of frame coverage) (Figures 2C-E and Videos S3-5, see method for the numerical definition of all visiting frequency rates). Quantitation of the different types of temporal clustering kinetics (Figure 2F) across the cell population revealed that low to medium visiting frequency are the most prevalent types of visiting frequency at both on and off state, and that a discrete population of cells (5.7%) display SOX2 molecules with high clustering activity only at off state. These different clustering activities indirectly reflect possible varying oligomeric status and recruitment of specific protein partners as well as random interaction with RNA products, all likely to influence this TF high-frequency rates ^20,27–30^. To show that the different SOX2 visiting frequencies are not restricted to the *Nanog* locus only, we took advantage of the dual SMT dataset described above to analyse the presence of different types of visiting rate relative to the fluctuation of SOX2 local concentration across the nucleus. This analysis has shown that all three types of clustering activity (low, medium and high) are broadly detected across the nucleus in regions that show elevated SOX2 concentrations (Figure S3A). Quantitation of these three types of visiting frequency revealed that, the occurrence of high SOX2 clustering is a broad mechanism used on a genome-wide scale most likely across many gene loci (Figure S3B).

Chromatin organisation and epigenetic modifications both play important roles in the activation of transcriptional initiation and are modulated in a cyclical, stepwise fashion. These periodic series of ordered and coordinated molecular events have been included in the theoretical modelling of transcription ^31–34^. Further, a mathematical model proposed a molecular framework for the transition between “off” and “on” states that considers transcriptional activator residence time relative to the burst frequency ^34^. Such a conceptual framework prompted us to assess SOX2 chromatin binding dynamics in relation to the progression through a full cycle of transcription.

### Stable SOX2 clusters are anti-correlated with its concentration and occur post *Nanog* mRNA elongation

Based on the population average of SOX2 long dwell time in mESC (4-5 sec, Figure S4) ^11,35^, and the average time of transcription burst at the *Nanog* locus (3 min) ^13^, we designed a second mode of acquisition to image continuously through a full transcription cycle (Figure 3A, image acquisition mode 2). Briefly, this approach alternates confocal imaging of the *Nanog* locus for 3 seconds and SMT measurement for 500 ms with a repeat of this cycle over 10 minutes (170 cycles total, see Methods, Video S6). Such a design allowed us to infer that a detected SOX2 molecule (long-dwell time ≥ 4 sec) prior to and after the *Nanog* locus imaging is the same molecule. To be conservative, in addition to the detection of long-lived fraction (2-frame detection, Figure S5A, schematic), we classified SOX2 short dwell time events that corresponded to the capture solely over one frame period (Figure S5B, schematic). This approach enables us to measure the presence of SOX2 clusters that persist near the *Nanog* locus over time.

**Figure 3.**
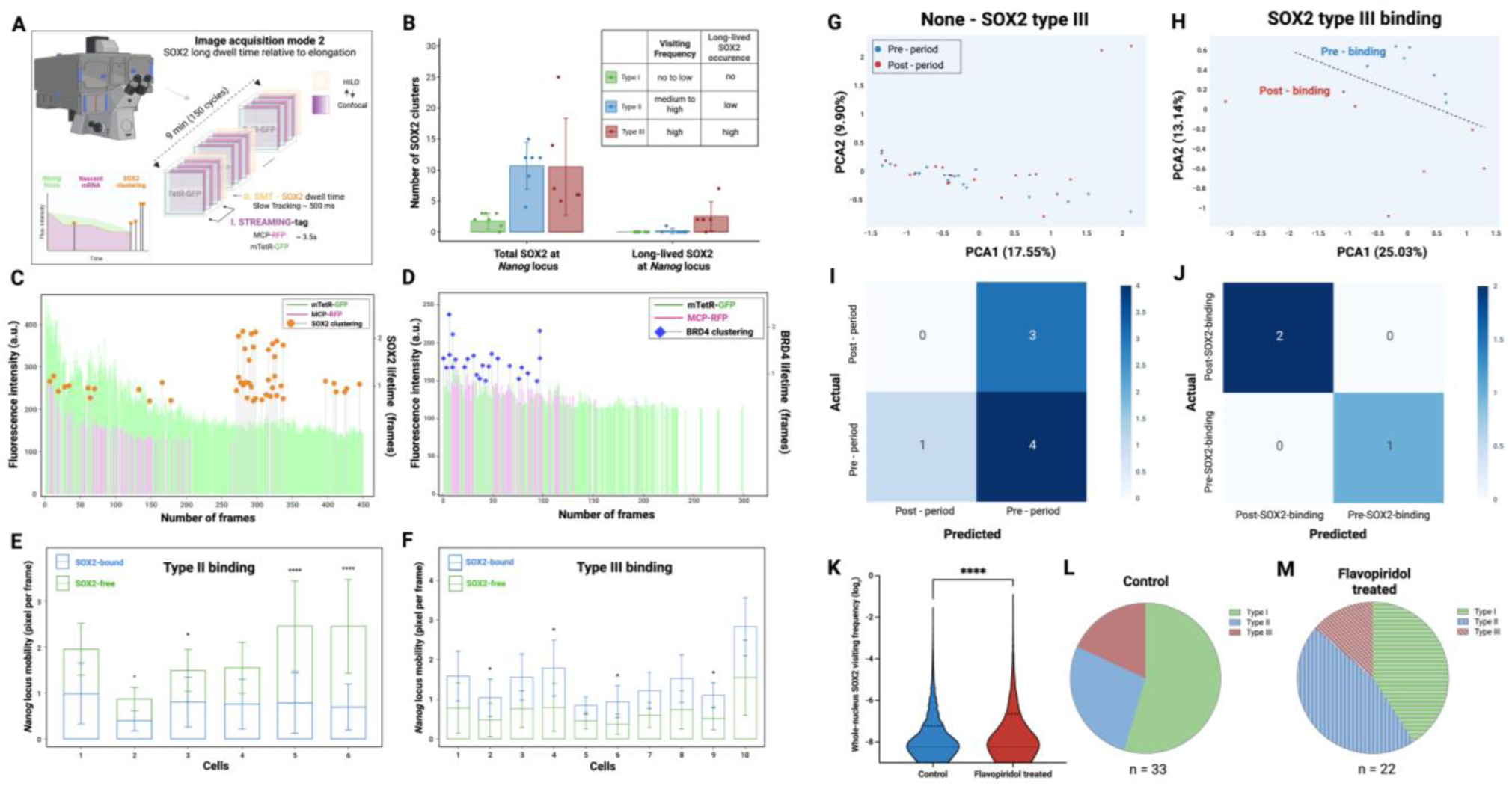
Distinct types of SOX2 burst dynamics associate with different locus mobility and frame the elongation process at the *Nanog* locus. **(A)** Image acquisition mode 2 set up to capture a full transcription cycle at the *Nanog* locus (STREAMING-tag) in parallel to SOX2 or BRD4 chromatin binding dynamics (SMT). **(B)** Quantification of detected SOX2 clusters and their relative survival rate at the *Nanog* locus. (**C** and **D**) SOX2 **(C)** or BRD4 **(D)** dwell time relative to the elongation process. Fluorescent intensity at the *Nanog* locus is shown relative to time. *Nanog* locus signal is labelled with GFP (green), and *Nanog* mRNA synthesis (magenta). SOX2 (orange) and BRD4 (blue) long dwell times (over 2 frames) and short dwell times (over 1 frame) reveal diverse binding kinetics relative to the elongation process. (**E** and **F**) 2D *Nanog* locus mobility in the presence (blue) or absence (green) of SOX2 binding. **(G-H)** Principal component analysis (PCA) of nuclear features extracted through temporal MIEL analysis. **(G)** PCA of nuclear features from cells without SOX2 Type III binding. For these cells, two frame segments were selected at random but matched in duration to the pre- and post-binding windows used for Type III cells and are labelled “pre-period” and “post-period” for comparability. These two segments show no clear separation along the principal components, indicating minimal change in nuclear morphology or texture over time in the absence of SOX2 Type III binding events. **(H)** PCA of frame segments (variable in exact frame numbers across cells) corresponding to the pre-binding and post-binding periods of SOX2 Type III events. Clear separation along the principal components indicates distinct multivariate nuclear morphology and texture associated with the pre- versus post-SOX2-binding states. **(I-J)** Confusion matrix illustrating the performance of nuclear-feature-based classification of SOX2 binding state, shown separately for **(I)** cells without SOX2 Type III events and **(J)** cells with SOX2 Type III binding. Rows represent the actual class labels, and columns represent the predicted labels. High diagonal values in **(J)** indicate correct classifications and strong separability between pre-SOX2-binding versus post-SOX2-binding nuclear states, whereas off-diagonal values in **(I)** reflect misclassification and weaker distinction between two states. **(K)** Violin plot showing the distribution of log₂-transformed whole-nucleus SOX2 visiting frequency in control and flavopiridol-treated cells. Individual data points are overlaid, with the median (thick dashed line) and interquartile range (thin dashed lines) indicated. Flavopiridol treatment shifts the overall distribution toward higher visiting frequencies compared with control. **(L-M)** Pie charts showing the distribution of SOX2 binding behaviours (Type I, Type II, and Type III) in the transcriptionally “off” state for **(L)** control cells (n = 33) and **(M)** flavopiridol-treated cells (n = 22). Flavopiridol treatment alters the proportion of binding modes, with a relative increase in Type II compared with control.

To fully cover a complete event of transcription burst, we analysed SOX2 lifetime during a period that encompasses a step of transcriptional elongation and its transition to a refractory period, until the nascent RNA signal disappeared (Figure 3A). Analysis of SOX2 visiting frequencies with acquisition mode 2 over a 10-minute period while the *Nanog* locus is in off-state further confirmed the presence of the three types of interaction lifetime (low, medium and high) reported above with mode 1 imaging. The analysis of the number of SOX2 detected molecules and their relative lifetime at the *Nanog* locus revealed that, in addition to different types of visiting frequencies, SOX2 clusters display both short-lived and long-lived fractions (Figure 3B). From there it is possible to classify SOX2 visiting frequency into three distinct types: type I - low to no visiting rates and short lived; type II - medium to high visiting rates and short residency time; and type III - high visiting frequency and long lived (Figures 3B and C and Figure S6). Importantly, type III behaviour is mostly found in off state, whereas type II and type I are found in both on and off states (Figure 3C).

Tri-modal confocal bulk imaging of the locus to assess relative SOX2 concentration revealed that this TF levels are the most elevated while *Nanog* is in on-state whereas residual but above background protein levels are present at off-states (Figure S7). Altogether these results suggest that SOX2 stable chromatin engagement is anti-correlated to its concentration and mostly take place post elongation when this TF is at low density.

SOX2 is well known to act in concert with OCT4 to regulate *Nanog* transcription ^36,37^. To gain a comprehensive view of this regulatory pair, we quantified OCT4 dynamics at the *Nanog* locus using a transgenic ES cell line carrying an endogenous OCT4-SNAP tag along with a *Nanog* STREAMING-tag reporter cassette. Both modes of image acquisition were performed to quantify OCT4 behaviours. Akin to SOX2, OCT4 exhibited low-to-moderate visiting frequency during transcriptional on-states (Figure S8A). However, unlike SOX2, OCT4 stable cluster formation was rarely detected during transcriptional off-states (Figure S8A) post elongation (Figure S8B). To firmly demonstrate this negative observation, we performed a comparative genome-wide analysis of SOX2 and OCT4 clustering activity. This approach showed that although OCT4 can engage in high interaction lifetime with high visiting frequency, the rate of this molecular behaviour is significantly lower than the one of SOX2 (Figure S8C).

To test whether high-visiting frequency is a feature of other SOX family members, we also examined SOX18, an endothelial-specific TF from the F-group. Interestingly, SOX18 showed high visiting rates across the nucleus (Figure S8D) at a similar rate than SOX2, indicating that pronounced clustering dynamics are not unique to SOX2 and may represent a broader property of SOX transcription factors whose functional basis remains to be defined.

To further understand the relationship between the temporality of SOX2/OCT4 residency time and mRNA synthesis, we next imaged the activity of a well-known elongation factor, the BRD4 protein (Figure 3D) as well as a component of the mediator complex MED22 (Figure S9). Here, we used a reporter cell line that harbours the *Nanog* STREAMING-tag reporter system combined with a SNAP-tag that has been knocked into the *Brd4* or *Med22* locus ^19^and employed the imaging acquisition mode 2. This approach revealed that BRD4 interaction lifetime is mostly concurrent with nascent mRNA production (Figure 3D and Figure S10) ^19^ whereas MED22 is observed either early or late in the transcription cycle (Figure S9). Importantly, this approach further confirmed that the long-lived and highly active SOX2 chromatin engagement (type III) is out-of-phase with the elongation step that involves BRD4, and RNA polymerase II activity (Figures 3C and S11).

Since earlier studies showed that open chromatin regions drift relatively slowly compared to closed ones that diffuse faster ^19,38^, this prompted us to investigate further how SOX2 residency time correlates with the *Nanog* locus mobility.

It has been shown that transcription constrains chromatin motion ^39–41^, hence, to couple the different types of SOX2 molecular behaviours with an active or inactive transcriptional state, we used as a proxy change in *Nanog* locus mobility in relation to the presence or absence of SOX2 type II/III binding dynamics. This quantitative approach revealed that the *Nanog* locus displays opposite diffusion kinetics that are associated to different types of SOX2 residency profiles. During SOX2 type II binding event, the *Nanog* locus harbours a slower diffusion rate while SOX2 is present compared to when there is no SOX2 detected (Figure 3E). Conversely, the locus diffusion kinetics increases in the presence of high SOX2 visiting frequency (type III) compared to when there is no SOX2 detected (Figure 3F). This observation was further validated by quantifying changes in chromatin density at the *Nanog* locus before and after SOX2 binding while the TF displays type III chromatin binding dynamics (Figures 3G-J). Using a modified MIEL-based analysis (threshold adjacency statistics ^42,43^) on NucBlue (Hoechst33342) labelling combined with trimodal imaging (Figure S12, schematic), we observed that chromatin texture differs significantly when comparing pre- and post-SOX2 binding states (Figures 3H and J). In contrast, analysing chromatin density fluctuation at the locus in the absence of SOX2 type III binding events did not reveal any significant changes (Figures 3G and I). Recent, work from Platania and colleagues has shown that locus diffusion is a reliable predictor of enhancer to promoter contact ^44^, therefore, by combining information gained on change in locus mobility and density with residency time and visiting frequencies measurement, we posit that SOX2 type II binding profile (high visiting frequency rates/short dwell time) is associated with open chromatin state and occurs prior to active nascent mRNA synthesis as a priming mechanism. This is opposed to type III (high visiting frequency rates/long dwell time) that is associated to a more compact chromatin state or possibly RNA-POLII eviction, likely to follow the elongation step and is related to elongation termination to reset the locus. To tease apart the involvement of SOX2 across these two different mechanisms, we next set out a series of pharmacological perturbations.

### Local RNA fluctuation but not chromatin availability influence SOX2 chromatin engagement

We imaged SOX2 mobility in presence of either a NuRD small molecule inhibitor (BPK-25), or TSA, or Flavopiridol (CDK 9 inhibitor). NuRD is a chromatin remodelling complex involved in gene repression, nucleosome repositioning and stem cell pluripotency ^45^. We performed both genome-wide and single-locus imaging of SOX2 at the *Nanog* locus and quantified TF visiting frequency. We compared control conditions with treatment using the NuRD inhibitor BPK25, the HDAC inhibitor Trichostatin A (TSA; increases chromatin accessibility through deacetylation inhibition, but via a mechanism distinct from BPK25), and the CDK9 inhibitor flavopiridol (which prevents the transition from promoter-proximal pausing to productive elongation). Neither BPK25 nor TSA altered the SOX2 high-frequency visiting rate, indicating that relieving chromatin compaction - whether by increasing acetylation or inhibiting NuRD - does not promote repetitive SOX2 rebinding (Figures S13A and B). In contrast, short pulse of flavopiridol treatment (3 hours) significantly increased the SOX2 high-frequency visiting rate on a genome-wide scale and at the *Nanog* locus (Figures 3K and L), suggesting that low residual mRNA concentration is insufficient to limit SOX2 access to chromatin. These findings support a model in which local fluctuations in nascent mRNA/Enhancer RNA (eRNA) production (and associated transcriptional activity) are a key determinant of SOX2 revisiting dynamics, whereas simply increasing the abundance of accessible chromatin template does not drive the same SOX2 molecule to engage with chromatin more actively.

Live imaging observations showing that SOX2 high-frequency revisiting is suppressed during active elongation but enhanced when transition from promoter-proximal pausing to productive elongation is blocked suggests a biphasic regulatory role for RNA. In addition, SOX2 has been shown to recruit RNA ^28,29^ and more broadly, TFs have been suggested to harbour an RNA-binding arm, which also influences their mobility ^46^. Altogether, these lines of evidence prompted us to examine whether direct SOX2–*Nanog* mRNA/eRNAs interactions occur.

eRNAs function involves stabilising chromatin loops and recruiting co-factor such as p300 and acting as a sponge for the negative elongation factor (NELF) ^47^. Since their role in modulating transcription is a possible mechanism, this prompted us to specifically investigate the role of *Nanog* eRNAs involvement in the transcription cycle. The *Nanog* locus is one of the most well-characterised genomic regions in the context of embryonic stem cell pluripotency, with the presence of three super-enhancers that frame the gene body within a 100 kb region ^48,49^. These regulatory regions are located at -5 kb, -45 kb, and +60 kb from the transcription start site (TSS), respectively ^48^ and are actively transcribed ^50^. The functional hierarchy of these three enhancer regions in mESCs has been validated using genome editing with -5 kb being the most critical one, -45 kb of modest influence and +60 kb with no impact on the expression ^49^.

Taking advantage of the SNAP-tag as a molecular handle to pulldown SOX2, we performed targeted PCR profiling of associated eRNAs/mRNAs and validated these interactions by sequencing (Figures 4A-D and S14A-B). This approach revealed that all three *Nanog* eRNAs from both sense and antisense strands, as well as first-exon transcripts, form detectable complexes with SOX2. These results highlight a close relationship between the local RNA micro-environment at the *Nanog* locus and SOX2 interaction lifetime.

**Figure 4.**
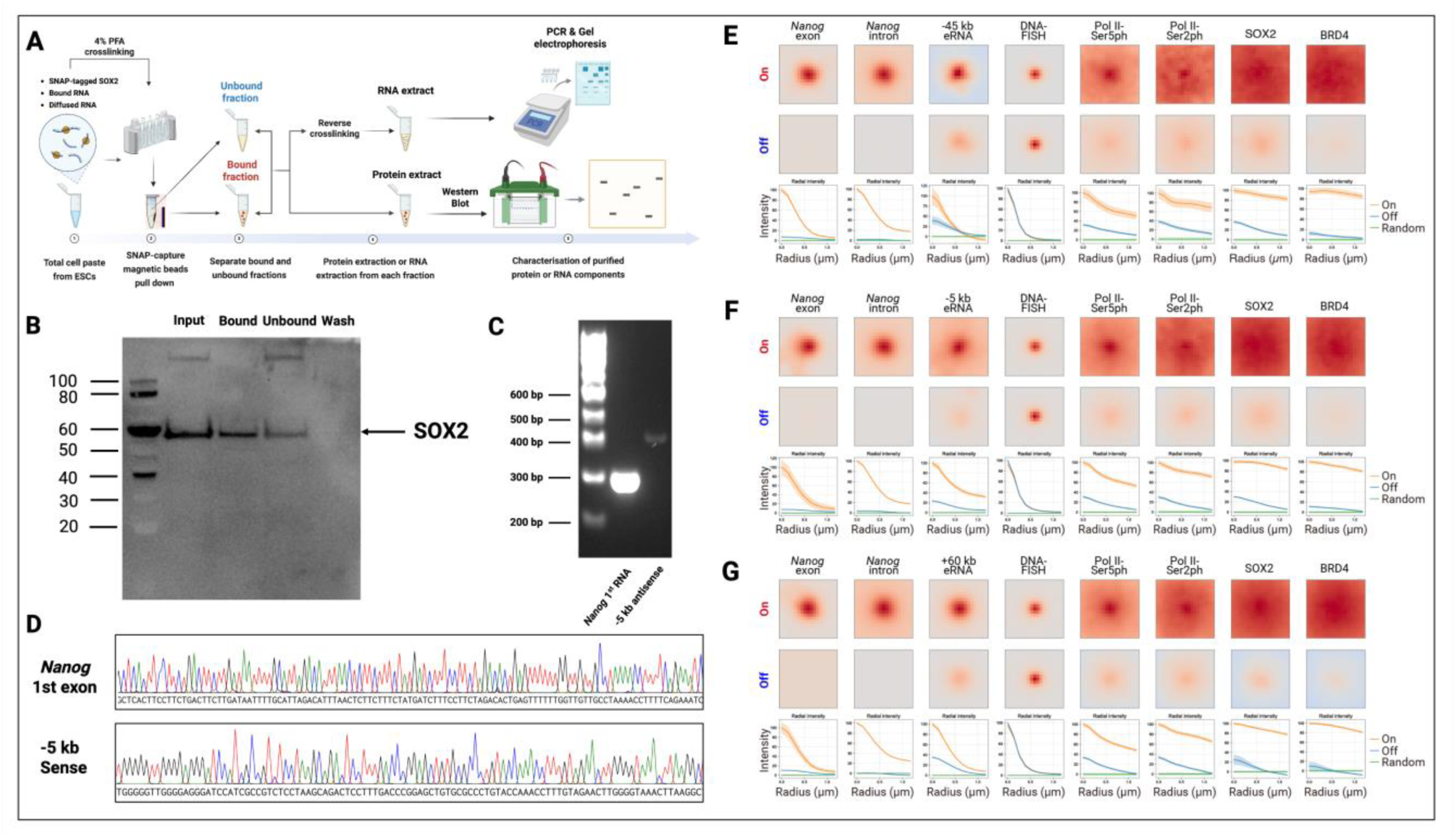
SOX2 and eRNA accumulation at the *Nanog* locus and evidence of their interaction. **(A)** Schematic of the crosslinking, pulldown, and detection workflow used to detect *Nanog* RNA and eRNA molecules associated with SNAP-tagged SOX2. Cells were crosslinked with PFA, nuclei were isolated, and RNA-protein complexes were captured using SNAP-Capture magnetic beads. Crosslinks were reversed, and the released RNA was extracted and subjected to PCR followed by Sanger sequencing. **(B)** Western blot confirming successful pulldown of SNAP-tagged SOX2. **(C)** Agarose gel electrophoresis showing specific PCR amplification of *Nanog* first-exon RNA and the - 5 kb eRNA fragment from SOX2 pulldown samples. **(D)** Sanger sequencing of PCR products validating the identity of the amplified *Nanog* first exon and -5 kb enhancer RNA fragments. Results for additional eRNAs are shown in Figures S14A and B. **(E–G)** Visualisation of RNAs, eRNAs, and protein or post-translational modifications around the *Nanog* locus using multiplexed DNA/RNA FISH combined with sequential immunofluorescence. *Nanog* exon and intron transcripts were detected by RNA-FISH; the −45 kb **(E)**, −5 kb **(F)**, and +60 kb **(G)** eRNAs were visualised using HCR-FISH; the surrounding genomic region was labelled by DNA-FISH; and Pol II-Ser5ph, Pol II-Ser2ph, SOX2, and BRD4 were detected by immunofluorescence. Experimental details are provided in Materials and Methods. Sample sizes for on and off states were as follows: **(E)** On: n = 622, Off: n = 6821; **(F)** On: N = 1386, Off: n = 11,642; **(G)** On: n = 2950, Off: n = 9270.

To contextualise variations of SOX2 concentrations relative to RNAs and regulatory factors, next we multiplexed DNA/RNA FISH with sequential immunofluorescence to quantify the local concentrations of SOX2, BRD4, active RNA Pol II, and *Nanog* eRNAs at the *Nanog* locus. Taking advantage of GRO-seq datasets ^48^, we designed probe sets that detect both sense and/or anti-sense eRNA strands for each of the three *Nanog* enhancers.

### Concentrations of regulatory factors and RNAs peak during active elongation

For each molecular effector, fluorescent signal at the *Nanog* locus was averaged over thousands of cells and the local concentration was assessed based on fluorescent intensity around the gene locus (Figures 4E-G). These measurements showed that during transcriptional on-states, regulatory factors and eRNAs reach their maximum concentration (Figures 4E-G), which correlate with SOX2 short residency time observed by SMT experiment. By contrast, off-states are characterised by lower factor abundance, still significantly above background that are correlated to increase in high SOX2 visiting frequency and longer residency time as measured by real time imaging.

Given that SOX2 physically associates with nascent RNA at the locus, our findings are fully compatible with RNA-mediated modulation of SOX2 chromatin binding dynamics, and they do not mandate a single mechanistic direction. Instead, fluctuations in nascent RNA abundance may influence SOX2 in multiple, potentially co-existing ways: at high RNA concentration, non-specific interactions could help retain or locally concentrate SOX2 molecules in the vicinity of the locus, whereas the same RNA pool could also transiently compete with chromatin for SOX2 binding. Conversely, when RNA levels are low, reduced competition might favour chromatin-bound SOX2.

To refine our understanding of the *Nanog* locus transcription burst rates at enhancers and promoter, we next set out a quantitation of relative mRNA and eRNAs concentration throughout the transcription cycle.

### *Nanog* eRNA and gene body transcription are independent

Using enhancer transcription as a proxy for TF activity at these regulatory regions, we interrogated the presence of eRNAs molecules in relation to *Nanog* nascent transcripts. We devised an approach for quantifying eRNA/nascent mRNA expression in a high-throughput manner over thousands of cells using single molecule RNA FISH (Figures 5A and S15). This analysis revealed that the expression of at least one or more sense or anti-sense eRNAs occurs in more than 80% of the stem cell population (Figure 5B). Co-expression of all three eRNAs was observed in approximately 15-20% of the cells (Figure 5B), with the +60 kb region being by far the most highly transcribed enhancer on the anti-sense strand (70%) as opposed to 30-35% for the -45 kb and -5 kb (Figure 5B). Pairwise analysis of each combination of eRNAs (sense and anti-sense) revealed that at least 30% of the stem cell population co-express an eRNA pair, with a set of sense eRNAs (+60 kb/-5 kb or +60 kb/-45 kb) being co-expressed in around 15% of the cells. Due to eRNAs rapid degradation rate, these co-expression events suggest that enhancers were probably transcribed almost simultaneously.

**Figure 5.**
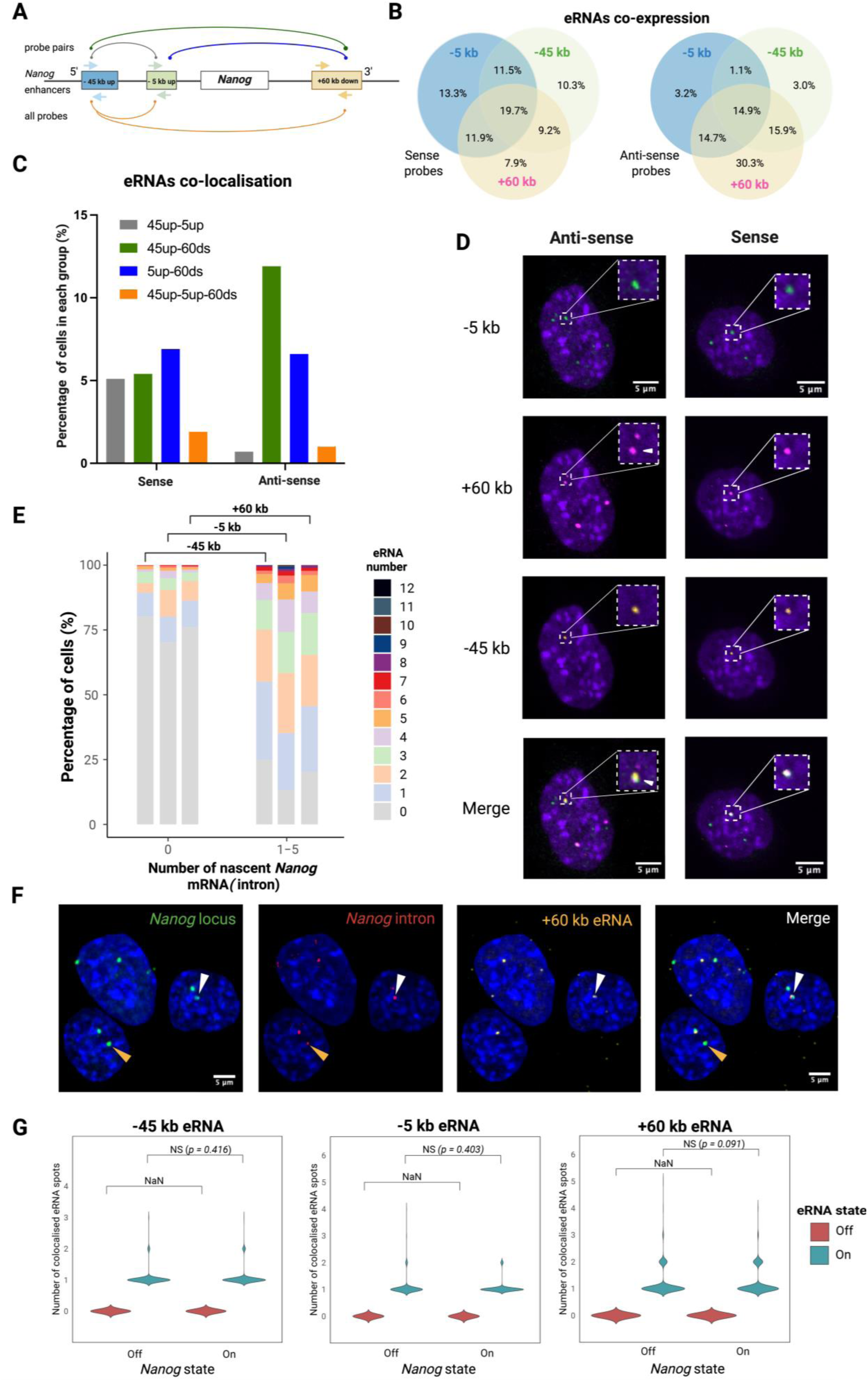
*Nanog* enhancers are individually or synergistically transcribed, but independently from *Nanog* gene body transcription. (**A**) Map of eRNAs probes (coloured arrows indicate sense and anti-sense orientations) at three regulatory regions involved with *Nanog* gene transcription. Distances from transcription start sites are -45 kb, -5 kb and +60 kb. Coloured lines connecting the different enhancers represent the different probe combinations used to assess co-localisation: grey [-45 kb/-5 kb]; green [-45 kb/+60 kb]; blue [-5 kb/+60 kb]; orange, all three probes. (**B**) Venn diagram showing the co-expression levels of different eRNAs in embryonic stem cells. **(C)** Quantitation of three eRNAs co-localisation either assessed by pairs or in sets of three for both sense and anti-sense transcripts (≥ 4000 cells imaged per condition). **(D)** Single-molecule FISH showing sub-nuclear co-localisation of *Nanog* eRNAs for both sense and anti-sense products (green -5 kb, magenta +60 kb, - 45 kb orange), scale bar 5μm. **(E)** Abundance of three different *Nanog* eRNAs correlated to the *Nanog* intron expression. **(F)** Single-molecule RNA-DNA FISH showing within a 4-pxiel (440 nm) radius, nuclear co-localisation of the *Nanog* locus (green), *Nanog* introns (red) and *Nanog* eRNAs (orange, +60 kb eRNA), scale bar 5μm. White arrowheads indicate a co-localisation event for the *Nanog* locus, *Nanog* introns and *Nanog* eRNAs, representing the *Nanog* gene body and its enhancer in a transcriptional on-state. Orange arrowheads indicate co-localisation of the *Nanog* locus and *Nanog* introns only, representing a state where the *Nanog* gene body is transcriptionally on, whereas the enhancer is off. **(G)** Quantitative analysis of the RNA-DNA FISH for all three *Nanog* enhancers. Temporal relationship between individual eRNAs transcription and *Nanog* gene body transcription.

Further evidence of these enhances coordinated activity came from genomics-based approaches. The analysis of chromatin conformation by Capture-C has demonstrated the contacts between *Nanog* promoter and all three enhancers (Figure S16), allowing us to posit that eRNA co-localisation may be correlated with a co-activation of their transcription in the proximity of the *Nanog* gene locus. To further assess whether transcription at these enhancers occurs synergistically, we quantified the co-localisation of these eRNAs either by pair or for all three transcripts (Figure 5C). This analysis reveals that all pairs tested display (5-15%) of sub-nuclear co-localisation. In some instances, co-localisation was also found for all three eRNAs for both the sense and anti-sense strands (Figures 5C and D). Altogether, these data suggest that transcriptional activity of *Nanog* enhancers is synchronised within a scaffold region for at least 30-45% of the mESCs population.

To explore the temporal coordination between *Nanog* eRNA transcription and nascent mRNA production, we examined the expression of each eRNA species relative to active *Nanog* transcription. All three eRNAs showed a positive correlation between their levels and the accumulation of endogenous *Nanog* intron products (Figures 5E and S17, -45 kb eRNA). This is consistent with our RNA seqFISH analysis, which using a broader spatial window detected local enrichment of *Nanog* eRNA signal around the locus in ON states. Together, these measurements support a coordinated enhancer–promoter activation, while the higher temporal resolution afforded by smFISH reveals additional nuances in the timing of eRNA and intron transcription.

Because the co-labelling of *Nanog* eRNAs and *Nanog* mRNA introns has limitations in defining the order of these transcriptional events, due to potential intron retention mechanism ^51^, and RNA mobility – we sought to refine the temporal relationship between their production. To achieve this, we combined DNA- and RNA-FISH experiments to specifically detect newly synthesised eRNA and intronic RNA at the *Nanog* locus. We employed DNA-FISH probes to label the *Nanog* locus alongside RNA-FISH probes to detect nascent *Nanog* intronic RNA and *Nanog* eRNAs and measured their co-localisation within a 4-pixel (440 nm) radius (Figure 5F and Figure S18, schematic).

Quantitative analysis of eRNA co-localisation with intronic RNA at the *Nanog* locus revealed that the transcriptional cycles of all three enhancers operate independently of *Nanog* gene transcriptional activity (Figure 5G). Notably, *Nanog* eRNA synthesis exhibits oscillatory behaviour regardless of whether the *Nanog* gene is active or inactive.

Further supporting evidence for this lack of coordination between enhancer and gene body transcription came from the analysis of eRNA RNA-FISH combined with RFP detection from the STREAMING-tag cassette (Figure S19). This enables us to quantify more accurately the abundance of *Nanog* enhancer transcripts in relation to the progression of *Nanog* elongation (Figure S19). This approach shows that in the “on” state, while nascent *Nanog* mRNA is actively produced, eRNA presence is rarely co-detected for any of these enhancers. By contrast, in the “off” states, there is a wide range of eRNA molecules found in the nucleus (Figures S19A and B), further confirming that enhancer transcription cycles independently of the *Nanog* mRNA synthesis cycle and occurs at a much faster rate. This observation is consistent with the fact idea that local fluctuations in eRNA density at the *Nanog* locus may modulate SOX2 mobility or dwell time to favour priming or prevent molecular interference with the basal machinery.

Integrating live imaging, fixed-cell quantification, and biochemical pulldown, we defined four distinct *Nanog* locus states, each associated with different crowding environments and clustering dynamics (Figure S18). Together, these findings reveal that SOX2 interacts with *Nanog* eRNAs and mRNA but also that the balance of regulatory factors and RNA concentrations shapes different mode of chromatin engagement. We propose a model in which SOX2 priming activity is coordinated with transcriptional bursts through distinct crowding-dependent states, whereby local RNA has a biphasic role either as a negative feedback to trap SOX2 during elongation and prevent interference with the basal machinery or at lower concentration would favour SOX2 chromatin repetitive rebinding contributing to promote Pol II eviction and reset of the locus (Figure 6).

**Figure 6.**
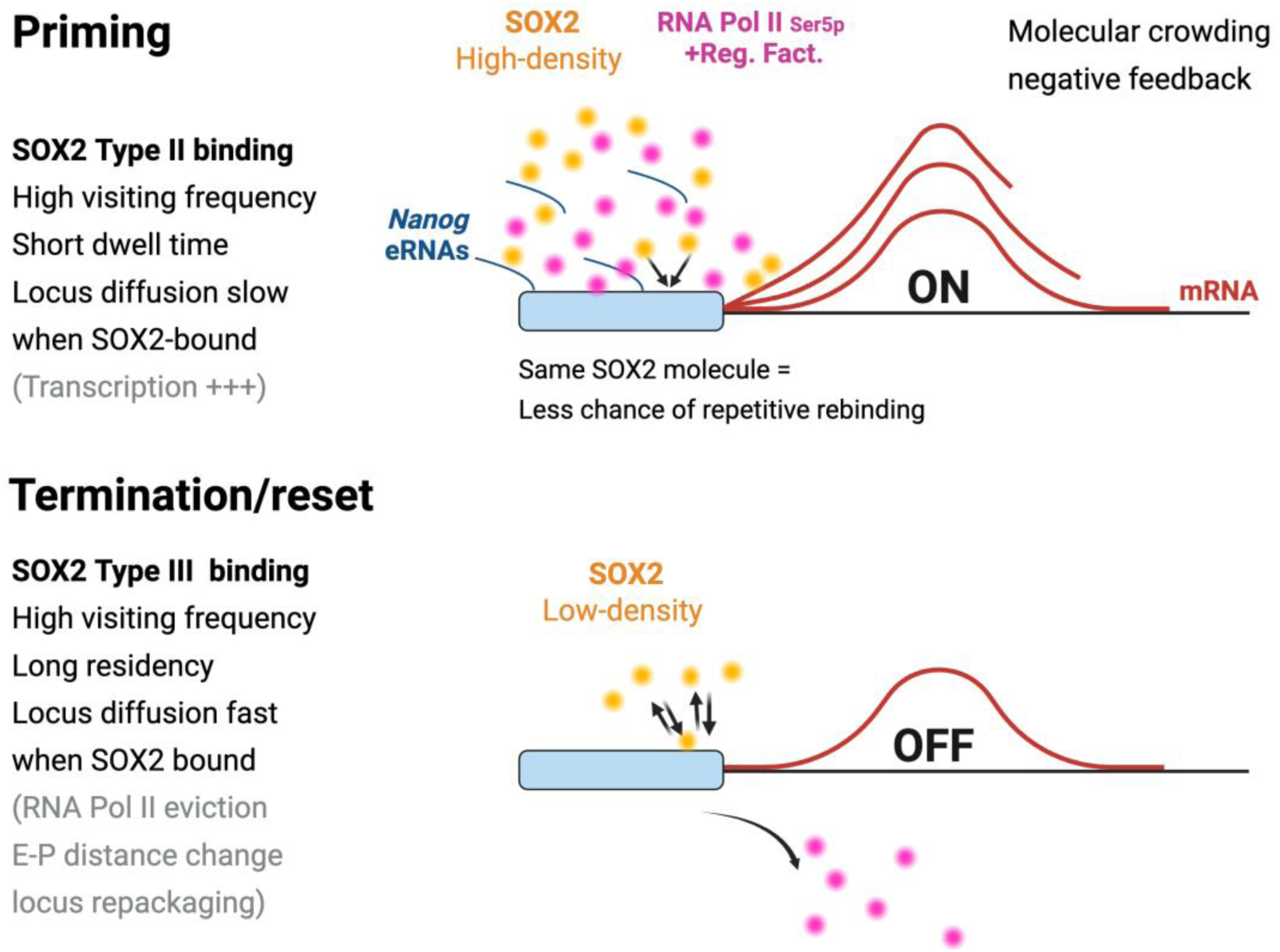
Schematic model illustrating SOX2 Type II and Type III behaviours in relation to chromatin crowding dynamics. Type II events more occurs during the priming state, during which SOX2 transiently samples the locus in a high-crowding environment limited by local RNA and protein assemblies. These short, frequent visits occur while the chromatin remains relatively accessible and transcriptionally permissive. As transcription proceeds into productive elongation, local molecular crowding around the *Nanog* locus increases and SOX2 binding becomes infrequent and highly transient. This reduced SOX2 occupancy prepares the locus for the chromatin repackaging phase marked by Type III long-lived binding. In this state, SOX2 engages in prolonged, stable interactions linked to post-transcriptional chromatin restructuring. This termination mode reflects a lower-crowding microenvironment, where SOX2 binding coincides with chromatin compaction eviction of Pol II and repositioning as the locus transitions toward the next transcriptional cycle.

In conclusion, our findings strengthen the link between the *Nanog* gene transcription cycle and the activity of its regulatory factors by integrating spatial and temporal SOX2, OCT4, MED22 and BRD4 clustering dynamics, locus mobility, enhancer transcription, and fluctuation in crowding states. This study establishes that SOX2 temporal clustering is out-of-phase with the active basal-transcriptional machinery to determine either the basis of permissive transcriptional "off" state that precedes transcription burst or a step that contributes to elongation termination or the early initiation of the next cycle (Figure 6, model).

## Discussion

Previous studies have provided insights into transcription factor (TF) binding dynamics during active transcription, primarily focusing on yeast (e.g., Gal4/Gal10 system) ^52^ and mammalian cells with nuclear receptors like the glucocorticoid receptor (GR) on an exogenous MMTV promoter ^53^. The Gal4-Gal10 system is a valuable model for understanding basic principles of transcriptional regulation, but transcription in yeast is often more straightforward due to fewer regulatory layers since the enhancer network is far less complex. By contrast, here we shed light on how the complex architecture of the endogenous *Nanog* locus shapes TFs visiting frequency and residency time throughout the transcription cycle.

Platania et al., ^44^ have recently reported that gene locus diffusion is a better indicator than Hi-C genomics to predict enhancer to promoter distance. Our approach using the STREAMING-tag reporter system enables a unique integration of locus diffusion data alongside transcriptional activity, enhancing the discovery of SOX2 clustering roles when chromatin is in different states. Findings from this study reveal that SOX2 behaviour is temporally distinct across different transcription cycle stages, a temporal resolution which could not be achieved in previous work ^52–55^.

A limitation of our imaging approach is that TFs, nascent RNA, and DNA could not be visualised simultaneously in a single acquisition. Instead, we relied on near-simultaneous sequential imaging and computational registration to integrate these modalities (3 ms lag). While this strategy provided high spatial and temporal resolution, it may miss transient TF binding events during RNA or DNA acquisition and underestimate short-lived interactions. Moreover, although robust alignment was applied, small registration errors could influence precise TF–RNA–DNA colocalisation. These are common challenges in multimodal single-molecule imaging pipelines, which is why our conclusions emphasise population-level trends and correlative relationships rather than single synchronous events. Importantly, the agreement across independent imaging modes (Mode 1 and Mode 2, described above) helps mitigate these technical constraints and supports the robustness of our findings.

The TF interaction lifetime pattern in yeast model system is typically synchronous with transcription bursts. Here, we observed an anti-phased relationship between SOX2 chromatin binding dynamics and the elongation phase, supporting emerging evidence of an inverse relationship between enhancer-bound condensates and active transcription ^56^. Unlike Du et al., who focused on basal transcriptional machinery condensate dynamics (e.g., RNA Pol II, Mediator) at the *Sox2* locus, our work directly investigates the pioneer TF SOX2. This distinction is critical, as SOX2’s regulatory function and influence on chromatin accessibility represent a more specialised aspect of gene regulation, distinct from the broader transcriptional machinery dynamics. Thus, our study offers a novel perspective on SOX2’s specific contributions to transcriptional regulation and chromatin engagement.

Our study demonstrates that SOX2 medium and high intensity-visiting frequency at the *Nanog* gene locus frame transcription burst events and do not appear to directly cooperate with the BRD4-associated elongation step. This finding fits with the observation that, in stem cells, SOX2 gain-of-function leads to a reduction in *Nanog* mRNA levels and a loss of pluripotency ^14,17^. In that instance, overcrowding of the *Nanog* locus may interfere with the elongation kinetics that does not require frequent SOX2 revisiting, disturb the pre-mRNA splicing ^57^ and derail the cycling pace of *Nanog* transcription required to drive pluripotency. Intriguingly, we observed oscillatory SOX2 behaviours akin to the GR molecular behaviour previously reported ^53^, however, for SOX2 this occurs independently of ligand activation or hormonal cues. This is an unexpected finding, as neither an endogenous ligand nor any pulsatile regulation has been identified to control SOX2 activity. Our finding suggests the existence of a ligand-independent mechanism governing the clustering and oscillatory dynamics of developmental TFs like SOX2.

The observation of different locus mobility responses observed during the "off" state in the absence or presence of SOX2 clusters, is consistent with the observation that binding of the NuRD repressor complex increases *Nanog* locus mobility through nucleosome repackaging ^58^ and also with the fact that SOX2 displays a pioneer activity which relies on binding to compacted nucleosome ^59^. Since it has been shown that SOX2 and NuRD interact ^60,61^, it is tempting to speculate that SOX2 type III clustering behaviour that occurs post-elongation might be associated with NuRD, however pharmacological interference with BPK-25 suggested that SOX2 type III behaviour is independent from NuRD activity. More broadly, these distinct SOX2 clustering activities are likely to indirectly represent changes in protein partners, variations in RNA concentration and chromatin epigenetic state at the locus that modulate SOX2 interaction lifetime ^10,28,62–64^.

This study further informs earlier work that found a role for the interaction between SOX2 and RNA ^28^ and the finding that most TFs have an RNA-binding arm that helps generate an RNA-containing condensate that affects the diffusion and dwell time of genome effectors ^46^. Interestingly, the fast pace of *Nanog* eRNA transcription makes it highly probable to coincide with SOX2 high-visiting frequency. This observation suggests that SOX2 TF must keep operating at the *Nanog* locus in an environment continuously crowded with RNA, either in the presence of eRNA or mRNA which concentration oscillates. Yet, SOX2 chromatin binding dynamics at the *Nanog* locus display multiple behaviours (type I-III), depending on whether elongation is “on” or “off” and whether it is preceding or occurring after the elongation step. This means that different types of RNA at the gene site, which change in concentration over time, are critical in determining how SOX2 behaves ^28,46^. Here, we provide multiple lines of evidence that different SOX2-dependent molecular reactions represent varying “off” states at the *Nanog* locus region and that RNA has a biphasic effect on SOX2 molecular persistence. These molecular reactions involve eRNAs, which have been shown to shape chromatin loops and recruit co-activators ^47,65,66^, likely influencing transcriptional output of *Nanog* in a cooperative manner with SOX2 clustering dynamics.

In conclusion, we posit that cell-to-cell *Nanog* transcriptional fluctuation may be driven by asynchronous variations of SOX2 chromatin binding dynamics, eRNA transcription cycles, and mRNA bursts frequencies and intensities, producing a continuum of *Nanog* expression levels across the stem cell population. These varying synchronicities between genome effectors may give rise to a range of transcription rates, leading either towards self-renewal or differentiation.

## Method

### Cell lines

Genetically modified (STREAMING-tag knocked-in) mESCs were cultured as described ^19^. In brief, all mESC lines were cultured in 2i medium (Knockout Dulbecco’s modified Eagle’s medium [DMEM, Thermo Fisher Scientific Australia Pty Ltd, AU, 10829018]; 15% fetal bovine serum [FBS, Thermo Fisher Scientific, USA, 16141079]; 0.1 mM 2-Mercaptoethanol [Sigma-Aldrich, USA, M3148-25ML]; 1×MEM nonessential amino acids [Thermo Fisher Scientific Australia Pty Ltd, 11140050]; 2 mM L-glutamine solution [Thermo Fisher Scientific Australia Pty Ltd, 25030081]; 1,000 U/mL leukemia inhibitory factor [Sigma-Aldrich, ESG1107]; 20 µg/ml Penicillin-Streptomycin [Thermo Fisher Scientific Australia Pty Ltd, 15140122]; 3 µM CHIR99021 [Sigma-Aldrich, SML1046-5MG]; and 1 µM PD0325901 [Sigma-Aldrich, PZ0162-5MG]) on 0.1% gelatin (Sigma-Aldrich, G1890-100G)-coated T25 flasks at 37°C and 5% CO_2_.

HaloTag-Human SOX2 KI cell line (HaloTag-hSOX2 KI cell), in which the endogenous single exon *Sox2* is biallelically replaced with *hSOX2* cDNA, were established using CRISPR-Cas9 expression vectors (https://benchling.com/s/seq-5qSmelnJy4kN3pPttBor?m=slm-Mcd0jPctZwhu5s453hCC; https://benchling.com/s/seq-wGlQp3vgsvlH7GJPB4Hn?m=slm-xz8aFivYu18VyIbZ81co) and a targeting vector (https://benchling.com/s/seq-63w0RtPOOFpQKTe1Xb25?m=slm-Bri31KJayf3dpQUFsIzX). C57BL/6NCr mESCs (derived from inbred mice, C57BL/6NCr, Japan SLC, Hamamatsu, Japan; 1.25 × 10^5^) were conditioned in 2i medium and plated onto gelatin-coated 24-well plates. After 1 hour, the cells were transfected using Lipofectamine 3000 (Life Technologies, Gaithersburg, MD, L3000015) with 250 ng of the targeting vector, 400 ng of each CRISPR vector, and 75 ng of pKLV-PGKpuro2ABFP (puromycin resistant, Addgene, plasmid #122372), following the manufacturer’s instructions. Puromycin (1 µg/ml) was added to the 2i medium 24 hours post-transfection for selection. The medium was replaced every 2 days, with the first exchange occurring 24 hours after puromycin addition. Seven days post-transfection, cells were treated with 300 µM JFX_549_-Halo-Tag ligand for 30 minutes, followed by three washes and a 30-minute culture in 2i medium. Cells were then trypsinised, analysed via FACS, and Halo-Tag-positive cells were sorted and seeded onto a gelatin-coated 6-cm dish. The medium was changed every 2 days. One week after sorting, 16 colonies were picked for downstream analysis and gene targeting verification. Polymerase chain reaction (PCR) was conducted using primers outside the homology arms, and clones that appeared to have successful biallelic knock-in were selected. These candidate clones underwent further analysis by Southern blotting. This cell line was maintained in 2i medium on 0.1% gelatin.

### HCR™ RNA-FISH

mESCs (1×10^5^) were seeded onto a laminin 511 (BioLamina, Sweden, LN511-0502)-coated 8-well chambered dish (Ibidi, Germany, 80807-90) and cultured for 1.5 h at 37°C and 5% CO_2_. Cells were washed with 1×Dulbecco’s phosphate-buffered saline (DPBS, Thermo Fisher Scientific Australia Pty Ltd, 14190250) 3 times and fixed with 4% paraformaldehyde (Emgrid Australia, AU, 15710) for 10 min at room temperature (RT). Then, cells were washed twice with 1×DPBS and stored in 70% ethanol at - 30°C. After aspirating ethanol at RT, cells were washed with 2×SSC buffer (Molecular Instruments Inc., Los Angeles, CA, USA) and incubated with probe hybridisation buffer (Molecular Instruments Inc.) for 30 min at 37°C. Cells were then hybridised with 1.2 pmol of each probe set dissolved in probe hybridisation buffer at 37°C overnight. The excess probe was removed by washing the cells 5 times with probe wash buffer (Molecular Instruments Inc.) at 37°C. Then, cells were washed twice with 5×SSCT (Molecular Instruments Inc.) buffer at RT. After incubation in the amplification buffer (Molecular Instruments Inc.) for 30 min at RT, the cells were hybridised with 12 pmol of pre-heated and snap-cooled hairpin h1 and h2 probes for 3 h at RT in the dark. The excess hairpin probes were removed by washing the cells 5 times with 5×SSCT buffer at RT. The hybridised cells were incubated with Hoechst33342 (1:125, Thermo Fisher Scientific Australia Pty Ltd, R37605) for 3 min, washed 3 times with 1×DPBS and then stored in 1×DPBS for imaging. Imaging acquisition was performed on a 3i Multimodal microscope (Intelligent Imaging Innovations, Inc., Denver, CO, USA) with a Spinning Disk Confocal (SDC) unit (CSU-W1, Yokogawa Electric Corporation, Tokyo, Japan), a 50 µm pinhole disk, a 100× 1.49 NA Oil Nikon HP TIRF objective (Nikon Instruments Inc., Melville, NY, USA) and a Prime 95B back illuminated sCMOS camera (Teledyne Photometrics, Tucson, AZ, USA), controlled by SlideBook software (Intelligent Imaging Innovations, Inc.). Far-red smFISH probes were imaged with a 640 nm excitation, a 630-644 nm dichroic and a 692/40 emission filter. Red smFISH probes were imaged with a 560 nm excitation, a 555-564 nm dichroic and a 617/73 emission filter. Green smFISH probes were imaged with a 488 nm excitation, a 485-493 nm dichroic and a 525/30 emission filter. Hoechst33342 was imaged with a 405 nm excitation, a 410-410 nm dichroic and a 445/45 emission filter. Images (512 by 512 pixels) were collected with a pixel size of 110 nm and with a Z-stack step size of 2.0 μm (typical range of 15 μm). Images were processed by Fiji software (NIH, Bethesda, MD, USA) ^67^ and analysed using the customised CellProfiler pipeline ^68^. The smFISH spots of each illumination channel were quantified through the customised MATLAB pipeline ^69^. smFISH probes were obtained from Molecular Instruments Inc. mESCs (3×10^4^) were seeded onto a 0.5% gelatin (Sigma-Aldrich, G1890-100G)-coated 30 mm round coverslip and cultured for 1.5 h at 37°C and 5% CO_2_. smFISH staining was performed on cells as described above. Images were acquired using a laser scanning Nikon Eclipse C2 (Nikon Instruments Inc., Melville, NY, USA) confocal microscope, with a 100× 1.40 NA Oil Nikon objective, controlled by Nikon Imaging Software (NIS). A 640 nm laser was used to excite the Far-red smFISH probes (dichroic 405/488/564/640, emission filter 700/80), MCP-RFP was excited with a 560 nm laser line (dichroic 405/488/564/640, emission filter 595/40), a 488 nm laser was used to excite mTetR-GFP (dichroic 405/488/564/640, emission filter 525/50), and a 405 nm laser was used to excite the DAPI channel (dichroic 405/488/564/640, emission filter 447/60). A detector gains of 20 was used for the 640 nm channel; a gain value of 30 was used for the 560 nm labelled samples; the 488 nm channel was imaged using a gain of 25; a gain value of 23 was used for the DAPI channel. No frame accumulation or frame integration was used. Offsets were kept at 0. A pinhole size of 60 µm was used. Pixel dwell time was 1.92 µs. Images (512 by 512 pixels) were collected with a pixel size of 90 nm. Images were processed by Fiji software data and then analysed through the customised CellProfiler pipeline. The GFP/RFP and smFISH spots of each illumination channel were quantified through the customised MATLAB pipeline. Of note, a large fraction of cells has a loss of mTetR-GFP spots after fixation, hence we strictly determined the on-state in presence of both RFP and GFP signal co-localisation.

### Sequential smFISH and HCR™ RNA-FISH

Mouse ESCs derived from inbred mice (Bruce 4 C57BL/6 J, male, EMD Millipore, Billerica, MA, USA) (5×10^4^) were seeded onto a laminin 511 (BioLamina, Sweden, LN511-0502)-coated 8-well chambered cover glass with #1.5 glass (Cellvis, Sunnyvale, CA, USA, C8-1.5HN) and cultured for 24 hours at 37°C and 5% CO_2_. Cells were washed with 1× phosphate-buffered saline (PBS, Nacalai Tesque, 14249-24) three times and fixed with 4% paraformaldehyde (Wako Pure Chemical Industries, USA, 168-20955) for 10 minutes at RT. After fixation, cells were washed twice with 1×PBS and stored in 70% ethanol at - 20°C. Ethanol was aspirated at RT, and cells were incubated with 0.5% Triton-X in 1×PBS for 15 minutes at RT. Cells were then washed three times with 1×PBS. Blocking solution (1×PBS, 10 mg/ml UltraPure BSA [Thermo Fisher Scientific, AM2616], 0.3% Triton-X, 0.1% Dextran sulfate, 0.5 mg/ml UltraPure Salmon Sperm DNA Solution [Thermo Fisher Scientific, 15-632-011]) was added, and cells were incubated for 15 minutes at RT. After 1×PBS washing, cells were post-fixed with 4% formaldehyde in 1×PBS at RT for 5 minutes, followed by six washes with 1×PBS. Cells were incubated at RT for 15 minutes. Samples were then further post-fixed with 1.5 mM BS(PEG)5 (PEGylated bis(sulfosuccinimidyl)suberate) [Thermo Fisher Scientific, A35396] in 1×PBS at RT for 20 minutes, followed by quenching with 100 mM Tris-HCl pH 7.5 at RT for 5 minutes. Cells were washed with 1×PBS and 2×SSC buffer. Cells were then incubated with probe hybridisation buffer (50% formamide, 10% dextran sulfate, 2×SSC, 10 nM each *Nanog* intron probes) for 24 hours at 37°C in a humidified chamber. Cells were washed with 55% wash buffer (55% formamide, 0.1% Triton-X, 2×SSC) for 30 minutes at RT and subsequently washed three times with 4×SSC buffer. Cells were incubated with 200 µL EC buffer (10% ethylene carbonate, 10% Dextran sulfate, 4×SSC, 100 nM readout probe, 150 nM secondary probe) for 20 minutes, followed by washing with 200 µL 4×SSCT buffer (4×SSC, 0.1% Triton-X). Cells were then washed with 200 µL 12.5% wash buffer (12.5% formamide, 0.1% Triton-X, 2×SSC) and 200 µL 4×SSC, followed by washing with 200 µL DAPI solution (4×SSC, 0.1% Triton-X, 5 µg/ml DAPI) for 30 seconds. Finally, cells were washed with 200 µL anti-bleaching buffer (50 mM Tris-HCl, 300 mM NaCl, 3 mM Trolox, 0.8% glucose, 10 µg/ml catalase [Sigma-Aldrich, C3155], pH 8.0) before imaging. Imaging acquisition was performed using a Nikon Ti-2 microscope with a CSU-W1 confocal unit (Yokogawa), a 100× 1.4 NA Nikon Plan Apo λ oil-immersion objective lens, and an iXon Ultra EMCCD camera (Andor Technology, Belfast, UK), with laser lines of 405 nm, 488 nm, 561 nm, and 637 nm. Z-stack images spanning 10 μm with 200 nm intervals (51 sections; 130 nm/pixel) were acquired.

Cells previously processed for smFISH were further subjected to HCR™ RNA-FISH staining, as described in the “HCR™ RNA-FISH” subsection. The hybridised cells were incubated with DAPI solution for 3 minutes, washed with anti-bleaching buffer. Imaging acquisition was performed using a Nikon Ti-2 microscope with a CSU-W1 confocal unit (Yokogawa), a 100× 1.4 NA Nikon Plan Apo λ oil-immersion objective lens, and an iXon Ultra EMCCD camera (Andor Technology, Oxford Instruments, Belfast, UK), with laser lines of 405 nm, 488 nm, 561 nm, and 637 nm. Z-stack images spanning 10 μm with 200 nm intervals (51 sections; 130 nm/pixel) were acquired.

Image processing was initiated using PyImageJ to perform several tasks ^70^. First, smFISH images and HCR-FISH images of the same field of view were converted into a single hyperstack image to manage multi-dimensional image data efficiently. Subsequently, bleaching correction was applied to minimise significant fluorescence intensity differences between the two experiments. Next, 3D drift correction was performed to compensate for any movement or displacement of the samples during image acquisition. The images were then subjected to maximum intensity projection. Following this, the “Extract SIFT Correspondences” and “bUnwarpJ” tools were used to correct for nuclear deformation between the two experiments. After these preprocessing steps, the nuclei were segmented using the Cellpose ^71^, which was specifically configured to identify nuclear regions. For spot detection, the Big-FISH ^72^ was employed. This involved several stages, including filtering the images to reduce noise using a Gaussian filter, detecting local maxima to identify potential spots, and applying adaptive thresholds to distinguish true signal spots from background noise. The detected spots were then masked using the nuclear segmentation results to ensure that only spots within the nuclear regions were considered.

### DNA-antibody conjugation

Oligonucleotide–conjugated primary antibodies were generated essentially as described previously ^73^ with minor modifications. Briefly, 100 µg of each antibody [RNA polymerase II Ser2-phosphorylated (RNAPII Ser2P; clone Pc26) and Ser5-phosphorylated (RNAPII Ser5P; clone Pa57)] in PBS was reacted with a 10-fold molar excess of the heterobifunctional crosslinker SM(PEG)₂ (Thermo Fisher Scientific, 22102) dissolved in anhydrous N,N-dimethylformamide for 2 h at 4°C. Excess crosslinker was removed using 7K Zeba spin desalting columns (Thermo Fisher Scientific).

In parallel, 5’-thiol–modified 18-nt DNA oligonucleotides (Eurofins), consisting of an AAA leader and a 15-nt secondary-probe binding sequence, were reduced with 50 mM dithiothreitol in PBS for 2 h at room temperature and purified on NAP-5 columns (Cytiva). The maleimide-activated antibodies were then mixed with an 11-fold molar excess of the reduced oligonucleotides in PBS and incubated overnight at 4°C to obtain DNA–antibody conjugates. The conjugates were washed and concentrated using 50-kDa Amicon Ultra centrifugal filters, and the concentrations of protein and DNA were determined by a BCA Protein Assay Kit (Thermo Fisher Scientific, 23225) and NanoDrop spectrophotometry, respectively.

Successful conjugation and preservation of antibody functionality were verified by SDS–PAGE followed by Coomassie Brilliant Blue staining, which showed the expected mobility shift upon DNA attachment, and by immunofluorescence using fluorophore-labelled secondary antibodies against the host species, confirming robust nuclear staining. Conjugated antibodies were adjusted to vendor-recommended working concentrations in PBS, pooled when required, and stored at −80 °C until use.

### Sequential smFISH, HCR™ RNA-FISH, and DNA-FISH and immunofluorescence

Sequential smFISH, HCR™ RNA-FISH, and DNA-FISH, and immunofluorescence were performed to visualise the spatial relationship between genomic loci and among nascent and mature RNA transcripts with specific proteins/post-translational modifications within the same cells. Unlike Sequential smFISH and HCR™ RNA-FISH, this method enables the simultaneous detection of *Nanog* intron RNA and exon RNA in separate fluorescence channels during the initial smFISH step, followed by HCR™ RNA-FISH for eRNA detection and DNA-FISH to identify genomic loci, and immunofluorescence to visualise specific proteins/post-translational modifications. The DNA-FISH probes used in this study were the same as previously described ^73^.

For experiments combining RNA/DNA-FISH with immunofluorescence, cells grown on laminin-511-coated glass-bottom chambers were fixed using a dual-fixation protocol optimised to preserve transcription factor localisation. Cells were rinsed once in DPBS and incubated in Fix A [2% (v/v) glyoxal, 5 mM MgCl₂, 0.25 M sucrose in 1× DPBS] for 7 min at 37°C. After a 5-min wash in DPBS containing 5 mM MgCl₂ and 0.25 M sucrose, cells were fixed for 10 min at room temperature in Fix B [4% (w/v) paraformaldehyde, 5 mM MgCl₂, 0.25 M sucrose in 1× DPBS], washed three times in PBS and stored in 70% ethanol at −20°C until further processing.

For immunostaining, samples were rehydrated in PBS, permeabilised in 0.5% Triton X-100 in PBS for 15 min at room temperature, and blocked for 15 min in blocking buffer (1× PBS, 10 mg/ml BSA, 0.3% Triton X-100, 0.1% dextran sulfate, 0.5 mg ml⁻¹ salmon sperm DNA). DNA oligonucleotide–conjugated primary antibodies against SOX2 (R&D, AF2018), BRD4 (Abcam, ab182446) ^73^, RNAPII Ser2P (clone Pc26), and RNAPII Ser5P (clone Pa57) were diluted in blocking buffer and incubated with the samples overnight at 4°C. The next day, samples were washed in PBS, post-fixed in freshly prepared 4% paraformaldehyde in PBS for 5 min, washed again, and further crosslinked in 1.5 mM BS(PEG)₅ in PBS for 20 min before quenching with 100 mM Tris-HCl (pH 7.5) for 5 min. After these steps, samples were processed for smFISH and HCR™ RNA-FISH as described above. After imaging of HCR™ RNA-FISH, samples were processed for DNA-FISH probe hybridisation. The specimens were first washed in PBS and incubated at 37°C for 1 hour with RNase A/T1 Mix (Thermo Fisher Scientific, EN0551, diluted 1:100). Following this treatment, samples were washed three times in PBS and an additional three times in a 50% denaturation buffer (50% formamide, 2× SSC) before incubation at room temperature for 15 min. The specimens were then denatured by heating at 90°C for 4.5 min in 50% denaturation buffer, with the chamber sealed using a PCR seal. Post-denaturation, the samples were rinsed in 2× SSC.

For primary hybridisation, DNA-FISH probes (final concentration: ∼1 nM per probe) were diluted in hybridisation buffer (40% formamide, 2× SSC, 10% (w/v) dextran sulfate) and incubated at 37°C for 96 hours in a humidified chamber. Following hybridisation, samples were washed with 40% wash buffer (40% formamide, 2× SSC, 0.1% Triton X-100) at room temperature for 15 min, followed by three additional washes in 4× SSC.

To stabilise the primary probes, samples underwent a padlock hybridisation step using a 31-nt global ligation bridge (100 nM final concentration) in 20% hybridisation buffer (20% formamide, 10% dextran sulfate, 4× SSC) at 37°C for 2 hours. This was followed by three washes in 10% wash buffer (10% formamide, 2× SSC, 0.1% Triton X-100) at room temperature for 15 min. Ligation was performed using Quick Ligation Kit (New England Biolabs, M2200, 1:20 dilution) supplemented with 1 mM ATP (Takara Bio, 4041) at room temperature for 1 hour. After ligation, samples were washed, fixed, and further processed for amine modification to stabilise the primary probes.

For secondary hybridisation, 200 μl of 100 nM secondary and readout probes in 10% EC buffer (containing ethylene carbonate, dextran sulfate, and 4× SSC) was applied and incubated at room temperature for 20 min. This was followed by sequential washes with 4× SSCT (4× SSC, 0.1% Triton X-100) and 12.5% wash buffer (12.5% formamide, 2× SSC, 0.1% Triton X-100). DAPI staining was performed by incubating the samples in DAPI solution (4× SSCT with 1:100 diluted DAPI; Dojindo, Japan, 340-07971) for 30 seconds at room temperature. Finally, an anti-bleaching buffer containing 50 mM Tris-HCl (pH 8.0), 300 mM NaCl, 2× SSC, 3 mM Trolox (Sigma-Aldrich, 238813), 0.8% D-(+)-glucose (Nacalai Tesque, Japan, 16806-25), catalase (Sigma-Aldrich, C3155, 1:1000 dilution), and glucose oxidase (0.5 mg/ml; Sigma-Aldrich, G2133) was applied, followed by imaging 150 seconds later.

Imaging acquisition was performed using a Nikon Ti-2 microscope with a CSU-W1 confocal unit (Yokogawa), a 60× 1.3 NA Nikon Plan Apo λ silicone-immersion objective lens, and an ORCA-Fusion BT sCMOS camera (Hamamatsu Photonics), with laser lines of 405 nm, 488 nm, 561 nm, and 637 nm. Z-stack images spanning 15 μm with 300 nm intervals (51 sections; 110 nm/pixel) were acquired.

Image processing was performed using the same pipeline as described for Sequential smFISH and HCR™ RNA-FISH, including drift correction, bleaching correction, maximum intensity projection, nuclear segmentation using Cellpose, and spot detection using Big-FISH.

### Live cell imaging

For live cell imaging, SNAP-tag SOX2 KI mESCs (5×10^4^) were transferred onto a laminin 511-coated 8-well chambered dish and cultured in 2i medium for 2 h. Then, cells were incubated with JF_646_-SNAP-tag ligand (5 nM, LAVIS LAB, Ashburn, USA, JBG-13-151) dissolved in imaging 2i medium (FluoroBrite DMEM [Thermo Fisher Scientific Australia Pty Ltd,. A1896701], 15% FBS [Thermo Fisher Scientific, USA, 16141079], 1× MEM nonessential amino acids [Thermo Fisher Scientific Australia Pty Ltd, 11140050], 2 mM L-glutamine solution [Thermo Fisher Scientific Australia Pty Ltd, 25030081], 1,000 U/mL LIF [Sigma-Aldrich, ESG1107], 3 µM CHIR99021 [Sigma-Aldrich, SML1046-5MG], 1 µM PD0325901 [Sigma-Aldrich, PZ0162-5MG]) for 30 min at 37°C and 5% CO_2_ in the dark. The excess SNAP-tag ligand was removed by washing the cells 3 times with the imaging 2i medium. Cells were imaged in the 2i medium containing VectaCell Trolox Antifade Reagent (1:500, ABACUS DX, AU, CB-1000) and Hoechst33342 (1:12.5, Thermo Fisher Scientific Australia Pty Ltd, R37605). Time-lapses were acquired using a 3i Multimodal microscope (Intelligent Imaging Innovations, Inc., Denver, CO, USA) equipped with a CSU-W1 Spinning Disk Confocal (SDC) unit (50 µm disk unit), a spinning TIRF (Vector2 TIRF, Intelligent Imaging Innovations, Inc.) unit, a high speed (3 ms) galvo port switcher (mSwitcher, Intelligent Imaging Innovations, Inc.) to select between SDC or HILO modalities, a 100× 1.49 NA Oil Nikon HP TIRF objective (Nikon Instruments Inc.) and Prime 95B back illuminated sCMOS cameras (Teledyne Photometrics). Microscope system was controlled by the SlideBook software (Intelligent Imaging Innovations). STREAMING-tagged reporters were imaged in SDC mode using the following filter settings: MCP-RFP was excited with a 560 nm laser line (dichroic ZT405/488/561/640tpc-uf1, emission filter FF02-617/73), a 488 nm laser was used to excite mTetR-GFP (ZT405/488/561/640tpc-uf1, emission filter FF01-525/30), and HILO SMT imaging was performed using a 640 nm laser, a Di03-R405/488/561/635 dichroic mirror and a FF01-692/40 for emission filter. For image acquisition method 1, the microscopy was set up first for SDC image acquisition (1 frame each, 300 ms camera exposure each channel, no delay time) and then shifted to the HILO-SMT module for slow tracking of fluorescent molecules (30 frames, 500 ms camera exposure, no delay time) to determine interaction lifetime kinetics. The imaging cycle was repeated 10 times with a total duration of 3 min. For image acquisition method 2, one frame in HILO-SMT mode was acquired first using a camera exposure time of 500 ms, followed by the acquisition of 3 frames of the SDC channels (no interval time). The imaging cycle was repeated 170 times for 10 min.

TetraSpeck 0.5 µm fluorescent beads (Thermo Fisher Scientific Australia Pty Ltd, T7281) were used as fiducials to correct for camera misalignment between the SDC and HILO imaging modules. Beads were plated on a laminin 511-coated well with imaging 2i medium and imaged under the same conditions as described above (green and red beads were imaged using the SDC camera, dark-red beads were imaged using the HILO-SMT imaging settings). The shifted distance between the SDC channels (GFP and RFP) and the HILO-SMT channel was realigned based on the central position of the beads in each channel.

### TFs dwell time analysis

The lifetime of the TFs in each cell was computed using two-component exponential decay curves as the following equation ^74^:

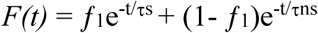

The lifetime was adjusted by the bleach rate (BR) as the following equation:

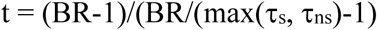

BR was computed by taking the inverse of the average intensity decay rate (I_DR_) ^10^:

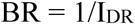

I_DR_ was computed as the following equation:

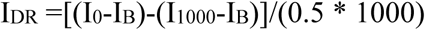

Where I_0_ is the intensity at the first frame, IB is the background intensity, I_1000_ is the intensity after a thousand frames.

### TFs residency time analysis

Fluorescent molecules were identified using a custom-written MATLAB implementation of the MTT algorithm. The parameters used in the pipeline were: localisation error: 10-6.5, emission wavelength: 580 nm, exposure time: 500 ms, number of deflation loops: 0, number of maximum expected diffusion constant: 0.2 μm^2^/s, number of gaps allowed: 1, point scale factor scaling: 1.35, detection box size: 9, minimum intensity: 0, maximum optimisation iteration: 50, position tolerance: 1.5, search exploration factor: 1.2, number of past frames data used in trajectories: 10, maximum trajectories competitors: 3, intensity probability weighting: 0.9 and diffusion probability weighting: 0.5. The intensity decay due to photobleaching was corrected during the TFs interaction lifetime analysis.

GFP/RFP spots were detected by the signal intensity. The intensity of acquired images was segmented using Otsu’s method. The pixels with the highest few segments were grouped into blobs based on their connectivity. The detected GFP and RFP signal was filtered by the pixel size (less than 10% of the image size).

On-state was defined when the post-aligned RFP signal appeared within the 4-pixel radius of the GFP signal. SOX2 was defined as colocalised with the GFP signal when appeared within the 4-pixel radius region. The GFP signal was detected in each acquired frame and used as a reference for the region of interest (4-pixel radius) in the subsequent frame for detecting RFP and SOX2 co-localisation.

### Locus mobility analysis

For locus mobility analysis, SNAP-tag SOX2 KI mESCs were plated into an 8-well chambered dish and cultured in 2i medium for 2 h. Then, mESCs were cultured in the imaging 2i medium for live imaging as described in the “Live cell imaging” subsection. The acquired images were processed through the customised MATLAB pipeline for GFP and SOX2 detection in each frame. The movement of detectable GFP spots (in pixels) between adjacent frames was recorded. The mean locus mobility changes with or without SOX2 binding events were calculated in each cell.

### Dual HILO imaging

HaloTag-hSOX2 KI mESCs (5×10^4^) were transferred onto a laminin 511-coated 8-well chambered dish and cultured in 2i medium for 2 h. Then, cells were incubated with JF_549_-Halo-Tag ligand (5 nM, LAVIS LAB, Ashburn, USA, NF-1-183) and with JF_646_-Halo-Tag ligand (250 nM, LAVIS LAB, Ashburn, USA, NF-1-005) dissolved in imaging 2i medium for 30 min at 37°C and 5% CO_2_ in the dark. The excess Halo-Tag ligand was removed by washing the cells 3 times with the imaging 2i medium. Cells were imaged in the 2i medium containing VectaCell Trolox Antifade Reagent and Hoechst33342.

Time-lapses were acquired using a Nikon H-TIRF microscope (Nikon Instruments Inc.), a 100× 1.49 NA Oil Nikon Apo TIRF objective (Nikon Instruments Inc.) and two Sona sCMOS cameras (Andor Technology, Oxford Instruments, Belfast, UK). The emission signal was split using a TuCam adapter (Andor Technology, Oxford Instruments, Belfast, UK) with an ‘FF640, 579/34-25, 679/41-25’ filter cube. The instrument was controlled by the Nikon NIS-Elements software (Laboratory Imaging s.r.o., Praha, Czech Republic). Dual HILO SMT imaging was performed simultaneously using a 560 nm laser line and a 640 nm laser line with a TRF89902-NK TIRF ET quad band filter cube (Chroma Technology Corp, Bellow Falls, VT, USA). The microscopy was set up for HILO image acquisition for slow tracking of fluorescent molecules (150 frames, 500 ms camera exposure, no interval time).

Dual-colour HILO SMT of SOX18 was performed in HeLa cells using the same Nikon H-TIRF microscope platform and acquisition settings described above for SOX2, with modifications only to sample preparation. HeLa cells were cultured in DMEM supplemented with 10% FBS (GE Healthcare, SH31195.01), 1% GlutaMAX (Gibco, Thermo Fisher Scientific, 35050079), and 1% non-essential amino acids (Gibco, Thermo Fisher Scientific, 11140050). HeLa cells were seeded at a density of 30,000 cells per well in 8-well glass-bottom chamber slides (Ibidi, 80827) coated with 0.5% gelatin, 24 h prior to transfection. Transfections were carried out using X-tremeGENE 9 transfection reagent (Roche) with 200 ng of plasmid DNA per well, following the manufacturer’s protocol. FluoroBrite DMEM (Gibco, Thermo Fisher Scientific, A1896702) supplemented with 1% GlutaMAX was used as the low-serum transfection medium (SMT medium). Cells were incubated for at least 12 h post-transfection at 37 °C and 5% CO₂ before imaging. For HaloTag labelling, cells were incubated with JF549-HaloTag ligand (2 nM; LAVIS Lab, Ashburn, USA; NF-1-183) and JF646-HaloTag ligand (250 nM; LAVIS Lab, Ashburn, USA; NF-1-005) diluted in SMT medium for 30 min at 37 °C and 5% CO₂ in the dark. Excess ligand was removed by washing the cells three times with SMT medium.

Time-lapse imaging was performed using the Nikon H-TIRF microscope and was carried out under HILO conditions for slow tracking of fluorescent molecules (150 frames, 500 ms exposure time, no interval).

### TFs cluster analysis

Fluorescent signal from the saturated labelling channel was quantified using a custom-written MATLAB pipeline. The high-, medium- and low- intensity regions in each nucleus were defined based on fluorescent signal intensity (the pixels with highest 8% signal as high-intensity region, the following 30% as medium-intensity region and the rest as low-intensity region). The single molecule detection, as described in the “TFs residency time analysis” subsection, was performed for the sparse labelling channel within various ROIs with a radius of 4 pixels (440 nm), selected from the high-intensity regions. The overlap analysis between sparse and saturated labelling channels were performed using the customised MATLAB pipeline.

### Visiting frequency classification and Type III identification in Mode 2 imaging

In Mode 1 imaging, visiting frequency was calculated as the proportion of frames in which SOX2 binding was detected relative to the total number of frames. Visiting frequencies were classified as low (0 - 3%), medium (3 - 10%), or high (> 10%).

In Mode 2 imaging, SMT frames were acquired at 3.6 second intervals with 500 ms exposure, such that a single long-dwell SOX2 binding event would span approximately 7-8 consecutive frames if continuously tracked. Accordingly, Type III behaviour in Mode 2 was defined as ≥3 detectable SOX2 binding events within a 100-frame window (corresponding to a high visiting frequency), with at least one event exhibiting a long-dwell duration.

### Drug perturbation and live-cell imaging conditions

For perturbation experiments, cells were treated with either 100 nM TSA (Sigma-Aldrich,T1952), 500 nM flavopiridol (Sigma-Aldrich, F3055-5MG), or 5 μM BPK-25 (MedChemExpres, HY-141550), with each drug applied in independent experiments, for 3 h prior to imaging. BPK-25-treated and untreated control samples were imaged using Mode 2 acquisition on the 3i Multimodal microscope, as described above. TSA- and flavopiridol-treated samples were imaged on the Nikon H-TIRF microscope (Nikon Instruments Inc.) equipped with a 100×/1.49 NA Apo TIRF oil objective. SMT imaging was performed using a 640-nm laser and a TRF89902-NK TIRF ET quad-band filter cube (Chroma Technology Corp., Bellows Falls, VT, USA). HILO illumination was used for slow single-molecule tracking (170 frames, 500 ms exposure, 3.6 s interval) to mimic Mode 2 imaging acquisition conditions. The whole-nucleus visiting frequency analysis was performed using a customised MATLAB pipeline.

### Nuclear feature tracking associated with SOX2 binding behaviour

SNAP-tag SOX2 KI mESCs were prepared for live-cell imaging and imaged in 2i imaging medium supplemented with VectaCell Trolox Antifade reagent (1:333, ABACUS DX, AU, CB-1000) and Hoechst 33342 (1:6.25, Thermo Fisher Scientific, R37605). Each cycle consisted of: a single HILO-SMT frame acquired using a 640 nm excitation line, a ZT 405/488/561/640 rpc (60% T 40% R), UF2 dichroic mirror and a FF01-692/40 for emission filter (500 ms exposure, no delay time); followed by one HILO-SMT frame acquired using a 405 nm laser, same dichroic mirror and a FF01-445/20 for emission filter (200 ms exposure, no delay time); and then three sequential spinning-disk confocal (SDC) frames of each of the GFP and RFP channels (300 ms exposure, no delay time). This four-channel imaging cycle was repeated 170 times over 11 minutes.

Images were first processed using the TFs occupancy pipeline to identify cells exhibiting SOX2 Type III behaviour at the *Nanog* locus. For cells showing Type III SOX2 binding, approximately 30 images before and 30 images after the binding event were extracted and designated as pre-binding and post-binding windows. The *Nanog* locus (GFP-labelled) was used to centre a region of interest (ROI) for nuclear feature tracking.

Within this region, nuclear texture was quantified using a 3x3 convolution kernel to compute 252 texture features. To mitigate the cellular heterogeneity, feature values were averaged across 10 random frames per condition and standardised using z-score normalisation (StandardScaler, scikit-learn v1.6.1). Dimensionality reduction was performed using principal component analysis (PCA). A support vector machine was trained to classify this nuclear texture, and its categorical accuracy was evaluated using a random 80% training-20% testing split.

Within each ROI, nuclear texture features were quantified using a 3×3 convolution kernel, generating 252 multiparametric descriptors ^43,75,76^. To account for cell-to-cell variability, feature values were averaged across 10 randomly sampled frames per condition and standardised using z-score normalisation (StandardScaler, scikit-learn v1.6.1). Dimensionality reduction was performed via principal component analysis (PCA). A support vector machine classifier was trained to distinguish pre- and post-binding nuclear states, and its performance was evaluated using an 80% training / 20% testing split.

### Live-cell imaging of SNAP-tagged SOX2 to quantify SOX2 accumulation at the *Nanog* locus during transcriptional “on” and “off” states

SNAP-tag SOX2 KI mESCs (5×10^4^) were transferred onto a laminin 511-coated 8-well chambered dish and cultured in 2i medium for 24 h. Then, cells were incubated with JF_646_-SNAP-tag ligand (300 nM, LAVIS LAB, Ashburn, USA, JBG-13-151) dissolved in imaging 2i medium (FluoroBrite DMEM [Thermo Fisher Scientific Australia Pty Ltd,. A1896701], 15% FBS [Thermo Fisher Scientific, USA, 16141079], 1× MEM nonessential amino acids [Thermo Fisher Scientific Australia Pty Ltd, 11140050], 2 mM L-glutamine solution [Thermo Fisher Scientific Australia Pty Ltd, 25030081], 1,000 U/mL LIF [Sigma-Aldrich, ESG1107], 3 µM CHIR99021 [Sigma-Aldrich, SML1046-5MG], 1 µM PD0325901 [Sigma-Aldrich, PZ0162-5MG]) for 30 min at 37°C and 5% CO_2_ in the dark for 30 min. The excess SNAP-tag ligand was removed by washing the cells 3 times with the imaging 2i medium. Then, cells were further incubated for 30 min at 37°C and 5% CO_2_ in the dark. Cells were imaged in the imaging 2i medium containing VectaCell Trolox Antifade Reagent (1:500, ABACUS DX, AU, CB-1000). Images were acquired using a Nikon Ti-2 microscope with a CSU-W1 confocal unit (Yokogawa), a 100× Nikon Plan Apo λ oil-immersion objective lens (NA 1.4), and an iXon Ultra EMCCD (Andor Technology), operated using NIS-Elements software (ver. 5.11.01; Nikon). The microscope was also equipped with 405, 488, 561, and 637 nm lasers (Andor Technology), and an ASI MS-2000 piezo stage (ASI). Z-stack images spanning 10 µm with 200 nm intervals (21 sections; 130 nm/pixel) were acquired.

For nuclear segmentation and per-cell intensity measurements, raw 3D image stacks were pre-processed using a custom Python pipeline. Briefly, Z-stacks were Gaussian-blurred in the SNAPtag and mTetR channels and projected along Z to generate a nuclear image used for segmentation. In parallel, median-filtered 3D stacks (11 Z sections × 3 channels) were saved as 16-bit TIFF files for downstream quantification. These projected images were segmented with Cellpose in “nucleus” mode, using a fixed object diameter of 100 pixels and the projected nuclear signal as input. Segmentation masks were saved as 16-bit label images and visually inspected; cells with obvious segmentation errors were excluded from further analysis. For each remaining cell, the mean fluorescence intensity of the SNAPtag, MCP and mTetR channels was computed within the nuclear mask on a central Z-slice, together with the nuclear area in pixels, yielding a per-cell summary table used to select cells within three standard deviations of the mean intensity distribution.

To identify the *Nanog* locus in each nucleus, mTetR foci were detected using the Big-FISH Python package. Median-filtered 3D stacks were cropped to the bounding box of each nucleus defined by the Cellpose mask, and the mTetR channel was processed with a 3D Laplacian-of-Gaussian filter. Spots were then detected as local maxima using Big-FISH with a voxel size of 500 × 130 × 130 nm (Z, Y, X) and an effective spot radius of 500 × 300 × 300 nm, parameters chosen to match the measured point-spread function of the optical system. Candidate spots were thresholded based on intensity using an adaptive cut-off derived from the nuclear mTetR intensity distribution, and only those confined within the nuclear mask were retained. For each cell, the brightest mTetR spot was taken as the *Nanog* locus and its 3D coordinates were stored for further analysis. In addition, random control positions were generated by uniformly sampling coordinates within the same nuclear masks at central Z-planes, and 19 × 19-pixel windows centred on these random coordinates were extracted from all three channels to serve as nucleoplasmic background controls.

For each cell with a valid *Nanog* locus, a 19 × 19-pixel (2.5 × 2.5 µm) ROI centred on the mTetR spot was extracted from the SNAPtag, MCP and mTetR channels, yielding a stack of locus-centred ROIs of fixed size. Equivalent ROIs were also extracted around random nuclear positions as described above. Spot detection within each 2D ROI was performed using trackpy. Local maxima in the SNAPtag, MCP and mTetR channels were first identified with a band-pass filter and then refined by 2D Gaussian fitting to obtain sub-pixel centroid positions. For each detected peak, a relative intensity value r was computed as the integrated fluorescence within a circular region of radius 3 pixels divided by the product of the region area and the mean intensity of the ROI.

To define the transcriptional state of each locus, MCP peaks were associated with the corresponding mTetR position. Loci were classified as transcriptionally active (“ON”) when an MCP peak with r_MCP ≥ 1.2 was detected within 3 pixels (390 nm at 130 nm/pixel) of the mTetR centroid; all remaining loci were classified as “OFF”.

To quantify the spatial enrichment of each factor around the *Nanog* locus, we computed radial intensity profiles from the locus-centred ROIs. For each ROI, baseline-subtracted images were obtained by subtracting the mean random-ROI image for the corresponding channel. Radial profiles were then calculated in concentric annuli of width 1 pixel (0.13 µm) around the central pixel up to a radius of 9 pixels (1.17 µm). For each channel, mean ± s.e.m. radial profiles were computed separately for On and Off loci as well as for the random controls, after scaling the images so that the maximum baseline-subtracted pixel value across On and Off conditions corresponded to 100 arbitrary units.

### Capture-C Extraction

Nuclei containing chromatin were isolated and fixed as detailed previously ^77,78^. Briefly, 10 x 10^6^ mouse embryonic stem cells E14 (donation from Rob Klose lab) were trypsinised and collected in 50ml falcon tubes in 9.3 ml medium (Dulbecco’s Modified Eagle Medium supplemented with 15% fetal bovine serum, 2 mM GlutaMAX, 0.5 mM beta-Mercaptoethanol, 1x non-essential amino acids, 1x penicillin/streptomycin solution and 10 ng/mL leukemia-inhibitory factor (produced in-house)) and crosslinked with 1.25 ml 16% formaldehyde (1.89% final, Thermo Fisher Scientific, 10751395) through rotation for 10 min at RT. Crosslinking reaction was quenched with 1.5 ml 1M cold glycine, washed with cold PBS and lysed for 20 min at 4°C in lysis buffer (10 mM Tris, 10 mM NaCl, 0.2% NP-40, cOmplete protease inhibitors [Roche, 05056489001], pH 8). Finally, cells were pelleted and resuspended in 500 µl lysis buffer per 5 x 10^6^ cells and snap frozen on dry ice. Fixed chromatin was stored at - 80°C.

### Capture-C Library Preparation

Capture-C libraries were prepared as described previously ^78^. Briefly, cell lysates from 5 x 10^6^ cells were thawed on ice, pelleted and subsequently resuspended in 250 µl 1x DpnII buffer (New England Biolabs, B0543). Each lysate was treated with SDS (0.28% final concentration, Thermo Fisher Scientific, AM9820) through interval shaking (500 rpm, 30 sec on/off) for 1 h at 37°C, then quenched with Triton X-100 (1.67% final concentration) through interval shaking (500 rpm, 30 sec on/off). Chromatin was digested with 3 x 10 µl DpnII (homemade) through interval shaking (500 rpm, 30 sec on/off) for 24 h at 37°C. 100 µl aliquot was taken to assess the quality of digest on agarose gel upon reverse crosslinking. The remaining chromatin was ligated with 8 µl T4 Ligase (240 U, Thermo Fisher Scientific, EL0013) in a volume of 1440 µl for 22 h at 16°C. Subsequently, the nuclei containing ligated chromatin were pelleted to remove non-nuclear chromatin, reverse-crosslinked and ligated DNA was purified by phenol-chloroform extraction. The resulting DNA was resuspended in 200 µl water and sonicated for 13 cycles (30 sec on/off, Bioruptor Pico) until a fragment size of approximately 200 bp was achieved. Fragments were size selected using Ampure XP beads (selection ratios: 0.85 x / 0.4x, Beckman Coulter, A63881). 1-5 µg DNA was adaptor-ligated and indexed using the NEBNext End Repair Module (New England Biolabs, E6050), dA-Tailing (New England Biolabs, E6053), Quick Ligation modules (New England Biolabs, E6056), and NEBNext Multiplex Oligos for Illumina Primer sets 1 (New England Biolabs, E7335) and 2 (New England Biolabs, E7500S). The libraries were amplified with Herculase II Fusion Polymerase kit (Agilent, 600677) for 6 PCR cycles. The probes for hybridisation (supp Table. 1) were designed in the following manner: two capture probes (aligned to mm10, 70 bp each) were designed for each promoter-containing DpnII digest fragment using the online tool by the Hughes lab ^79^ with filtering parameters as follows: Duplicates: < 2, Density < 30, SRepeatLength < 30, Duplication: FALSE. If no probes could be designed for the restriction fragment directly overlapping the TSS, the next-nearest DpnII fragment was chosen, if it was within 500 bp of the TSS.

Indexed libraries were measured with Qubit dsDNA BR Assay (Invitrogen, Q32850) and pooled by mass (1.5 µg each) prior to hybridisation. Hybridisation was performed with the pool of probes (2.9 nM each) using KAPA HyperCapture Reagent Kit (Roche, 9075828001) and KAPA HyperCapture Bead Kit (Roche, 9075798001) according to Roche protocol for 72 h followed by a 24-hour hybridisation (double capture). Quantification of captured libraries was performed via qPCR using the KAPA Sybr Fast Universal kit (KAPA, KK4602) and sequenced on Illumina NextSeq 550 (Illumina, SY-415-1002) using High-Output v2.5 Kit (75 cycles, Illumina, 20024906).

### Capture-C analysis

Paired-end reads were aligned to mm10 and filtered for Hi-C artefacts using HiCUP v 0.8.3 ^80^ and Bowtie2 v2.3.5.1 ^81^, with fragment filter set to 100 - 800 bp. Read counts of reads aligning to captured gene promoters and interaction scores (significant interactions) were then called by the Bioconductor package Chicago v1.6.0 ^82^. For visualisation of Capture-C data, weighted pooled read counts from Chicago data files were normalised to the total read count aligning to captured gene promoters in the sample and further to the number of promoters in the respective capture experiment. Normalised reads were then multiplied by a constant number to simplify visualisation: (normCounts = counts/cov*nprom*100000). The enrichment data for H3K4mel ^83^ and SOX2 ^84^ were obtained from previous papers.

### RNA immunoprecipitation

RNA immunoprecipitation was performed using an ESC line with a SNAP-tag knocked into the *Sox2* locus to enable specific isolation of SOX2-RNA complexes. ESCs (∼6 × 10⁶ cells per biological replicate) were crosslinked in 1 mL 4% paraformaldehyde (PFA, EMGRID AUSTRALIA, 15710) for 10 min on ice. Crosslinking was quenched by adding glycine to a final concentration of 125 mM for 5 min, and the supernatant was removed. The cell pellet was then lysed in 1 mL of freshly prepared lysis buffer containing 150 mM NaCl (Sigma-Aldrich, S5150-1L), 50 mM Tris-HCl (pH 8), 100 mM MgCl₂ (Sigma-Aldrich, M2670-100G), 25 mM EDTA (Sigma-Aldrich, E6758-100G), 10% glycerol (ThermoFisher Scientific, AJA242-500ML), 1% Triton X-100 (Sigma-Aldrich, T8787-100ML), and 0.1 mM DTT, supplemented with protease inhibitor (Sigma-Aldrich, P8340-1ML) and RNaseOUT™ RNase inhibitor (Thermo Fisher Scientific, 10777019), in nuclease-free water (Thermo Fisher Scientific, 10977015). Cell pellets were resuspended in lysis buffer and incubated on a rotating wheel for 1 h at 4 °C (180 rpm). Lysates were then centrifuged at 10,000 × g for 30 minutes at 4 °C, and the supernatant was collected. The clarified lysate was incubated overnight at 4 °C with 160 µL of pre-washed SNAP-Capture Magnetic Beads (New England Biolabs, S9145S) on a rotating wheel to enrich SOX2-RNA complexes.

Following incubation, bead-bound complexes were separated from the unbound supernatant using magnetic separation. Beads were washed three times with the lysis buffer to remove nonspecific material. The SOX2-enriched (bead-bound) fraction was resuspended in 500 µL of reverse crosslinking buffer (50 mM Tris, pH 8, 150 mM NaCl, 10 mM EDTA, and 1% SDS) containing proteinase K (QIAGEN) and RNase inhibitor (New England Biolabs, M0314L) and incubated for 1 h at 50 °C to release bound RNA. RNA was extracted using 1 mL of TRI reagent (Sigma-Aldrich, T9424-200ML) followed by chloroform-based phase separation according to the manufacturer’s instructions. The isolated RNA was treated with DNase I to remove residual genomic DNA prior to downstream analysis. The cDNA was synthesised from the purified, DNase-treated RNA using random hexamer primers and the ProtoScript® First Strand cDNA Synthesis Kit (New England Biolabs, E6300L). PCR amplification of specific transcripts was performed using gene-specific primers listed in Table S1. PCR amplicons were purified with the Monarch® DNA Gel Extraction Kit (New England Biolabs, T1020) to remove residual primers and nonspecific fragments, and the cleaned products were then sent to the Australian Genome Research Facility (AGRF) for Sanger sequencing.

## Supporting information

Supp video 1, and will be used for the link to the file on the preprint site.

Supp video 2, and will be used for the link to the file on the preprint site.

Supp video 3, and will be used for the link to the file on the preprint site.

Supp video 4, and will be used for the link to the file on the preprint site.

Supp video 5, and will be used for the link to the file on the preprint site.

Supp video 6, and will be used for the link to the file on the preprint site.

## Acknowledgments

We thank Sydney Microscopy and Micro-Analysis core facilities for ongoing technical support with imaging. We thank Dr Alexandros Pertsinidis and Dr Jieru Li (MSK, NY) for their valuable input and feedback on reviewing the manuscript. We thank Dr Luke Lavis Janelia Farm (Virginia Tech) for providing reagents necessary to establish the SOX2 HaloTag knock-in cell lines. We thank the Faulkner laboratory for generously providing HeLa cells. We thank Dr Yew Yan Wong for developing and implementing the custom computational scripts used for image processing and quantitative analysis oupput, and for contributing technical improvements that enhanced analysis efficiency and reproducibility.

## Funding

NHMRC idea Grant (APP 2029719) (MF)

NHMRC idea Grant (APP2019904) (MF)

NSW Cardio-vascular Research Capacity Program – Senior Researcher grant (MF)

The Cooperative Research Project Program of the Medical Institute of Bioregulation, Kyushu University (MF)

Grants-in Aid for Scientific Research from the Ministry of Education, Culture, Sports, Science, and Technology JP24H02326 (HOc)

Grants-in Aid for Scientific Research from the Ministry of Education, Culture, Sports, Science, and Technology JP22H02609 (HOc)

CREST from the Japan Science and Technology Agency to HOc (JPMJCR23N3)

NIG-JOINT 69A2024 (HOc)

NIG-JOINT 14R2024 (HOc)

Medical Research Centre Initiative for High Depth Omics, Kyushu University, Japan (HOc)

MEXT Promotion of Development of a Joint Usage/Research System Project: the cooperative Research Project Program (HOc)

MEXT Promotion of Development of a Joint Usage/Research System Project: Coalition of Universities for Research Excellence Program (CURE) JPMXP1323015486 (HOc)

Takeda Science Foundation (HOc)

Mitsubishi Foundation (HOc)

## Author contributions

Conceptualisation: HOc, MF

Methodology: HOc, BL, YW, NR, TD, YK, Hoh

Investigation: BL, MG, NR, YW, TD, YK, XG, SU

Visualisation: YW, BL, MF, HOc

Funding acquisition: HOc, MF

Project administration: HOc, MF

Supervision: MF, TD

Writing – original draft: MF

Writing – review & editing: MF, BL, HOc

## Extended data

**Figure S1.**
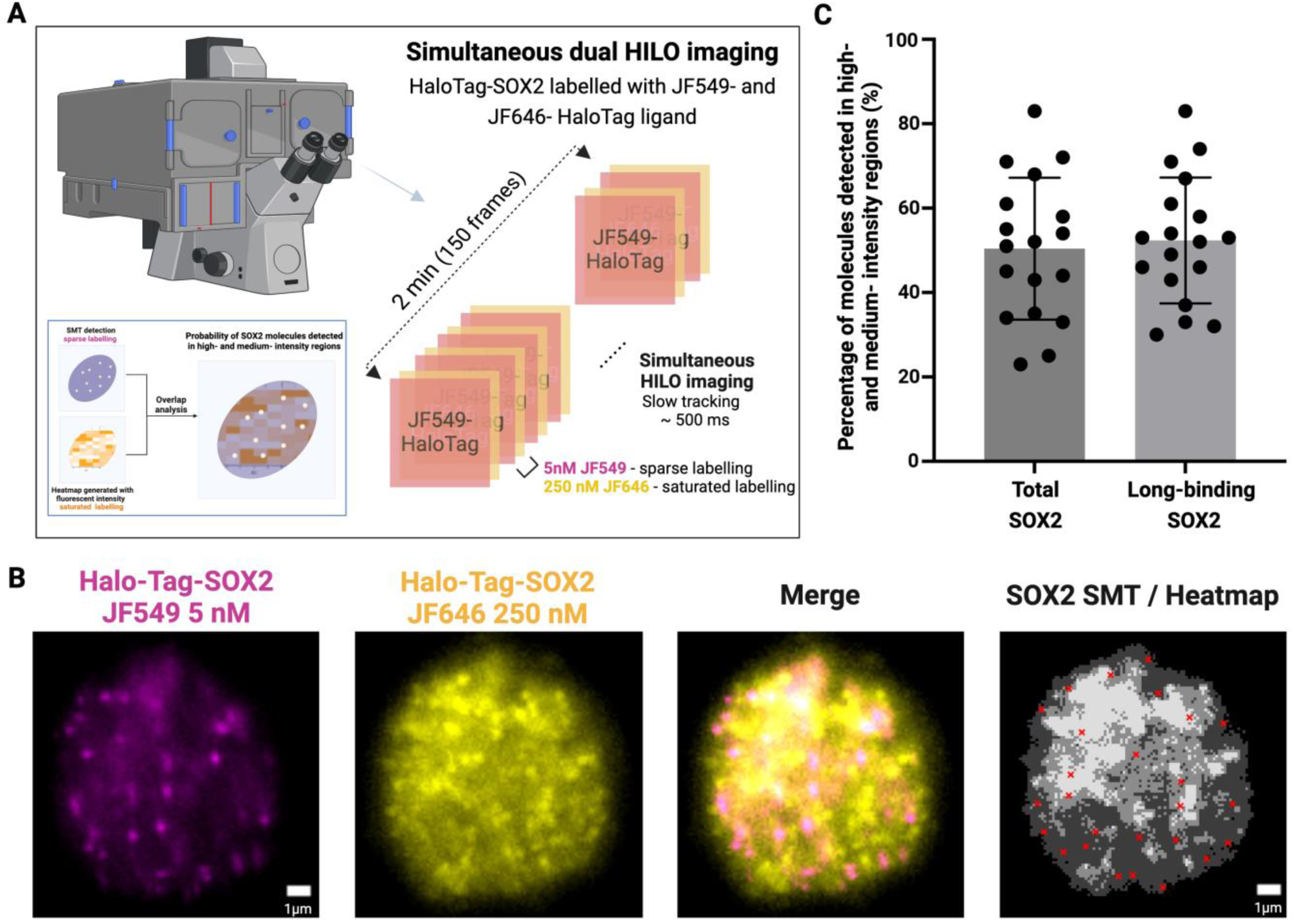
Simultaneous dual-colour HILO imaging to assess SOX2 clustering events. **(A)** Simultaneous SMT imaging of sparse labelling HaloTag-SOX2 (JF549, magenta) and saturated labelling HaloTag-SOX2 (JF646, yellow). The saturated signal is used to define a SOX2 heatmap of low, medium and high fluorescent intensity. This heatmap is then overlaid to SOX2 SMT signal to determine occupancy kinetics. **(B)** Fluorescent signals from HaloTag-SOX2 with sparse labelling (magenta) and saturated labelling (yellow). The merged image displays the combined signals from both channels. The greyscale image shows the SMT-SOX2 detected events overlaid onto the corresponding heatmap signal, with white representing high-intensity regions (top 8%), light grey as medium-intensity (top 30%), and dark grey as low-intensity (remaining 62%). **(C)** Percentage of total SOX2 and long-dwell time events detected by SMT (JF549) and overlapped with the high- and medium- intensity regions.

**Figure S2.**
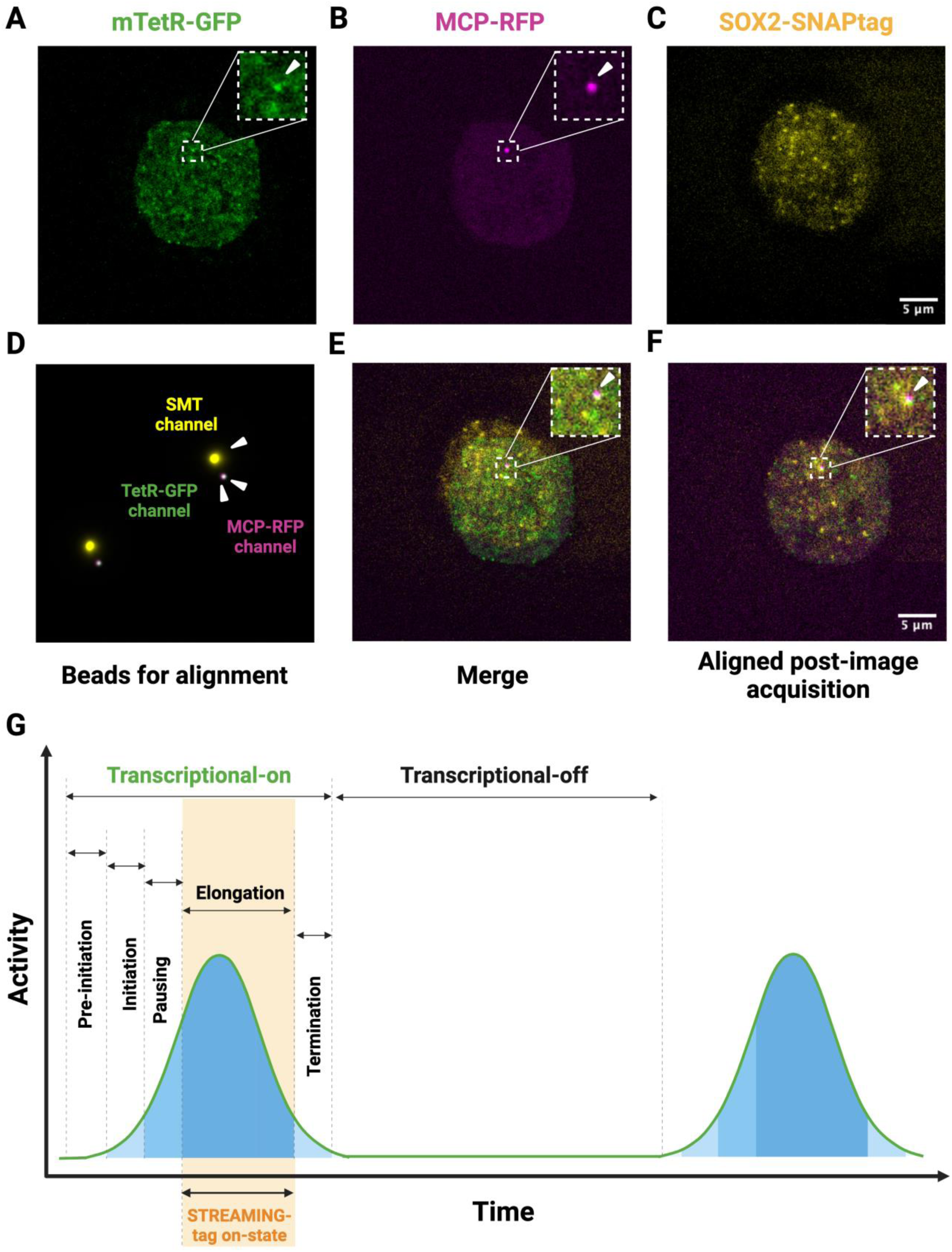
Observation of SOX2 occupancy at the *Nanog* locus in relation to its transcriptional state using a combination of SDC (STREAMING-tag) and HILO (SMT) microscopy, related to Figure 1. **(A-C)** Alignment of multimodal imaging for live cell imaging using a combination of SDC (STREAMING-tag) and HILO (SMT) microscopy. Simultaneous imaging of the *Nanog* locus (GFP, green), *Nanog* mRNA (RFP, magenta) and SOX2-SNAP SMT (yellow). White arrowheads indicate locations where MCP-RFP and mTetR-GFP spots were observed at the same time. **(D)** Fluorescent signal from the beads was obtained using the same setup in the GFP, RFP, and SMT channels. **(E)** Merged signal of GFP, RFP and SMT channels. **(F)** The signals from GFP, RFP, and SMT channels were merged, with spatial alignments achieved using fluorescent beads as a reference. **(G)** The STREAMING-tag reporter system (orange) is a faithful reporter of the elongation step.

**Figure S3.**
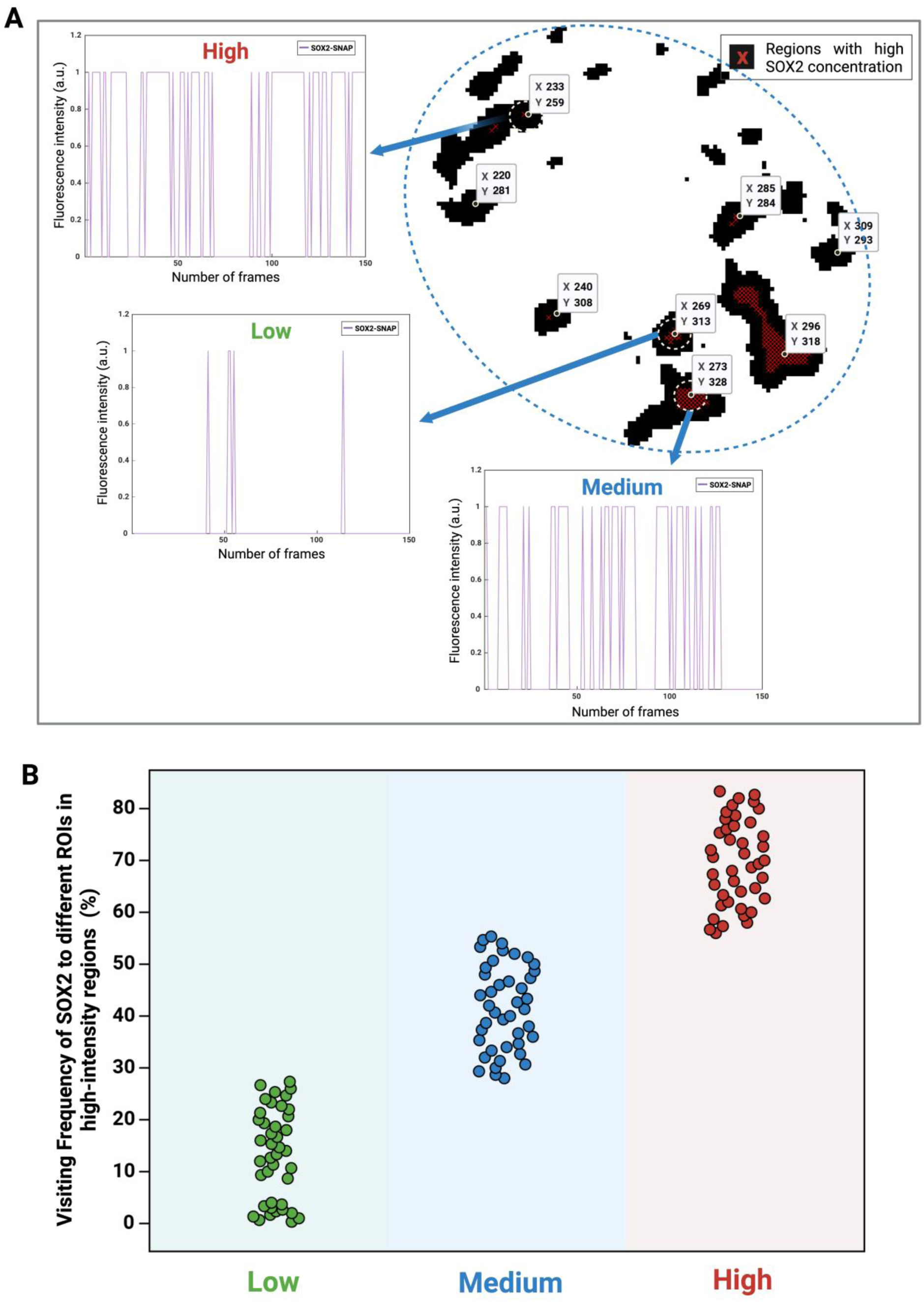
Dual-colour HILO imaging analysis to assess SOX2 clustering behaviours at high-intensity regions, related to Figure 2. **(A)** Representative images showing the frequency of SOX2 clusters (purple) tracked over a 2-minute window across different ROIs in high-intensity regions. The dashed circles indicate three representative ROIs with a radius of 4 pixels (440 nm). **(B)** Quantitation of different SOX2 behaviours classified based on the visiting frequency. Data represent n = 125 ROIs from 18 nuclei.

**Figure S4.**
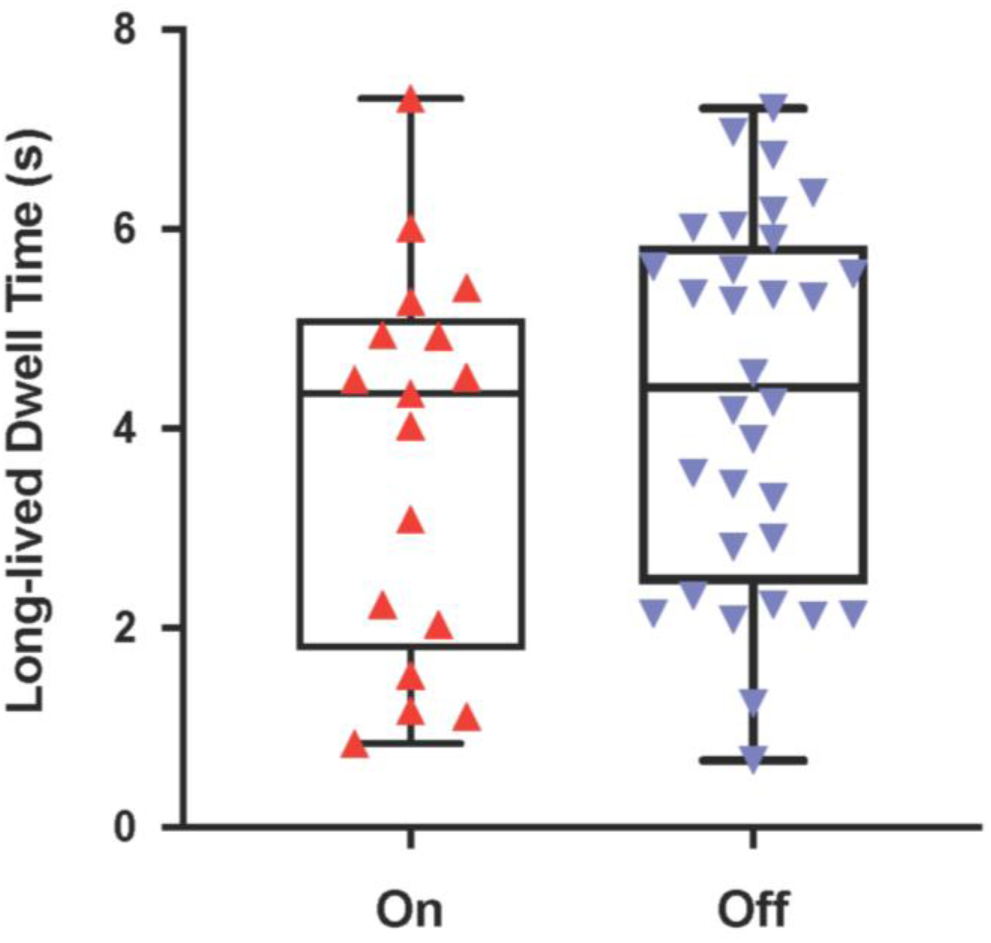
Quantification of SNAP-tagged SOX2 average population long-lived dwell time while the *Nanog* locus is active, related to Figure 3. Bar chart showing the mean value of SNAP-tagged SOX2 whole nucleus population long-lived dwell time during transcriptional on- and off-states of the *Nanog* locus.

**Figure S5.**
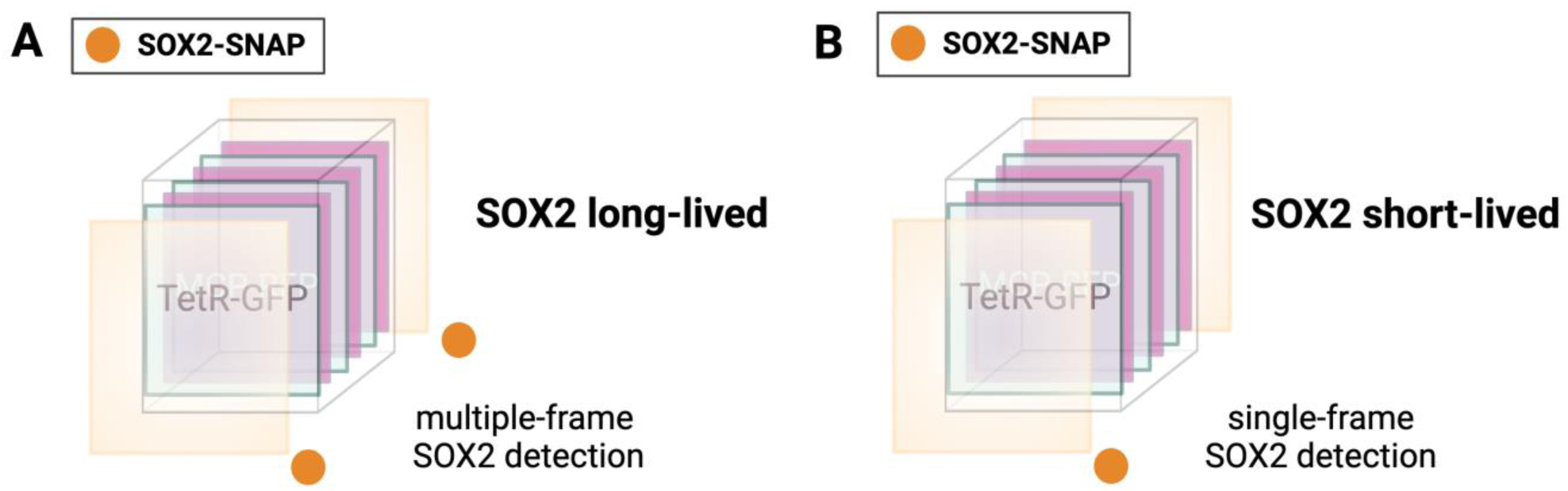
Co-detection of SOX2 clustering with *Nanog* mRNA elongation, related to Figure 3. **(A and B)** Schematic of SOX2 long-lived **(A**, 2 frames) and short-lived binding events **(B**, 1 frame).

**Figure S6.**
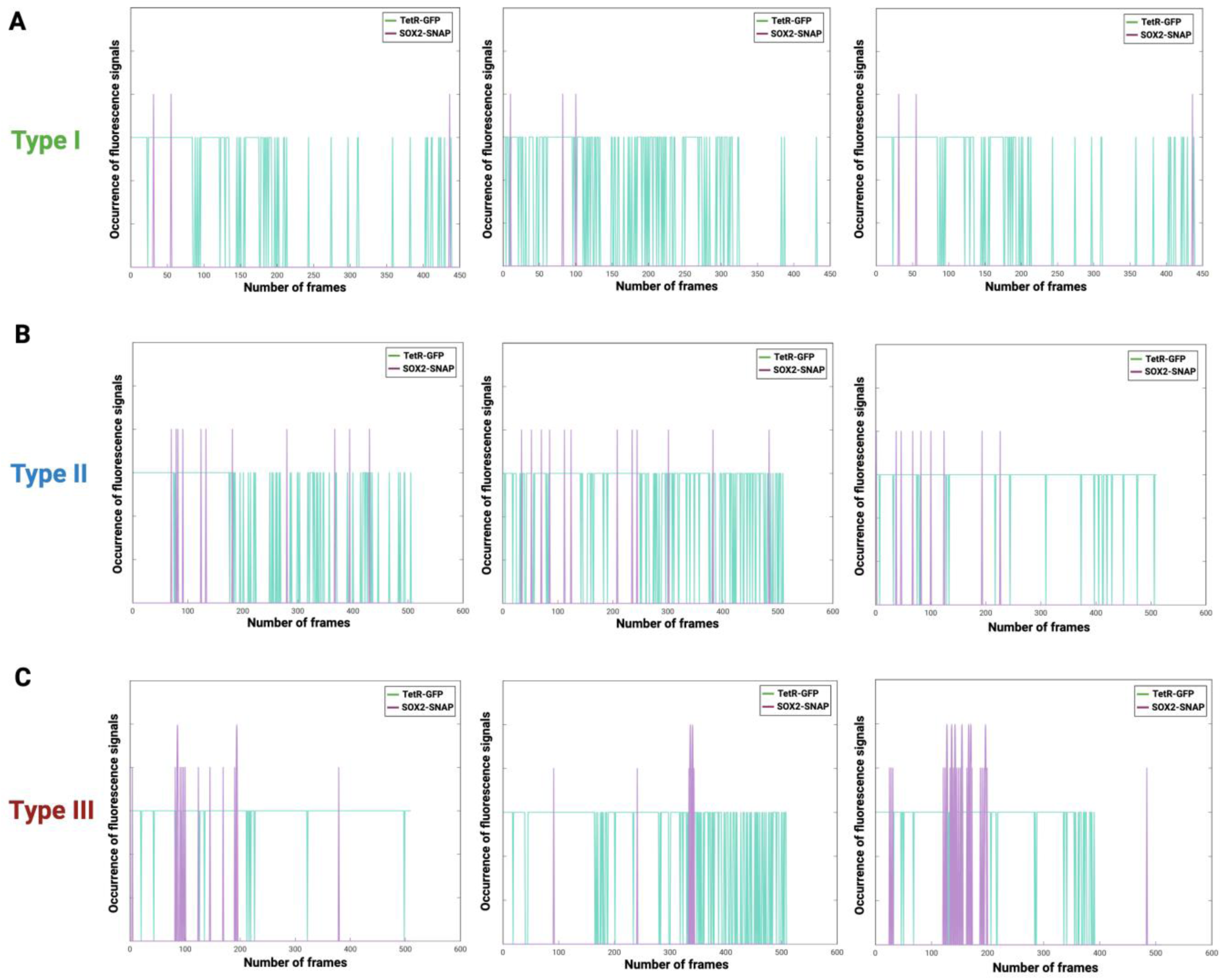
SOX2 visiting frequencies at the *Nanog* locus during off-state over 9 minutes (image acquisition mode 2), related to Figure 3. **(A-C)** Type I **(A),** Type II **(B)** and Type III **(C)** visiting frequencies represent SOX2 clustering dynamics (purple) during the off-state (GFP) within ROIs. Short peaks (purple) represent SOX2 short-lived binding events, while high peaks (purple) in **C** indicate SOX2 long-lived binding events. The visiting frequency accounts for the number of SOX2 detected molecules that have visited the GFP spot while *Nanog* transcriptional elongation is off.

**Figure S7.**
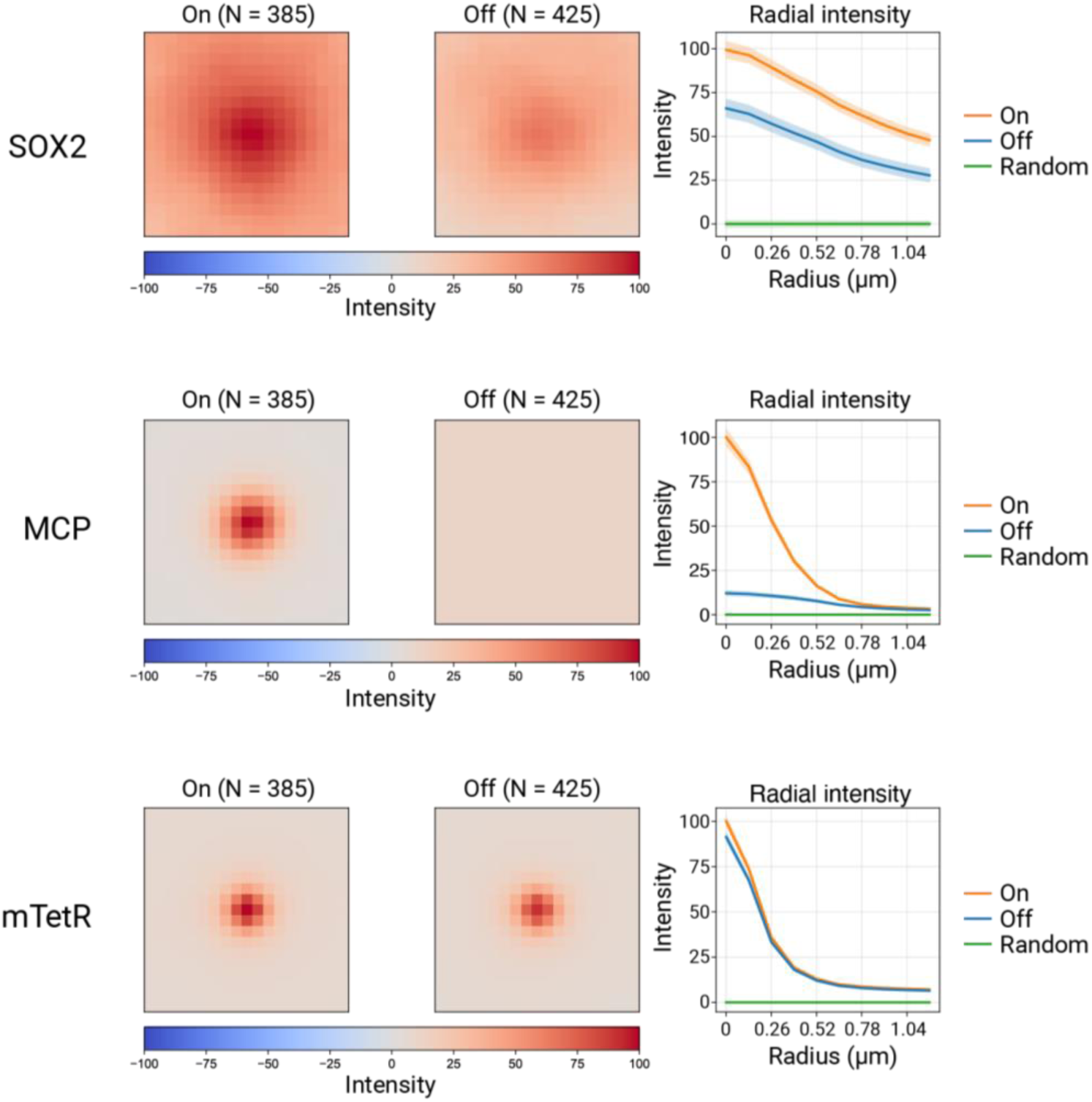
Live-cell imaging of SNAP-tagged SOX2 to quantify SOX2 accumulation at the *Nanog* locus during transcriptional “on” and “off” states, related to Figure 3. A *Nanog* STREAMING-tag knock-in cell line expressing mTetR-GFP (*Nanog* locus) and MCP-RFP (nascent *Nanog* RNA), together with endogenously SNAP-tagged SOX2, was imaged under live conditions. Images were extracted by centring analysis windows on the mTetR-GFP focus and mean-intensity projections were generated for each channel. SOX2 signal at the *Nanog* locus was consistently higher during transcriptionally “on” state compared with “off” state. Notably, although SOX2 intensity in the off state was lower than in the on state, it remained elevated relative to randomly sampled nuclear positions.

**Figure S8.**
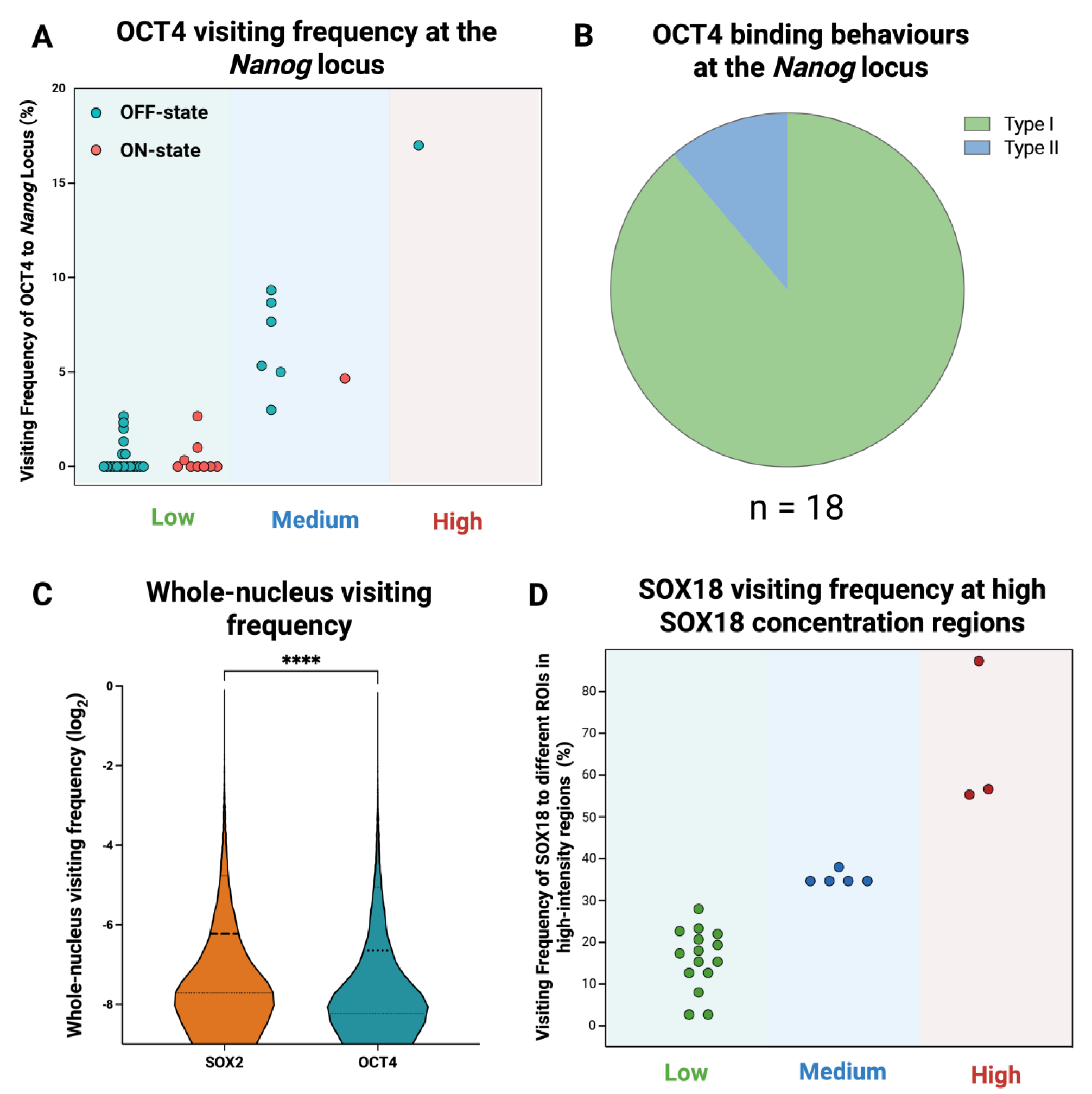
Comparison of SOX2, OCT4, and SOX18 visiting frequencies at the *Nanog* locus level and the genome-wide level, related to Figure 3. **(A)** Quantification of OCT4 visiting frequencies across transcriptional states during Mode 1 imaging acquisition. OCT4 behaviours were measured and classified for cells in the transcriptionally “on” state (n = 10) and “off” state (n = 26). **(B)** Pie charts showing the distribution of OCT4 binding behaviours (Type I, Type II, and absence of Type III events) during Mode 2 imaging acquisition. Data represent OCT4 dynamics in transcriptionally “on” (n = 14) and “off” (n = 4) states. **(C)** Violin plot comparing whole-nucleus visiting frequencies of SOX2 and OCT4. Data are log₂-transformed. Individual data points are overlaid, and the median (thick line) and interquartile range (thin lines) are indicated. **(D)** Dual-colour HILO imaging analysis of SOX18 to assess clustering behaviours at high-intensity regions. Visiting frequencies were quantified and classified using the same behavioural categories applied for SOX2 (Figure S3). Data represent n = 23 ROIs from 7 nuclei.

**Figure S9.**
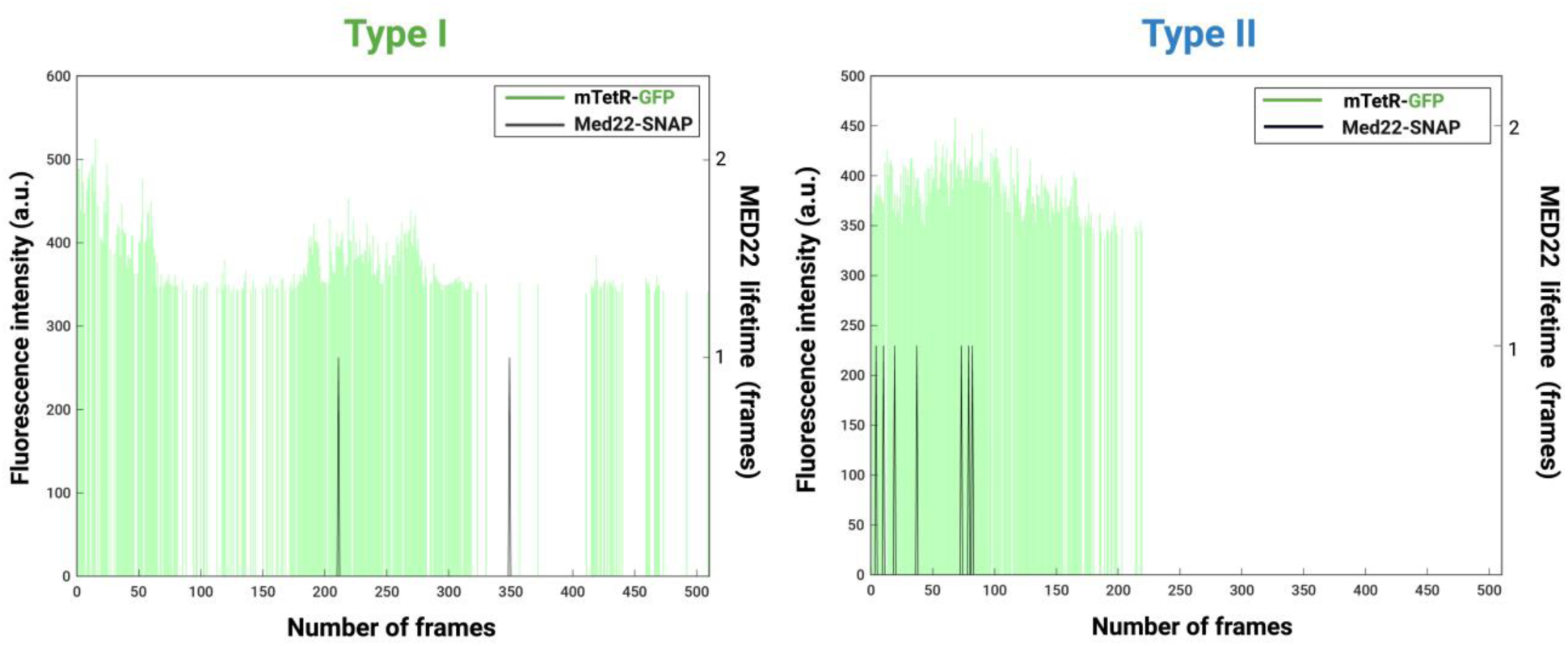
Two types of Med22 burst dynamics at the *Nanog* locus, related to Figure 3. Quantification of detected Med22 clusters and their relative survival rate at the *Nanog* locus. Fluorescent intensity at the *Nanog* locus is shown relative to time. *Nanog* locus signal is labelled with GFP (green) Med22 (black) short dwell times (over 1 frame) reveal Type I and II binding kinetics observed in the “off” state at the *Nanog* locus.

**Figure S10.**
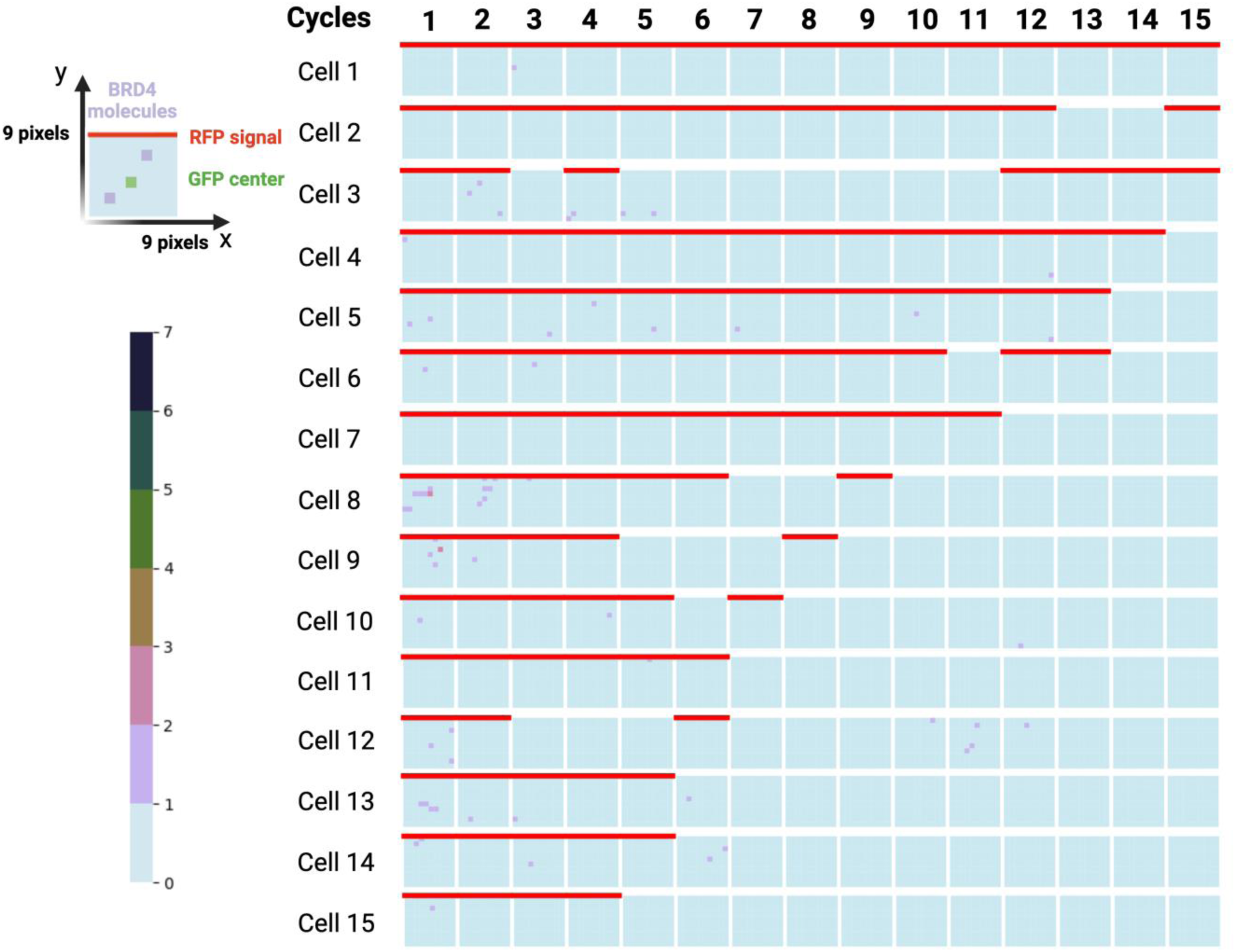
BRD4 clustering behaviour during the elongation process at the *Nanog* locus in live mESC, related to Figure 3. Each row represents an individual cell across a 9-minute time window (n = 15). Square plot showing BRD4 clustering behaviour during *Nanog* transcription on-state for 15 imaging cycles (9 min, 1 cycle = 10 frames) in live mESC. The heat map shows the number of BRD4 detected molecules. Each ROI (9 x 9 pixels, with a pixel size of 110 nm) has the GFP signal (*Nanog* locus) in the centre, the RFP signal (*Nanog* mRNA) is shown as a red line and the BRD4 SMT signal as a pink square. This plot summarises BRD4 visiting frequency during mRNA synthesis (on-state). Square plotting showing BRD4 clustering behaviour during *Nanog* transcription on-state for 10 imaging cycles (9 min) in mESC population (n = 15).

**Figure S11.**
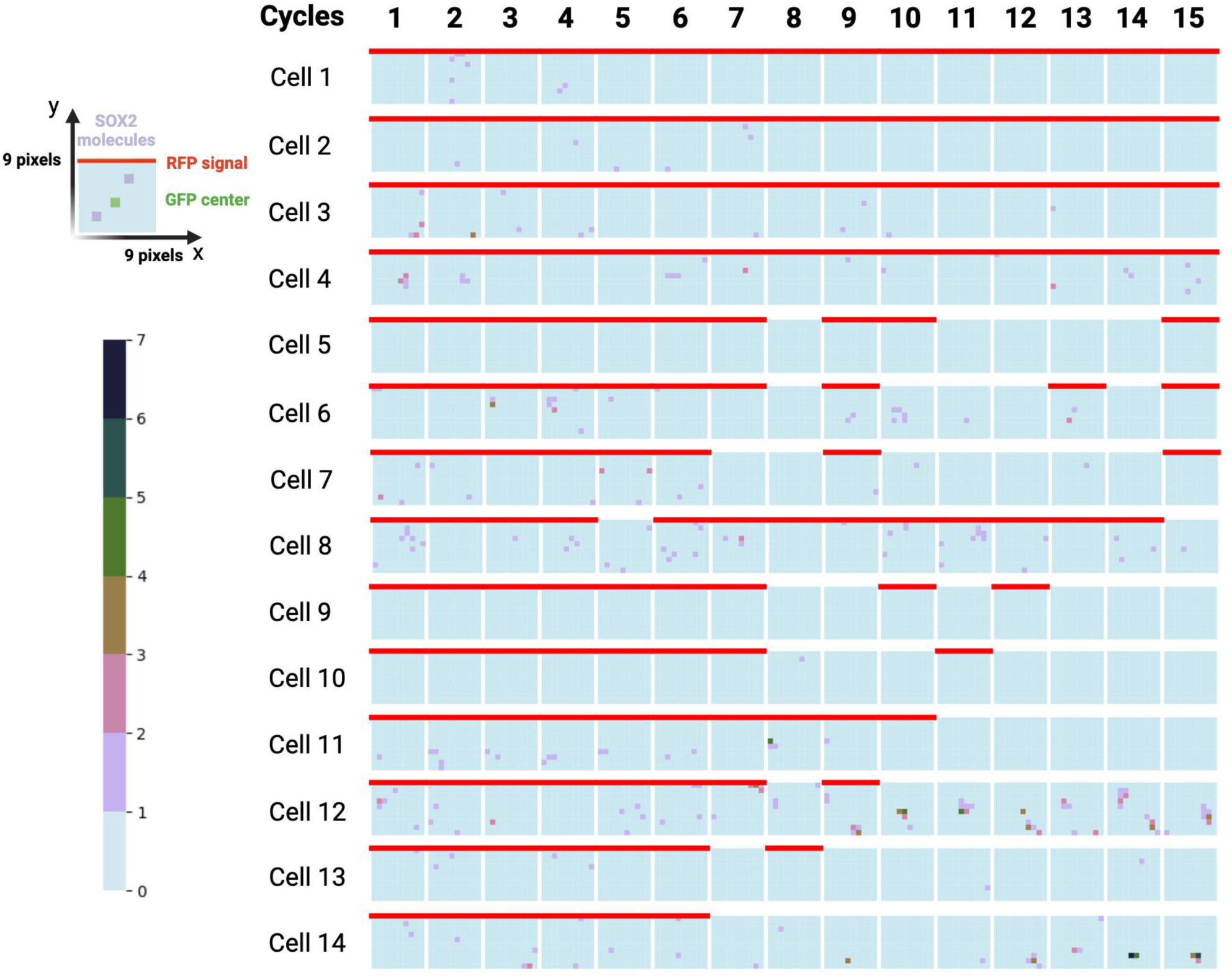
SOX2 clustering behaviour during elongation process at the *Nanog* locus in mESC population, related to Figure 3. Each row represents an individual cell across a 9-minute time window (n=14). Square plot showing SOX2 clustering dynamics during *Nanog* transcription on-state for 15 imaging cycles (9 min, 1 cycle = 10 frames) in live mESC. The heat map shows the number of SOX2 detected molecules. Each ROI (9 x 9 pixels, with a pixel size of 110 nm) has the GFP signal (*Nanog* locus) in the centre, the RFP signal (*Nanog* mRNA) is shown as a red line and the SOX2 SMT signal as a pink square.

**Figure S12.**
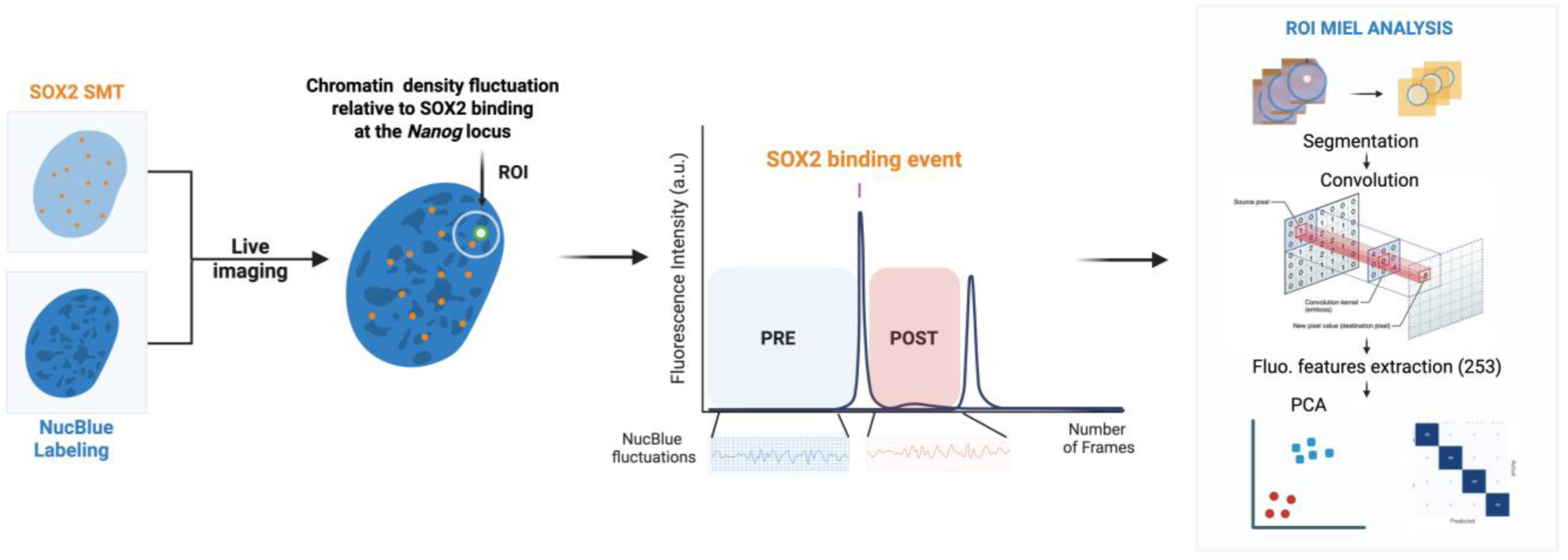
Workflow for ROI-based MIEL analysis of nuclear feature dynamics at the *Nanog* locus, related to Figure 3. SOX2 SMT and NucBlue^TM^ live-cell HILO imaging were performed during Mode 2 acquisition to monitor nuclear feature changes at the *Nanog* locus. SOX2 binding events were identified from SMT analysis, and the corresponding nuclear-feature traces within each ROI were aligned to the SOX2 binding frame to define pre- and post-binding intervals. Frame-wise NucBlue image patches within each ROI were extracted and subjected to temporal MIEL analysis. ROIs were segmented and convolved with a bank of texture filters to generate multiparametric fluorescence-feature vectors (253 features). PCA was then applied to quantify multivariate changes in chromatin morphology and texture associated with the transition from the pre- to the post-SOX2-binding state.

**Figure S13.**
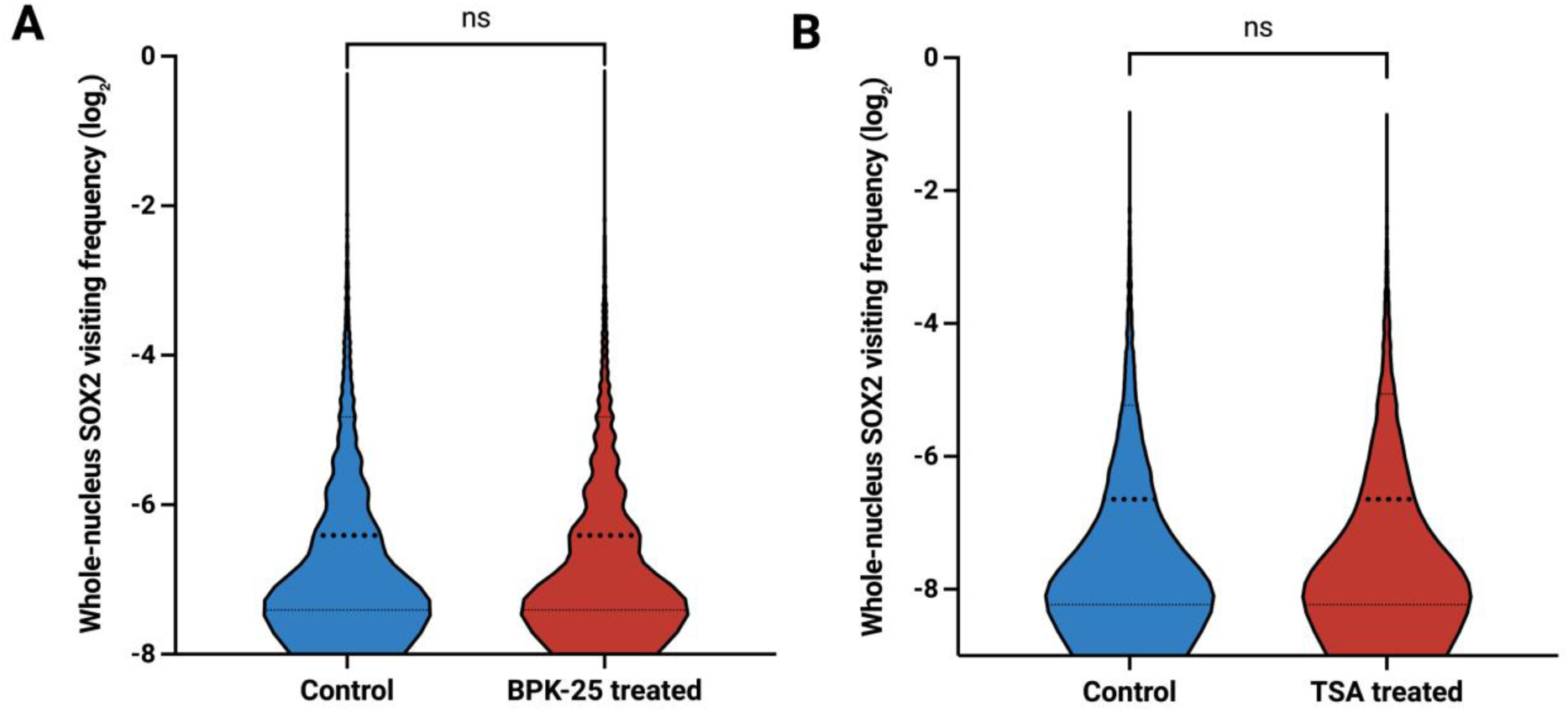
Violin plots showing the distribution of whole-nucleus SOX2 visiting frequency (log₂-transformed) in control versus inhibitor-treated cells, related to Figure 3. Individual datapoints are overlaid, with the median (thick dashed line) and interquartile range (thin dashed lines) indicated. **(A)** BPK-25 treatment does not significantly alter SOX2 visiting frequency compared with control. **(B)** TSA treatment similarly shows no significant change relative to control. “ns” indicates no statistically significant difference between groups (Mann-Whitney test).

**Figure S14.**
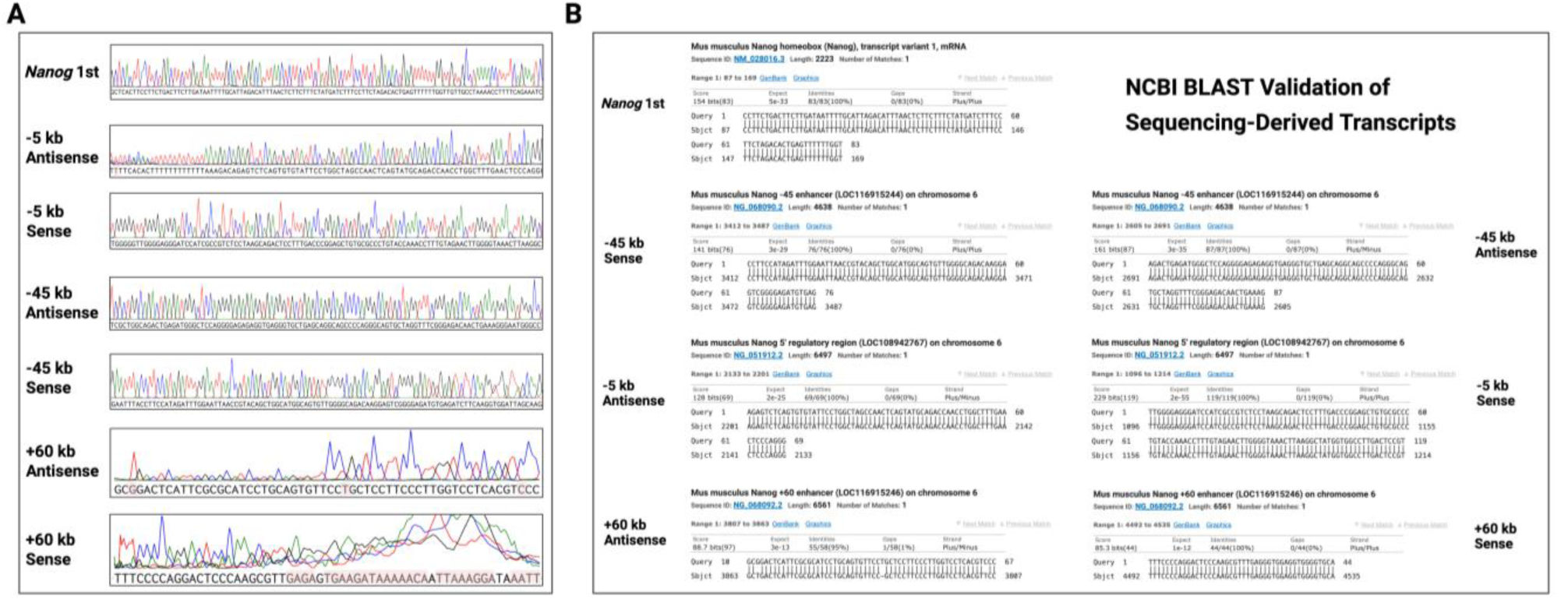
Validation of SOX2-associated *Nanog* nascent RNA and eRNA fragments by Sanger sequencing and NCBI BLAST alignment, related to Figure 4. **(A)** Sanger sequencing chromatograms of PCR-amplified RNA fragments isolated from the SOX2 pulldown, including the *Nanog* first exon and six eRNAs. Chromatograms display the corresponding regions of each PCR product, with the diagnostic mismatch base highlighted in red. **(B)** NCBI BLAST alignment results for each Sanger sequence, confirming that the recovered RNA fragments uniquely map to the correct genomic loci, including the *Nanog* gene body and the respective enhancer regions.

**Figure S15.**
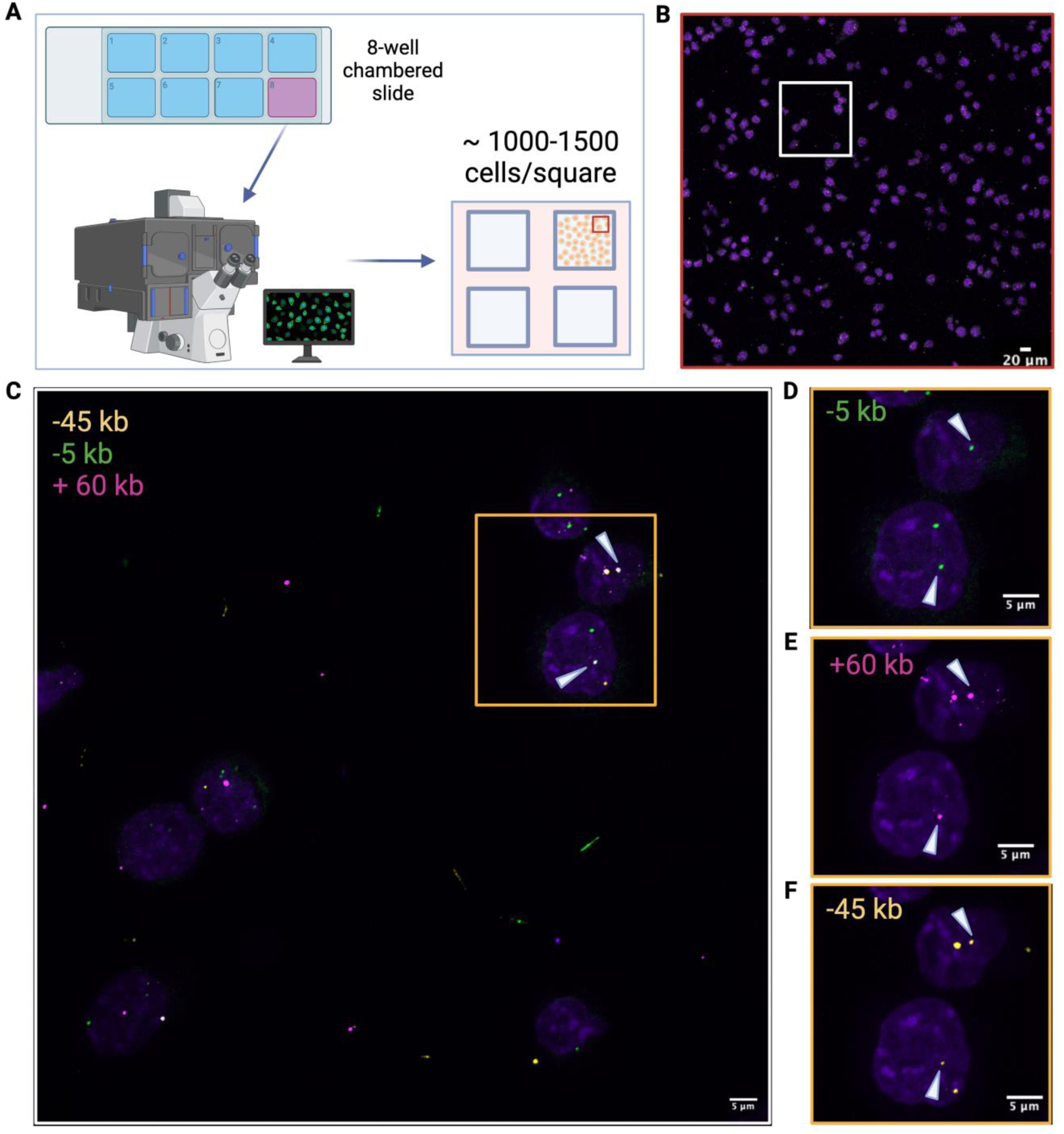
Sample images used for the quantification of *Nanog* sense eRNAs co-expression and co-localisation in mESCs population, related to Figure 5. **(A)** Schematic of the large-scale imaging acquisition setup to capture four regions of signal from one 8-well chambered slide (around 4000-6000 cells imaged per condition). **(B)** Microscope image showing the fluorescent signal captured from a portion of the whole cell population (circled in red in suppl fig 10A). **(C)** Zoomed-in image with the fluorescent signal of -45 kb (yellow), -5 kb (green) and +60 kb (magenta) eRNAs. White arrowheads indicate locations where all 3 eRNAs were observed colocalised. **(D-F)** Zoomed-in images with the fluorescent signal of 3 eRNAs respectively, **(D)** -5 kb (green), **(E)** +60 kb (magenta) and **(F)** -45 kb (yellow), scale bar 5μm.

**Figure S16.**
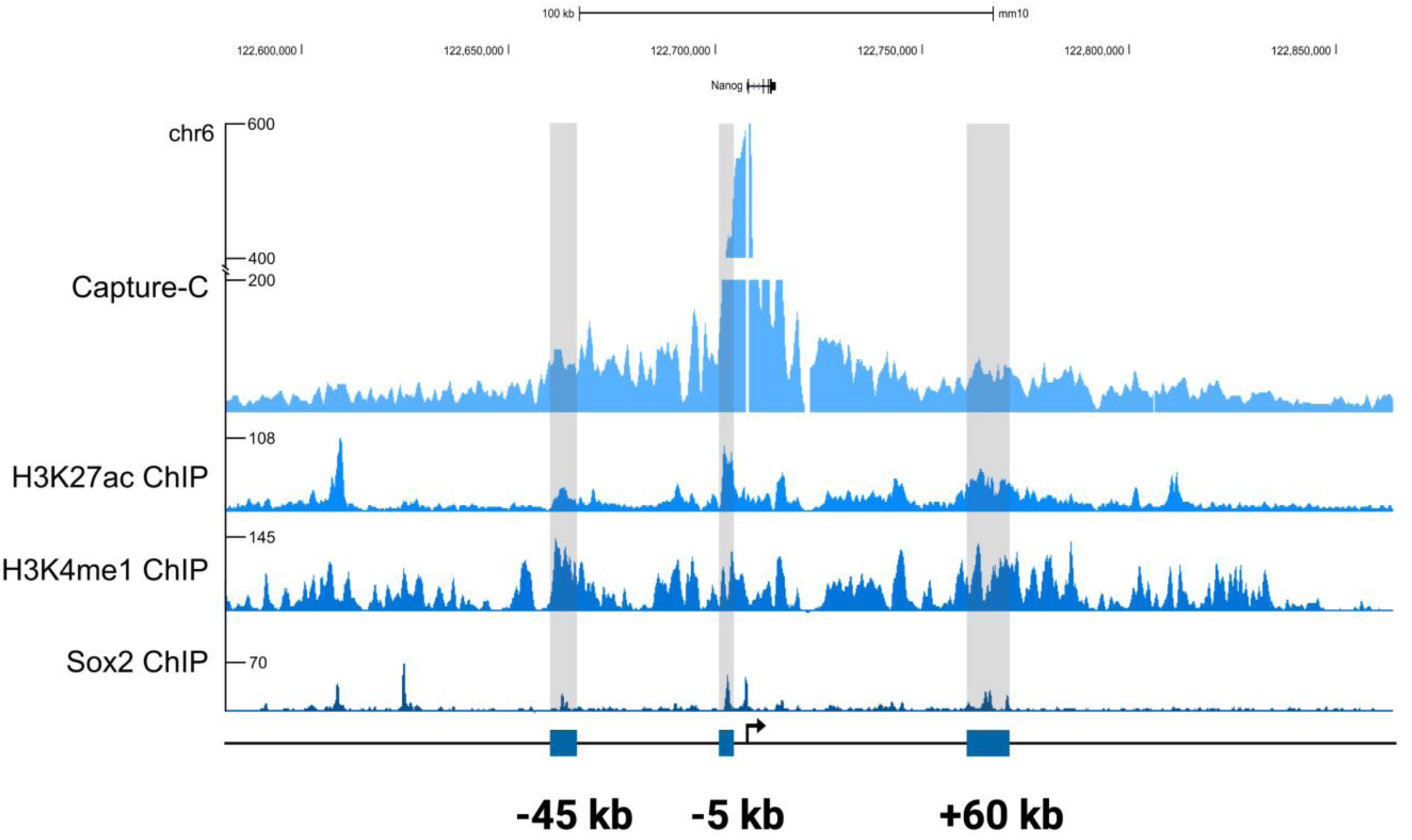
The *Nanog* promoter interacts with three enhancer regions, related to Figure 5. Contact points between *Nanog* promoter and enhancer regions (significant interactions when score >= 5) using Capture-C analysis. The highlighted regions (grey) correspond to -45 kb, -5 kb and +60 kb *Nanog* enhancers.

**Figure S17.**
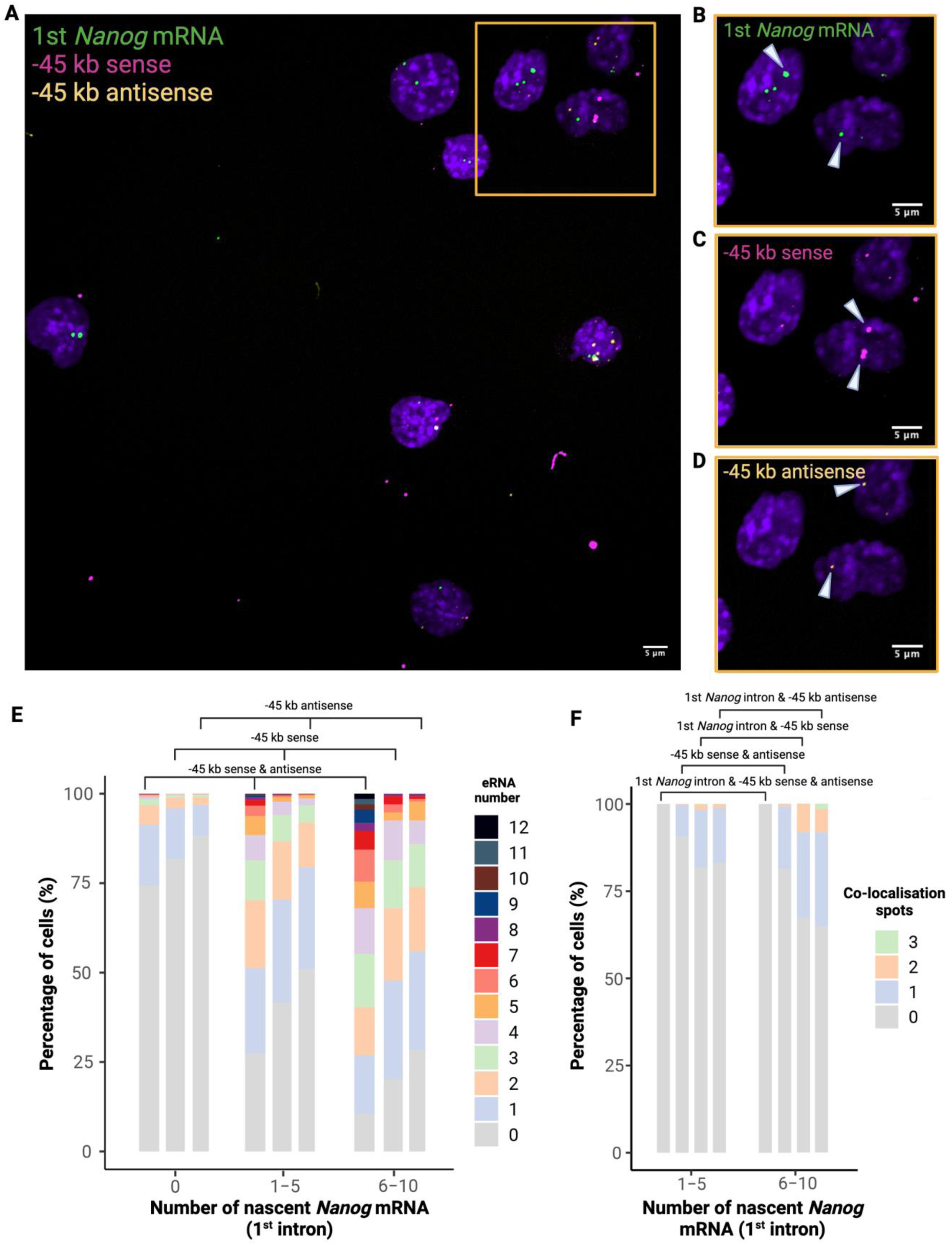
Quantification of *Nanog* -45 kb eRNAs expression relative to *Nanog* mRNA first intron synthesis in mESCs population, related to Figure 5. **(A)** Microscope image showing the fluorescent signal of first intron *Nanog* mRNA (green), *Nanog* -45 kb sense (magenta) and antisense (yellow) eRNAs captured from mESCs. **(B-D)** Zoomed-in images (orange inset) showing individual fluorescent signal of first intron *Nanog* mRNA and eRNAs respectively. White arrowheads point to where eRNA spots were observed. **(E)** Quantitation of *Nanog* -45 kb sense and antisense eRNAs co-expression with first intron *Nanog* mRNA shows a positive correlation between enhancer and gene body transcription. **(F)** Quantitation of *Nanog* -45 kb sense and antisense eRNAs co-localisation with first intron *Nanog* mRNA suggests that enhancers are lowly transcribed while *Nanog* mRNA is produced.

**Figure S18.**
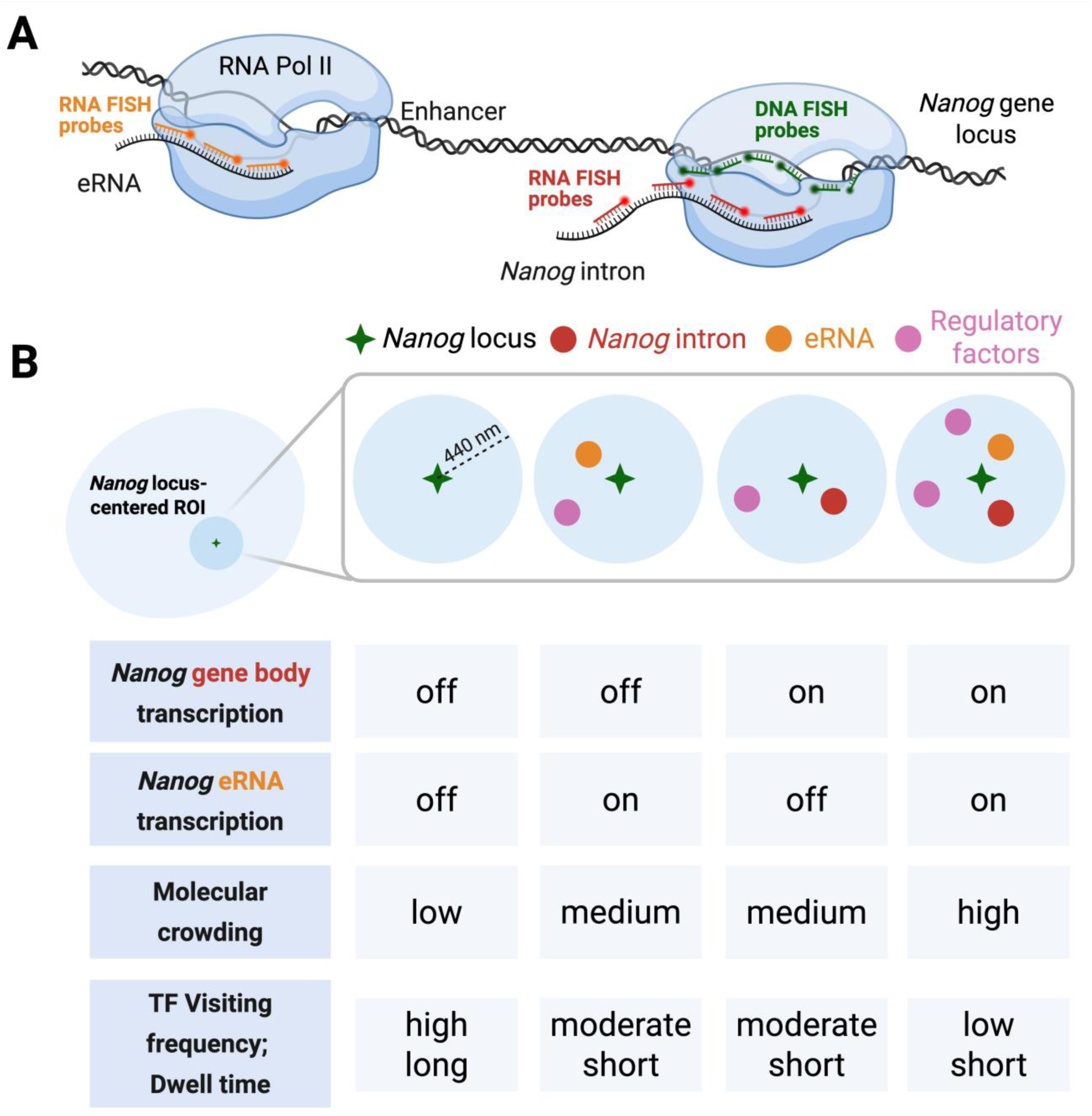
Schematic representation of sequential RNA-FISH and DNA-FISH, related to Figure 5. Sequential RNA-DNA FISH was performed to visualise the spatial relationship between *Nanog* genomic loci (green), *Nanog* introns (red), and *Nanog* eRNAs (orange) within the same cells. Quantification of *Nanog* eRNAs and intron mRNA co-localisation at the *Nanog* locus reveals the temporal dynamics of individual eRNA transcription in relation to *Nanog* gene body transcription.

**Figure S19.**
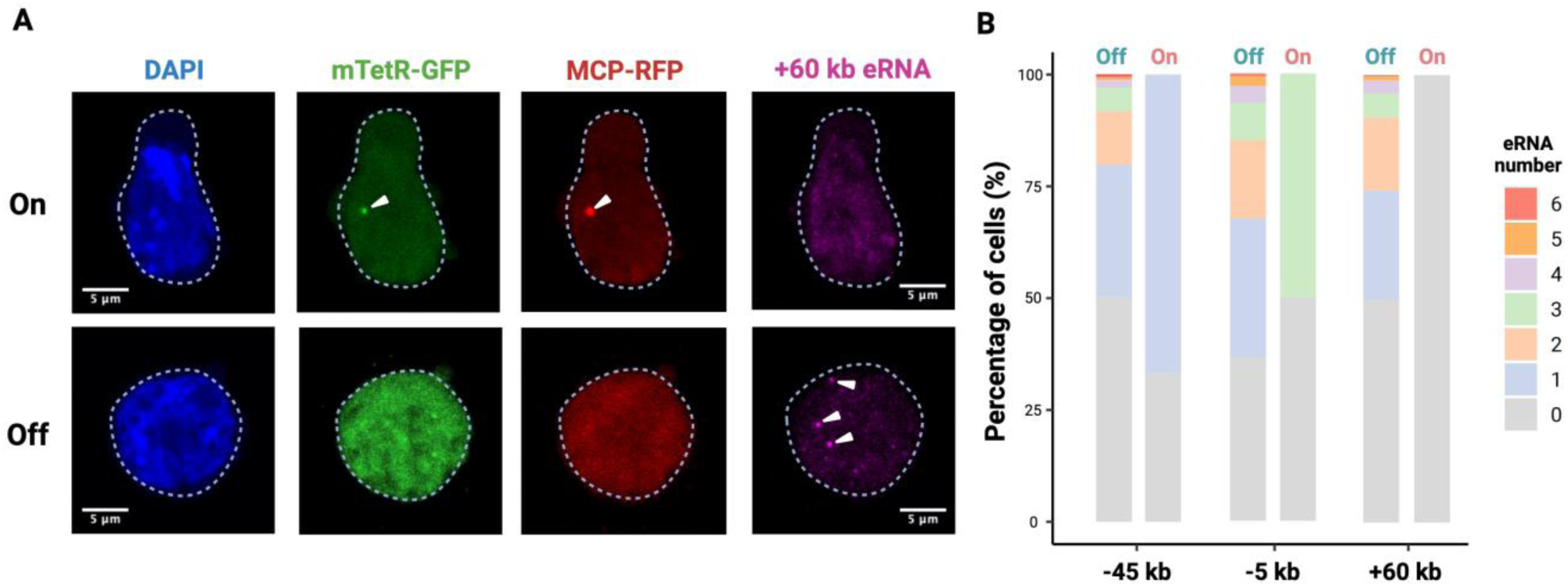
Quantification of *Nanog* eRNAs expression relative to *Nanog* mRNA synthesis using RNA-FISH and STREAMING-tag reporter system, related to Figure 5. **(A)** Single-molecule FISH for the +60 kb eRNA combined with STREAMING tag imaging shows that the +60 kb eRNA is expressed while *Nanog* mRNA nascent transcription is off (green *Nanog* locus, red *Nanog* mRNA, magenta +60 kb eRNA, blue DAPI). **(B)** Temporal relationship between individual eRNAs from three different *Nanog* enhancers relative to the co-expression of the *Nanog* STREAMING-tag reporter system.

**Video S1.**
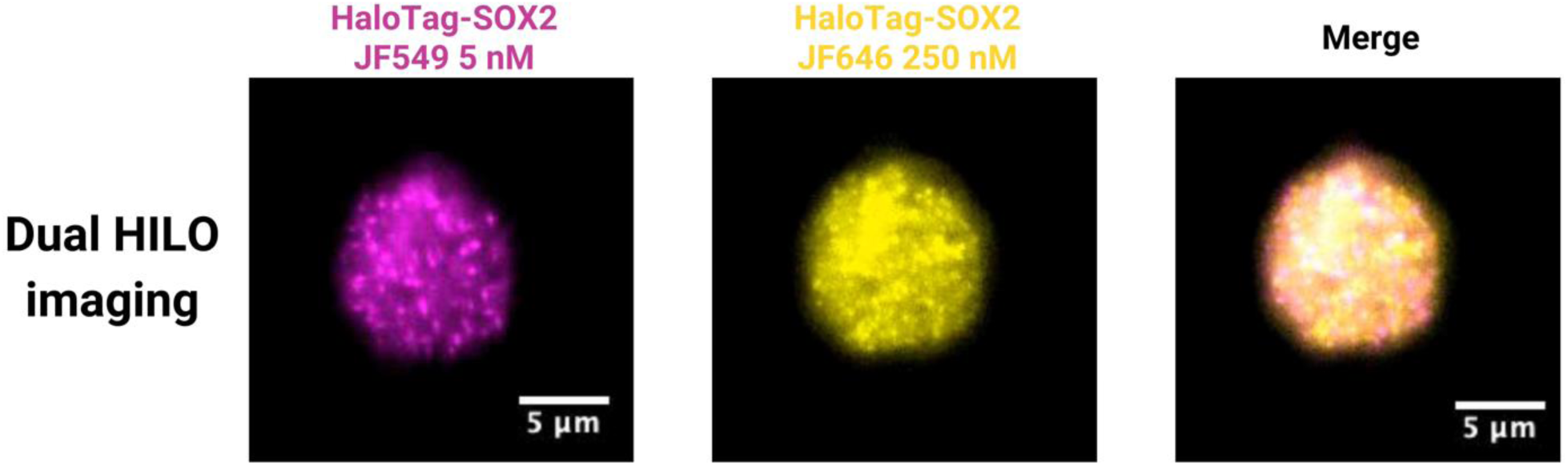
Simultaneous dual-channel imaging for SOX2 clustering events, related to Figure 1. Fluorescent signals from the sparse labelling HaloTag-SOX2 (magenta) and saturated labelling HaloTag-SOX2 (yellow). Merged video displaying the signal from both channels.

**Video S2.**
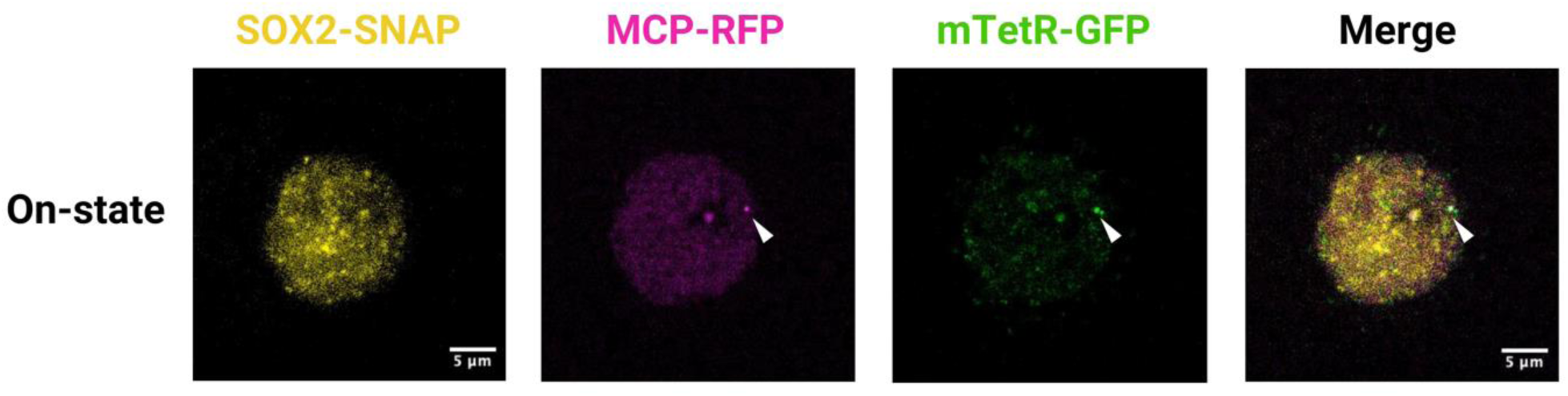
Multimodal live imaging of on-state cells using mode 1 image acquisition method, related to Figure 2. Fluorescent signals from the *Nanog* locus (GFP, green), *Nanog* mRNA (RFP, magenta) and SOX2-SNAPtag SMT (yellow). Merged video displaying the signal from all channels with the spatial alignment. White arrowheads indicate locations where MCP-RFP and mTetR-GFP spots were observed simultaneously.

**Video 3.**
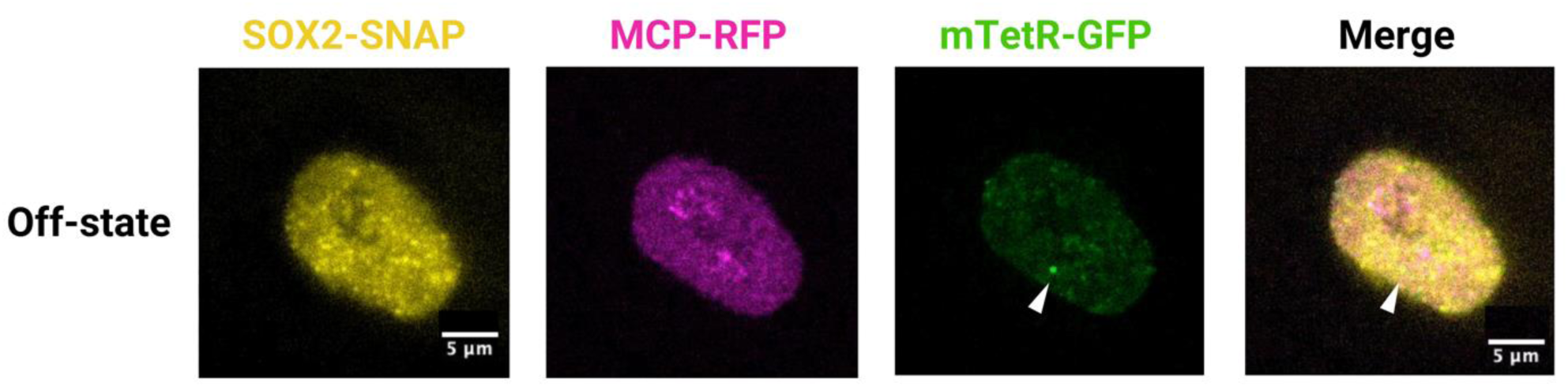
Multimodal live imaging of off-state cells (low SOX2 visiting frequency) using image acquisition mode 1, related to Figure 2. Fluorescent signals from the *Nanog* locus (GFP, green), *Nanog* mRNA (RFP, magenta) and SOX2-SNAP SMT (yellow). Merged video displaying the signal from all channels with the spatial alignment. White arrowheads point to the location where the mTetR-GFP spot was observed.

**Video 4.**
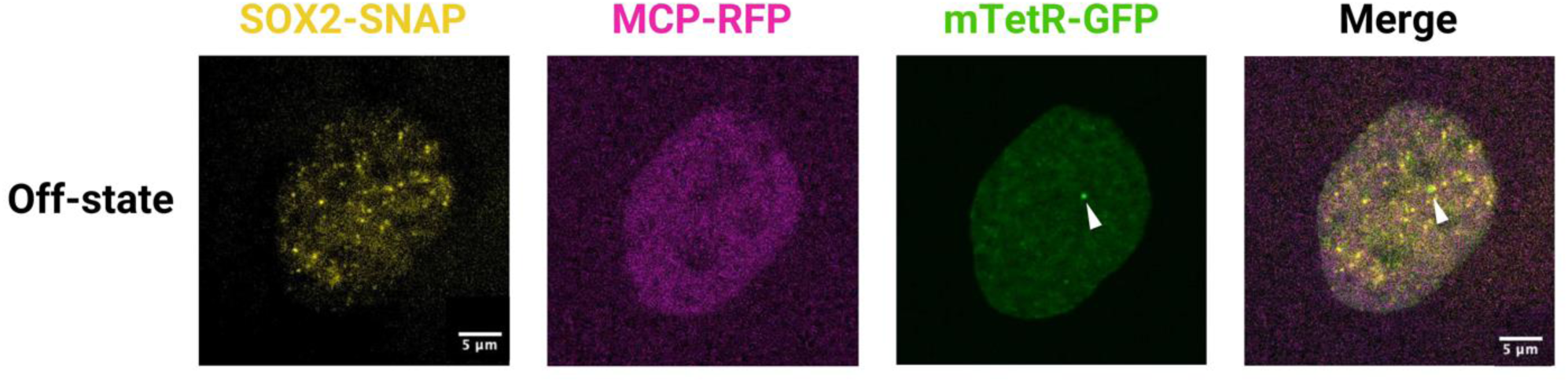
Multimodal live imaging of off-state cells (moderate SOX2 visiting frequency) using image acquisition mode 1, related to Figure 2. Fluorescent signals from the *Nanog* locus (GFP, green), *Nanog* mRNA (RFP, magenta) and SOX2-SNAP SMT (yellow). Merged video displaying the signal from all channels with the spatial alignment. White arrowheads point to the location where the mTetR-GFP spot was observed.

**Video 5.**
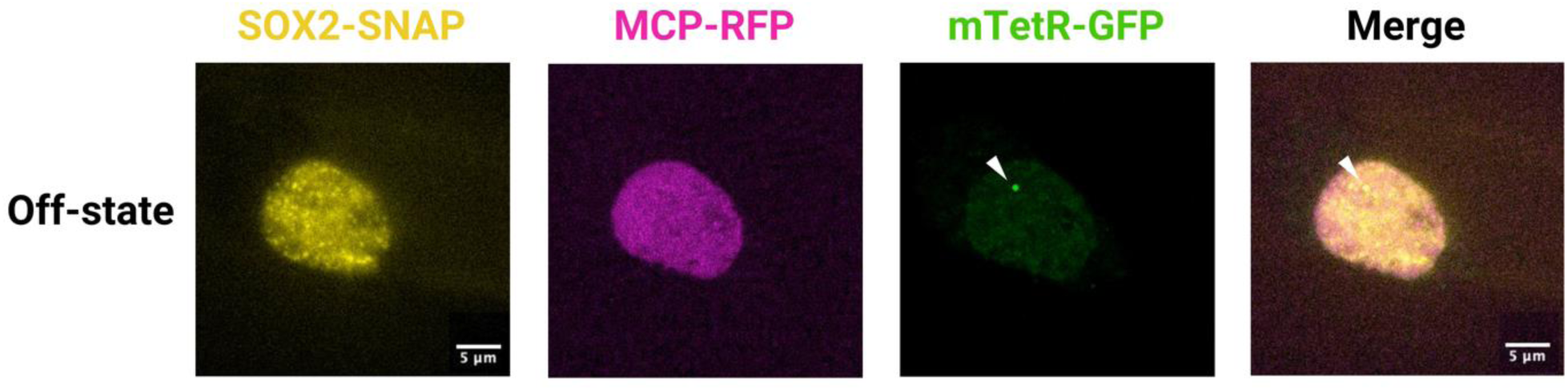
Multimodal live imaging of off-state cells (high SOX2 visiting frequency) using image acquisition mode 1, related to Figure 2. Fluorescent signals from the *Nanog* locus (GFP, green), *Nanog* mRNA (RFP, magenta) and SOX2-SNAPtag SMT (yellow). Merged video displaying the signal from all channels with the spatial alignment. White arrowheads point to the location where the mTetR-GFP spot was observed.

**Video 6.**
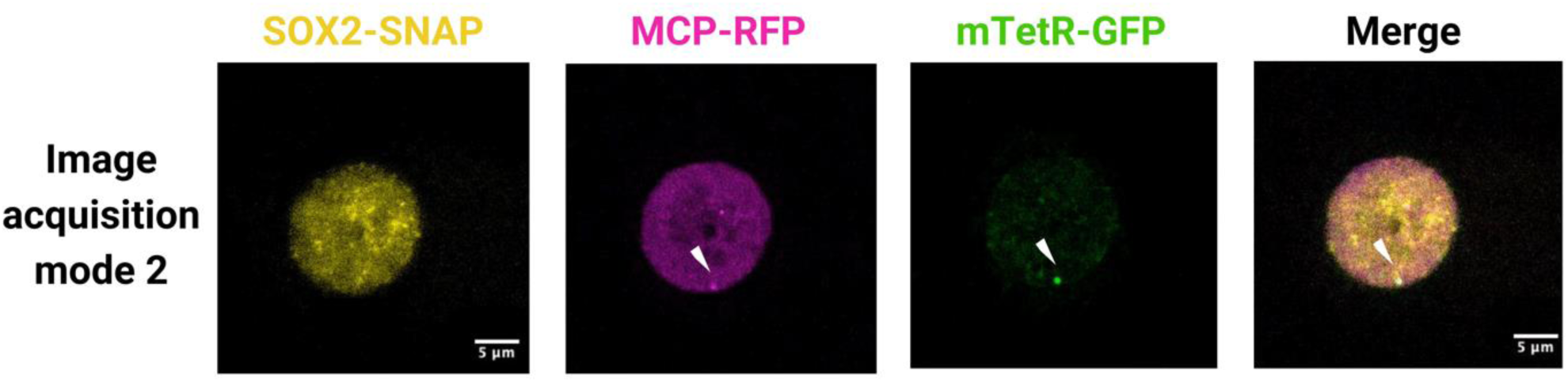
Multimodal live imaging of cells using mode 2 image acquisition method. This type of imaging acquisition contains both an on-state and an off-state, related to Figure 3. Fluorescent signals from the *Nanog* locus (GFP, green), *Nanog* mRNA (RFP, magenta) and SOX2-SNAPtag SMT (yellow). Merged video displaying the signal from all channels with the spatial alignment. White arrowheads indicate locations where MCP-RFP and mTetR-GFP spots were observed simultaneously.

**Table S1.**
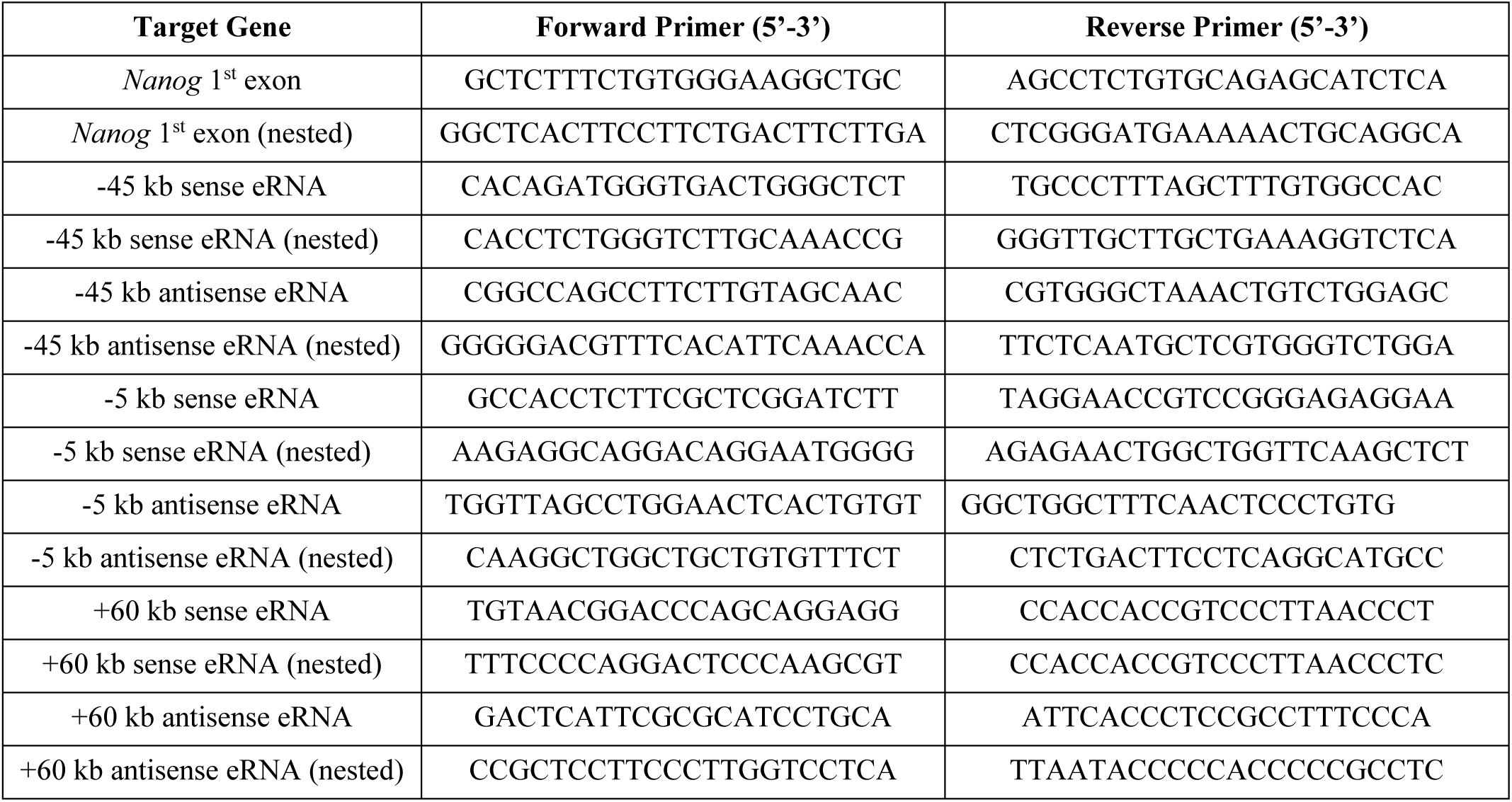
Primer sequences used for verification of SOX2–RNA interactions. List of primers used to amplify *Nanog* first exon and eRNA fragments recovered from SOX2 pulldown samples.

## Data and materials availability

All data are available in the main text or the supplementary materials and have been deposited to Datadryad for access to raw files.

Accession number and link: https://doi.org/10.5061/dryad.wpzgmsbw4

